# An Enzymatic TMPRSS2 Assay for Assessment of Clinical Candidates and Discovery of Inhibitors as Potential Treatment of COVID-19

**DOI:** 10.1101/2020.06.23.167544

**Authors:** Jonathan H. Shrimp, Stephen C. Kales, Philip E. Sanderson, Anton Simeonov, Min Shen, Matthew D. Hall

## Abstract

SARS-CoV-2 is the viral pathogen causing the COVID19 global pandemic. Consequently, much research has gone into the development of pre-clinical assays for the discovery of new or repurposing of FDA-approved therapies. Preventing viral entry into a host cell would be an effective antiviral strategy. One mechanism for SARS-CoV-2 entry occurs when the spike protein on the surface of SARS-CoV-2 binds to an ACE2 receptor followed by cleavage at two cut sites (“priming”) that causes a conformational change allowing for viral and host membrane fusion. TMPRSS2 has an extracellular protease domain capable of cleaving the spike protein to initiate membrane fusion. A validated inhibitor of TMPRSS2 protease activity would be a valuable tool for studying the impact TMPRSS2 has in viral entry and potentially be an effective antiviral therapeutic. To enable inhibitor discovery and profiling of FDA-approved therapeutics, we describe an assay for the biochemical screening of recombinant TMPRSS2 suitable for high throughput application. We demonstrate effectiveness to quantify inhibition down to subnanomolar concentrations by assessing the inhibition of camostat, nafamostat and gabexate, clinically approved agents in Japan. Also, we profiled a camostat metabolite, FOY-251, and bromhexine hydrochloride, an FDA-approved mucolytic cough suppressant. The rank order potency for the compounds tested are: nafamostat (IC_50_ = 0.27 nM), camostat (IC_50_ = 6.2 nM), FOY-251 (IC_50_ = 33.3 nM) and gabexate (IC_50_ = 130 nM). Bromhexine hydrochloride showed no inhibition of TMPRSS2. Further profiling of camostat, nafamostat and gabexate against a panel of recombinant proteases provides insight into selectivity and potency.

## Introduction

The severe acute respiratory syndrome-related coronavirus 2 (SARS-CoV-2) pandemic has driven the urgent need to rapidly identify therapeutics for both preventing and treating infected patients. Given that no approved therapeutics for treating any coronaviruses existed at the time SARS-CoV-2 emerged (late 2019), early attention has focused on drug repurposing opportunities^1, 2^. Drug repurposing is an attractive approach to treating SARS-CoV-2, as active drugs approved for use in humans in the United States or by other regulatory agencies, or unapproved drug candidates shown to be safe in human clinical trials, can be nominated for fast-track to the clinic. For example, remdesivir (GS-5734, Gilead Sciences Inc.), is an inhibitor of viral RNA-dependent RNA polymerase that had previously been in clinical trials for treating Ebola virus. Remdesivir was rapidly shown to be active against SARS-CoV-2 *in vitro* and in clinical trials, which resulted in the FDA granting emergency use authorization and full approval in Japan^3^. The delineation of targets and cellular processes that mediate SARS-CoV-2 infection and replication forms the basis for the development of assays for drug repurposing screening and subsequent full-fledged therapeutic development programs.

One therapeutic target receiving significant attention is the human host cell transmembrane protease serine 2 (TMPRSS2, Uniprot – O15393^4^) that is expressed in epithelial cells of the human respiratory and gastrointestinal tracts^5^. TMPRSS2 is anchored to the extracellular surface of the cell, where it exerts its enzymatic activity. While its precise physiologic substrate is not clear, TMPRSS2 gene fusions are common in prostate cancer, resulting in its overexpression^6, 7^. The SARS-CoV-2 virus enters cells *via* its spike protein first binding to the cell-surface angiotensin-converting enzyme 2 (ACE2), and evidence suggests that TMPRSS2 then proteolytically cleaves a sequence on the spike protein, facilitating a conformation change that ‘primes’ it for cell entry (Figure 1A). TMPRSS2 was first shown to facilitate viral entry of the coronaviruses SARS-CoV and HCoV-NL63 in cells engineered to overexpress TMPRSS2, and by inhibition with the trypsin-like serine protease inhibitor, camostat^8^. When the Middle East respiratory syndrome-related coronavirus (MERS-CoV) outbreak occurred, TMPRSS2-overexpressing cells were again shown to facilitate cell infection, TMPRSS2 was shown to degrade the MERS-CoV spike protein, and camostat was shown to limit cell entry^9^. The structurally related trypsin-like serine protease inhibitor nafamostat was shown to similarly inhibit spike protein-mediated cell fusion of MERS-CoV^10^. Additionally, *in vivo* mouse models for SARS-CoV and MERS-CoV with a TMPRSS2 gene knockout caused a reduction in lung pathology after viral infection while not seeing any effect on development or survival. However, it was noted that a decreased inflammatory chemokine and cytokine response mediated by a Toll-like receptor 3 agonist may suggest an unidentified physiological role^11^. The protein sequence between mouse and human is conserved, with 77% sequence identity suggesting structure and functional similarity^12^. Given the strong evidence that TMPRSS2 mediates coronavirus entry, when SARS-CoV-2 emerged it was soon demonstrated through loss- and gain-of-function experiments that TMPRSS2 is retained as a mediator of cell infection, and that this can be inhibited by camostat^13-17^.

**Figure 1:**
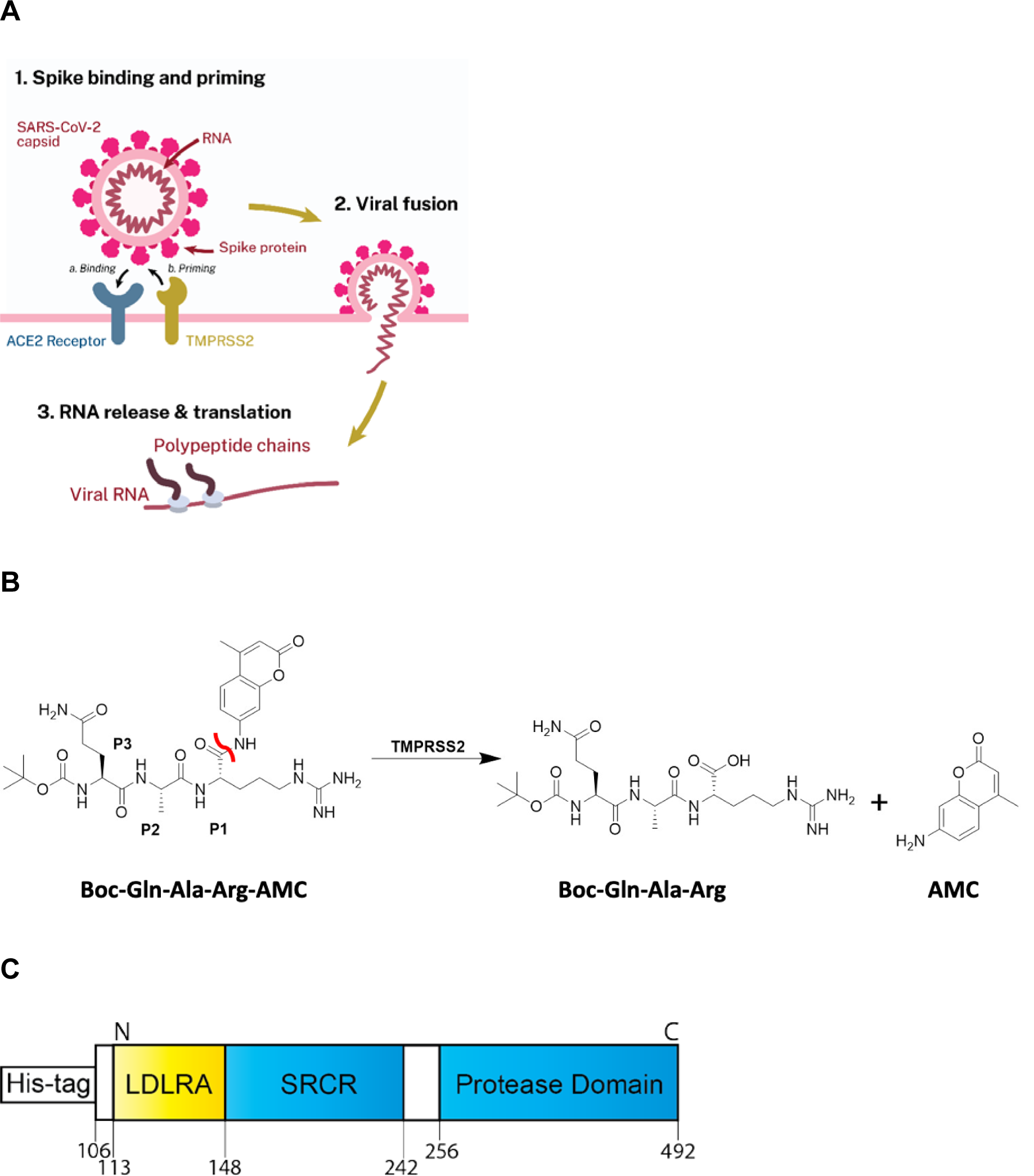
**A)** Scheme demonstrating the role TMPRSS2 plays in priming SARS-CoV-2 for cellular entry. Spike protein first binds to ACE2 (‘Binding’), followed by proteolytic action of TMPRSS2 (‘Priming’) prior to viral fusion. **B)** Scheme displaying the enzymatic assay principle. The fluorogenic peptide substrate Boc-Gln-Ala-Arg-AMC has low fluorescence compared to the fluorescent 7-amino-4-methylcoumarin (AMC), which is released upon proteolytic cleavage. The scissile bond is indicated in red. **C)** Schematic of the truncated Yeast expressed recombinant TMPRSS2, containing the low-density lipoprotein receptor A (LDLRA) domain, scavenger receptor cysteine-rich (SRCR) domain and protease domain used in the biochemical assay.

Camostat (also called FOY-305) is a trypsin-like serine protease inhibitor approved in Japan (as the mesylate salt) for the treatment of pancreatitis and reflux esophagitis^18^. Given its status as an approved agent that is orally administered, safe and well tolerated in humans, and can inhibit cellular entry, camostat mesylate received attention as a drug repurposing candidate. At least eight clinical trials for treating patients are currently underway^19^. It was developed by Ono Pharmaceuticals (Japan, patented in 1977^18, 20^). While a specific report of its development does not appear to be published, it is a highly potent inhibitor of trypsin (IC_50_ ≈ 50 nM)^21^, and it cross-inhibits other proteases. When given as a treatment, it is hydrolysed in plasma (t_1/2_ < 1 min) to 4-(4-guanidinobenzoyloxy)phenylacetic acid (FOY-251), which has shown a similar profile for inhibition of trypsin, thrombin, plasma kallikrein and plasmin^21^. Two other structurally related inhibitors, nafamostat (Torii Pharmaceutical, Japan) and gabexate (also from Ono, FOY-307), are also approved in Japan for treating pancreatitis and show potential for activity against SARS-CoV-2, and trials with nafamostat have also been reported^22^. While TMPRSS2 biochemical assays have been reported for understanding its role in prostate cancer and influenza^23, 24^, these three inhibitors have not yet been directly demonstrated to inhibit TMPRSS2, and mechanism has been inferred by inhibition of SARS-CoV-2 infection in cell models such as the lung-derived human cell line Calu-3^25^. Conversely, bromhexine (BHH), an FDA-approved mucolytic cough suppressant, has been identified within a TMPRSS2 biochemical screen as the most potent hit (IC_50_ = 750 nM) and further shown to demonstrate efficacy in mouse models to reduce metastasis of prostate cancer^24^. With this reported activity as a TMPRSS2 inhibitor, it is being investigated within a COVID19 clinical trial^26^.

In response to the COVID19 public health emergency, we are developing both protein/biochemical and cell-based assays to interrogate several biological targets to enable identification of potential therapeutic leads. Our initial focus is on performing drug repurposing screening for each assay and rapidly sharing the data through the NCATS OpenData portal for COVID19^27^. As part of this effort, we sought to develop a biochemical assay for measuring the activity of TMPRSS2 to enable the evaluation of existing drug repurposing candidates, virtual screening^28-30^, and new inhibitors.

Herein we report the development of a TMPRSS2 fluorogenic biochemical assay and testing of a number of clinical repurposing candidates for COVID19. Activity of TMPRSS2 constructs, and assessment of several substrates was first performed. The best substrate was then used to assess enzyme kinetics and establish a K_m_ value for the substrate, to define assay conditions and demonstrate suitability of the assay in 384- and 1536-well plates. The inhibitors camostat, FOY-251, nafamostat, gabexate and BHH were assessed. To understand their relative activities, we chose camostat, nafamostat and gabexate to be profiled against a panel of human recombinant proteases.

## Results and Discussion

To identify inhibitors of TMPRSS2 that may be used to validate its role in SARS-CoV-2 entry and potentially expedite to clinical trials, we developed a biochemical assay using active TMPRSS2 protease and a fluorogenic peptide substrate (Figure 1B). Initially, we screened six candidate fluorogenic peptide substrates, Boc-Gln-Ala-Arg-AMC^24^, Cbz-D-Arg-Gly-Arg-AMC^23^, Cbz-D-Arg-Pro-Arg-AMC^23^, Boc-Leu-Gly-Arg-AMC^23, 31^, Cbz-Gly-Gly-Arg-AMC^23, 32^ and Ac-Val-Arg-Pro-Arg-AMC, most of which had been demonstrated within the literature to be cleaved by TMPRSS2 and related proteases. Each peptide contains a 7-amino-4-methylcoumarin (AMC) fluorophore that is released following enzymatic cleavage. Candidate substrates were tested to both confirm that the TMPRSS2 construct was biochemically active (thus far commercial protein expressed in HEK293, *E*.*Coli* and in vitro wheat germ constructs have not shown activity, data not shown), and to identify the most avid substrate, indicated by the greatest production of fluorescence from the released fluorogenic product AMC (Figure 2A). The peptide Boc-Gln-Ala-Arg-AMC had 27% released AMC at 60 min, which was the highest observed, and was used for further assay optimization and inhibitor screening with Yeast expressed recombinant TMPRSS2 from Creative BioMart (Figure 1C).

**Figure 2:**
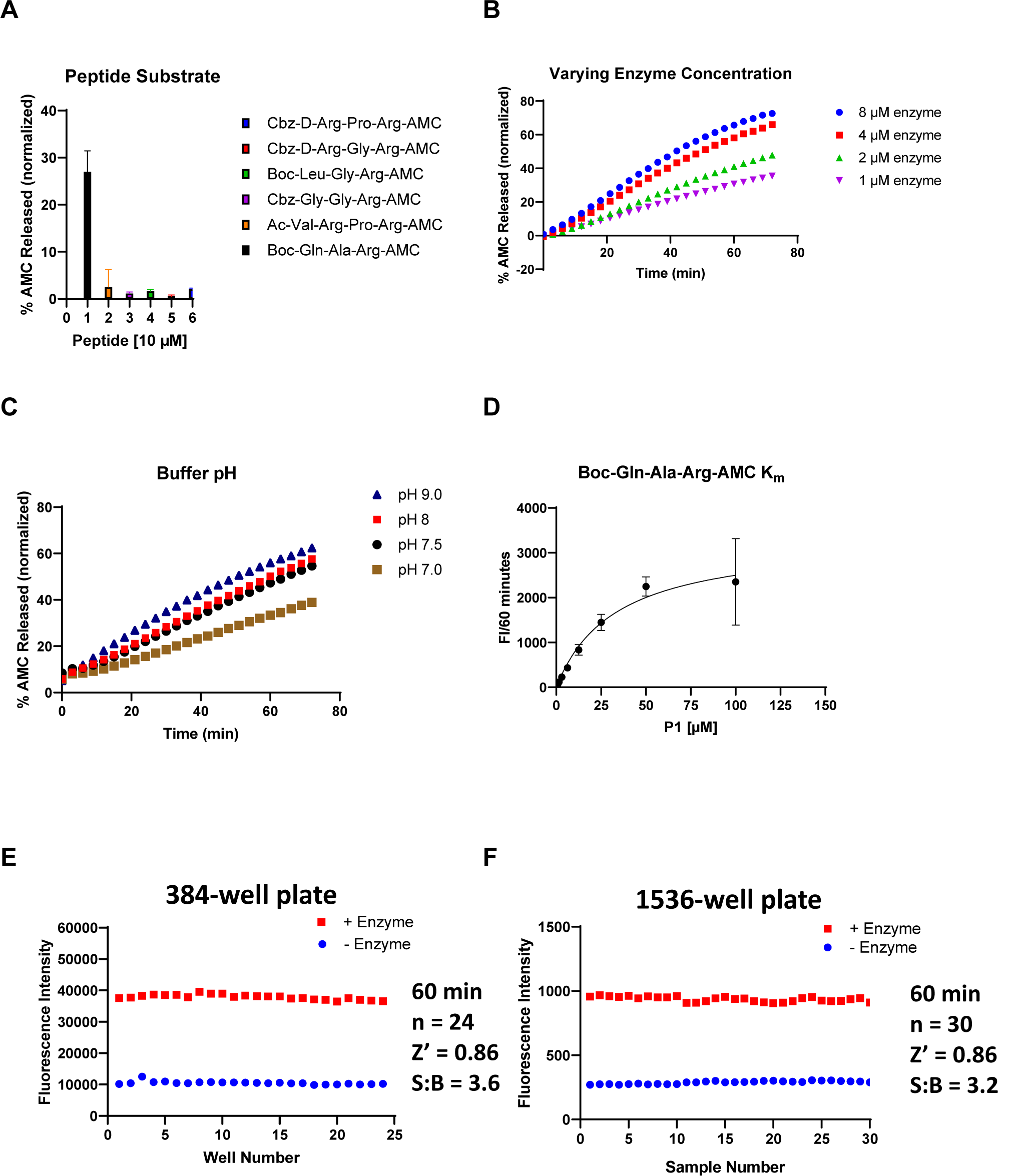
Assessment of enzymatic activity and optimization of TMPRSS2 biochemical assay. **A)** Various AMC-labeled peptides. Peptide [10 µM], TMPRSS2 [1 µM] in Tris-HCl pH8. **B)** TMPRSS2 titration using Boc-Gln-Ala-Arg-AMC peptide [25 µM] in Tris-HCl pH8. **C)** Varying Tris-HCl buffer pH. TMPRSS2 [4 µM], Boc-Gln-Ala-Arg-AMC [25 µM]. **D)** K_m_ of Boc-Gln-Ala-Arg-AMC while TMPRSS2 [1 µM], Tris-HCl pH8. **E)** 384-well plate S:B and Z’. TMPRSS2 [1 µM], Boc-Gln-Ala-Arg-AMC [10 µM], Tris-HCl pH8. **F)** 1536-well plate S:B and Z’. TMPRSS2 [1 µM], Boc-Gln-Ala-Arg-AMC [10 µM], Tris-HCl pH8.

Next, a TMPRSS2 titration was performed at constant substrate concentration (25 µM) to identify an appropriate enzyme concentration that achieves ∼20% substrate cleavage in 60 min, and this was found to be 1 µM (Figure 2B). We then varied assay buffer conditions, such as Tris-HCl buffer pH, DMSO and Tween20 concentrations to further optimize enzymatic activity and determine tolerance to DMSO and Tween20 that are required for inhibitor screening. Noticing that trypsin activity is optimal at pH 7.5 – 8.5^33^, we tested a few different pH > 7 and demonstrated that pH of 9 had highest % released AMC (Figure 2C). However, a pH of 8, which had nearly identical % released AMC, was chosen to proceed for further assay optimization and inhibitor screening. Next, enzymatic activity was shown to be tolerant of Tween20 at 0.01% (data not shown), and DMSO up to 4% (data not shown), well above the DMSO concentrations of <1% v/v typically applied during the testing of inhibitors.

Finally, using our optimized assay buffer conditions, we determined the K_m_ of our peptide substrate to be 33 µM (Figure 2D). The concentration of substrate selected for the biochemical assay was set below K_m_ at 10 µM to ensure susceptibility to competitive inhibitors (detailed protocol provided in Table 1). Positive and negative control conditions were assessed in 384- and 1536-well plate formats to determine a signal:background (S:B) and a Z’ appropriate for HTS. The S:B and Z’ in a 384-well plate were 3.6 and 0.86, respectively (Figure 2E), and in a 1536-well plate the S:B and Z’ were 3.2 and 0.86, respectively (Figure 2F). These data demonstrate appropriate performance for this assay within both plate formats to be useful for HTS.

**Table 1.** Detailed TMPRSS2 biochemical protocol.

| Step # | Process | Notes |
| --- | --- | --- |
| 1 | 20 nL of peptide substrate dispensed into 1536-well plates. | Peptide (dissolved in DMSO) dispensing performed using an ECHO 655 acoustic dispenser (LabCyte). Corning® 1536-well Black Polystyrene, square well, high base, non-sterile, non-treated; Cat #3724 |
| 2 | 20 nL of inhibitor or vehicle control (DMSO) dispensed into 1536-well plates. | Inhibitor or vehicle control (DMSO) dispensing performed using an ECHO 655 acoustic dispenser (LabCyte). |
| 3 | TMPRSS2 diluted in assay buffer dispensed into 1536-well plates. | TMPRSS2 (33.5 μM, 150 nL) in assay buffer (50 mM Tris pH 8, 150 mM NaCl) dispensing performed using a BioRAPTR (Beckman Coulter). Total reaction volume of 5 μL. |
| 4 | Incubate at RT for 1 hr at RT | Final assay conditions are 10 μM peptide and 1 μM TMPRSS2 in assay |
|  |  | buffer (50 mM Tris-HCl, pH 8, 150 mM NaCl) |
| 5 | Read on PHERAstar FSX (BMG Labtech) | Fastest read settings, Fluorescence Intensity module, 340 nm excitation, 440 nm emission) (Cat # 1601A2, BMG Labtech) |

Using the established assay, we tested the inhibition of TMPRSS2 using COVID19-relevant and putative TMPRSS2 inhibitors: camostat, FOY-251, nafamostat, gabexate and BHH (Figure 3). We found that nafamostat (currently in clinical trials) (IC_50_ = 0.27 nM) is the most potent among the inhibitors, while camostat (currently in clinical trials) shows low nM potency (IC_50_ = 6.2 nM) and its primary metabolite, FOY-251, has a ∼5-fold potency loss (IC_50_ = 33.3 nM). Gabexate was significantly less potent (IC_50_ = 130 nM). Notably, we did not detect inhibition by BHH (currently in clinical trials). Due to our reported IC_50_ values being ∼10-100-fold lower than our enzyme concentration (1 µM), based on the manufacturer’s supplied concentration, we used the TMPRSS2 activity (%) from our dose-response curves for camostat and nafamostat to calculate an active TMPRSS2 concentration and apparent dissociation constants for the enzyme-inhibitor complex (K_i_^app^) using the tight-binding equation, also known as Morrison equation (Eq. 1)^34, 35^.

**Figure 3:**
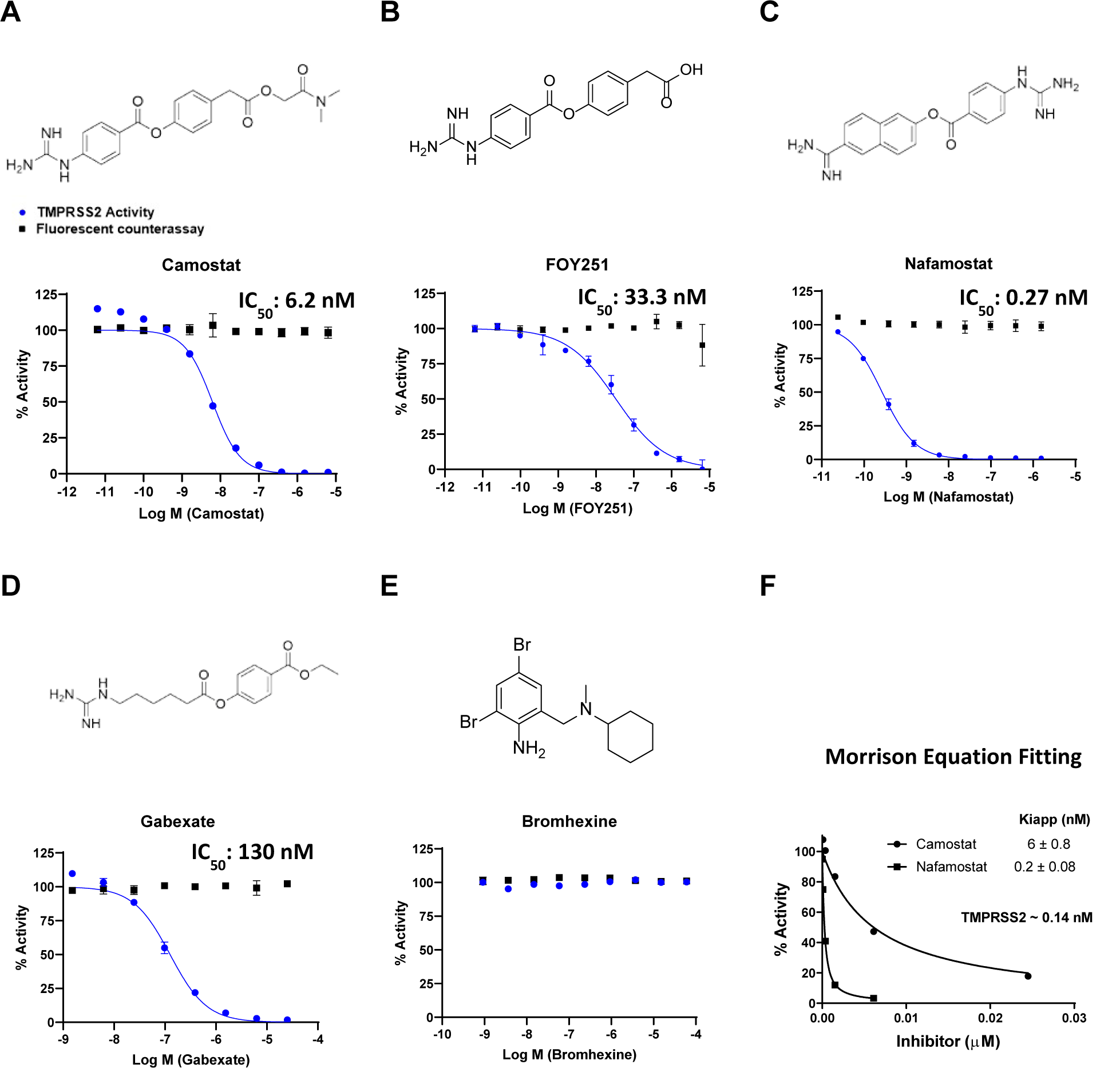
Activity of clinically approved inhibitors against TMPRSS2 (blue), and fluorescent counter-assay (black). The molecular structures and dose-response inhibition of TMPRSS2 by **A)** camostat, **B)** FOY251, **C)** nafamostat, **D)** gabexate and **E)** bromhexine are shown. The calculated concentrations required for 50% inhibition (IC_50_) are displayed in nM. **F)** Calculated active TMPRSS2 concentration and apparent dissociation constants for the enzyme-inhibitor complex (K_i_^app^) by fitting does-response data from camostat and nafamostat to the Morrison equation.

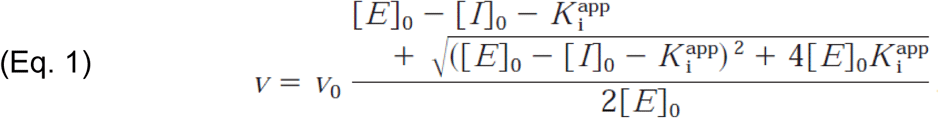

Fitting the TMPRSS2 activity (%) data to this equation while setting S to 1 and K_m_ to 1000 within GraphPad Prism determined an active TMPRSS2 concentration of approximately 0.14 nM and K_i_^app^ values for camostat and nafamostat matching the reported IC_50_ values (Figure 3F). Graphs to determine K_i_^app^ for FOY-251 and gabexate are shown in Supplementary Figure 1. To ensure these inhibitors are not false-positive artifacts of the assay by quenching the fluorescence of the cleaved AMC, a counter-assay to detect quenching of AMC was performed. The counter assay involved addition of inhibitors in various concentrations to AMC held at 1 µM, which approximates 10% enzymatic cleavage resulting in 1 µM of released AMC. The counter assay demonstrated that these inhibitors had no dose-response effects on the fluorescence from AMC, indicating there was no quenching of AMC fluorescence from the inhibitors tested (overlaid data, Figure 3a-e). This data supports the conclusions drawn in cell-based studies that the ability of camostat, nafamostat and gabexate to inhibit cell entry of virus is caused by direct inhibition of TMPRSS2 (though inhibition of other proteases may contribute to cellular activity observed, *vide infra*).

To assess potential activity against other physiologically relevant proteases, camostat, nafamostat and gabexate were profiled by Reaction Biology Corp. against a panel of human recombinant protease assays, spanning multiple enzyme families and classes in 10-point concentration response (Figure 4). Consistent with their similar chemical structures and defined activity as trypsin-like serine protease inhibitors^36^, inhibition patterns were similar among the three inhibitors tested, demonstrating concentration-response inhibition of several trypsin-like serine proteases, including members of the trypsin family (trypsin, kallikreins). All three inhibitors also demonstrated potent activity towards the plasma trypsin-like proteases, plasmin and FXIa, as well as Matriptase 2 (a.k.a. TMPRSS6), a member of the type II transmembrane protease family which includes TMPRSS2^37^. Nafamostat again demonstrated the most potent inhibitory activity, followed by camostat, with gabexate demonstrating the lowest potency. Of the proteases tested, only Kallikrein 12 demonstrated greater sensitivity to gabexete relative to nafamostat and camostat. Dose-response data revealed no inhibitory activity against several matrix metalloproteases (MMP), caspases, and only modest activity against the cysteine protease, Cathepsin S.

**Figure 4:**
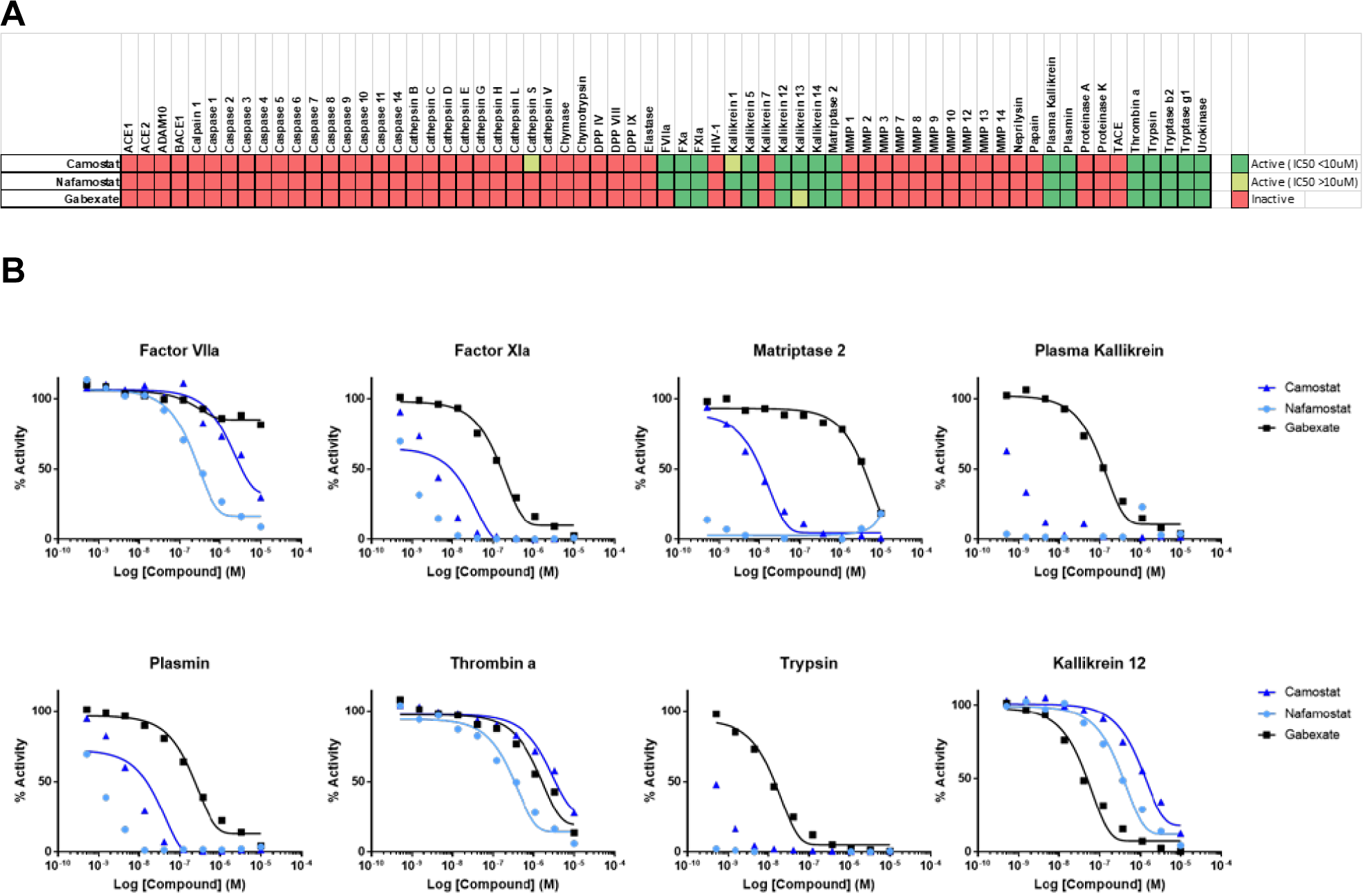
Activity of camostat, nafamostat and gabexate against a panel of proteases. **A)** Compounds were tested against all proteases in dose-response, and activity data was conditionally formatted, dark green = inhibition (IC_50_ ≤10µM), light green = (IC_50_ >10µM), and red = inactive. **B)** Dose-response curves for all three compounds against the eight most sensitive proteases in the panel (full report in Supplemental Reaction Biology Report).

Additional profiling of 40+ human proteases by BPS Bioscience Inc. at single concentration using 10 µM (Supplementary Figure 2) confirmed sensitivity observed by Matriptase 2 and several kallikreins, with highest inhibitory activity against another plasma trypsin-like serine protease, APC, which plays a critical role in coagulation^38^. This diverse panel also included several caspases and ubiquitin specific proteases (USP), all of which demonstrated little to no sensitivity to either inhibitor. The conserved rank order demonstrated here against numerous proteases tested suggests that neither compound conveys improved target selectivity but rather improved potency, which highlights the need for novel, more specific inhibitors of TMPRSS2.

We developed a fluorogenic biochemical assay for measuring recombinant human TMPRSS2 activity for high-throughput screening that can be readily replicated and used to demonstrate inhibition as we show with nafamostat (most potent), camostat, FOY-251 and gabexate. The fluorogenic assay approach taken here has advantages and disadvantages. The assay was readily scaled to 1536-well format for potential high-throughput robotic screening, can be monitored in real-time, and activity is easily detected by the liberation of a fluorophore. The substrate was selected based on the maximal activity of TMPRSS2 against it compared with other substrate candidates, and can be considered a tool substrate, rather than one that is physiologically relevant in the context of the action of TMPRSS2 against its SARS-CoV-2 spike protein cleavage site. A disadvantage that is common to all fluorescence-based assay readouts is the potential for inhibitory compounds from screening to be false-positive artifacts, by quenching the fluorescence of the AMC product, but a simple counter-assay for AMC quenchers can be used to identify false-positives. This counter assay was done on those inhibitors profiled here to demonstrate there was no dose-response quenching of AMC fluorescence (overlaid data, Figure 3a-e).

Two other reports of biochemical assays for TMPRSS2 exist, though their studies were unrelated to the role of TMPRSS2 in SARS-CoV-2 entry. Lucas *et al*. examined TMPRSS2 in the context of prostate cancer. They reported a HTS at a single concentration that produced several hits including their most potent hit, BHH (IC_50_ of 750 nM reported), which is an FDA-approved mucolytic cough suppressant^24^. It has also been included as a potential drug repurposing TMPRSS2 inhibitor and been included in clinical trials^26, 39^. Unfortunately, few details of the assay utilized, scale of assay, or its development were described, but the substrate identified in our study as the most amenable for HTS (Boc-Gln-Ala-Arg-AMC) was also used in the Lucas *et al*. study. Notably, we did not detect inhibition of TMPRSS2 from BHH, and confirmed 99.9% purity of our BHH sample (Supplementary Figure 3). One significant distinction between the reports is the recombinant TMPRSS2 construct used. Both were expressed in yeast, yet the Lucas *et al*. TMPRSS2 (aa 148-492) is without the LDLRA domain, whereas our TMPRSS2 (aa 106-492) composes the full extracellular domain. It is not known how this difference could lead to a discrepancy in inhibition data. In potential agreement with our results, a recent report demonstrated a lack of inhibiting SARS-CoV-2-S-mediated pseudotyped particle entry from bromhexine, suggesting no direct inhibition of TMPRSS2^40^. Meyer *et al*. examined several peptide AMC substrates for TMPRSS2 and used a biochemical assay to assess modified peptide substrates as TMPRSS2 inhibitors, some with observed inhibition constants of approximately 20 nM^23^. They found a high correlation between inhibition constants of inhibitors between TMPRSS2 and matriptase, similar to the cross-inhibition seen from the inhibitors profiled within our assay. Additionally, they report a low active TMPRSS2 concentration in the enzyme stock solution. In our study, the enzyme concentration used was 1 µM based on the manufacturer’s supplied concentration. However, using our dose-response data from camostat and nafamostat for fitting to the Morrison equation allowed us to calculate the active TMPRSS2 concentration to be approximately 0.14 nM.

We report here for the first time the direct biochemical inhibition of TMPRSS2 by camostat and nafamostat, clinical agents of interest in COVID19. Given the clinical trial attention on camostat and nafamostat for treating COVID19, our finding that nafamostat demonstrates greater potency against TMPRSS2 supports its evaluation in clinical trials. A recent molecular dynamics and Markov modeling study explains the increased potency from nafamostat and the molecular mechanism of TMPRSS2 inhibition^41^. Protease profiling revealed activity against a range of trypsin-like serine proteases (and greater potency than against TMPRSS2), but activity was restricted to this protease class and compounds did not generally inhibit other protease classes such as matrix metalloproteases (MMPs), caspases, and ubiquitin-specific proteases (USPs). Camostat and nafamostat development was reported to focus on trypsin, plasmin, and kallikrein^21, 42^, because of the role these targets played in pancreatitis, reflux esophagitis and hyperproteolytic conditions, and activity against these enzymes was certainly observed in the protease panel.

Beyond the evaluation of current inhibitors, we demonstrated acceptable reproducibility and S:B indicating its suitability in a drug repurposing screen and to support the development of new TMPRSS2 inhibitors through virtual screening efforts^28-30^. There are several reasons why a new clinical candidate may be valuable. A number of coronaviruses have been shown to rely on TMPRSS2 for cellular entry (described in the introduction), so a potent, orally available TMPRSS2 inhibitor could be invaluable as a repurposing candidate for treating future emergent coronaviruses. COVID19 is associated with acute respiratory distress syndrome (ARDS). The lung pathology of the ARDS shows microvascular thrombosis and hemorrhage, and has been characterized as disseminated intravascular coagulation (DIC) with enhanced fibrinolysis, or as diffuse pulmonary intravascular coagulopathy^43, 44^. This coagulopathy can lead to pulmonary hypertension and cardiac injury. Trypsin-like serine proteases are involved in SARS-CoV-2 cell entry (TMPRSS2), in the coagulation cascade (APC) and in the enhanced fibrinolysis (plasmin). As shown from profiling the protease panels, camostat, nafamostat and gabexate directly inhibit enzymes involved in all these processes. Clinical development of this class of compounds could thus be directed towards treatment of the infection through inhibition of viral entry, towards treatment of the coagulopathy, or conceivably, both. However, the strategies used to treat viral infection and coagulopathies are very different. The former aims to achieve maximum viral suppression, with dose limited by safety and tolerability. The latter seeks to strike the delicate balance between suppression of thrombosis while managing the inevitable increased risk of bleeding. For this reason, the known clinical trials planned or underway for nafamostat and camostat can be divided into three categories: those whose primary endpoint is focused on limiting or preventing infection, those whose primary endpoint is focused on management of ARDS and advanced disease, and those which may capture treatment benefit through either mechanism.

Nafamostat is approved and marketed in Japan and S. Korea and is typically prescribed to treat acute symptoms of pancreatitis and to treat DIC^45^. It is administered by IV infusion and has a plasma half-life (t_1/2_) of about 23 min^46^. Clinical trials to test the hypothesis that nafamostat can lower lung function deterioration and need for intensive care admission in COVID-19 patients, and that nafamostat can improve the clinical status of COVID-19 patients with pneumonia have been registered^22^. Camostat is approved in Japan for the treatment of acute symptoms of chronic pancreatitis and postoperative reflux esophagitis^47^. The treatment regimen is typically 200 mg PO, every eight hours. Camostat is a pro-drug as the parent drug is not detected in plasma. The terminal ester is rapidly hydrolysed in plasma (t_1/2_ < 1 min) to 4-(4-guanidinobenzoyloxy)phenylacetic acid (FOY-251) which has a mean terminal half-life of 1 h^48^. FOY-251 and camostat are reported to have similar activities against trypsin, thrombin, plasma kallikrein and plasmin^21^, and here we show a ∼5-fold decreased potency by FOY-251 (IC_50_ = 33.3 nM) compared to camostat (IC_50_ = 6.2 nM) for TMPRSS2 inhibition. There are eight registered clinical trials studying camostat, of which only one is focused on coagulopathy^19^. The other trials are focused on early, mild, or moderate disease. Gabexate is an IV drug approved for marketing in Italy and Japan and was shown to be effective in treating patients with sepsis-associated DIC and treating acute pancreatitis^49^. It is clearly much less potent than nafamostat and there are no registered COVID-19 clinical studies with it. In summary, of the three drugs in the class, nafamostat is being studied as the preferred drug in an ICU setting as it can be titrated against coagulation markers as a treatment for coagulopathy, while considering its antiviral effect as a bonus. In outpatient, early diagnosis, or prophylactic settings, camostat is being studied predominantly with the primary purpose as an antiviral. In these latter settings there is room for new, selective TMPRSS2 inhibitors which could achieve higher levels of inhibition without incurring a bleeding risk.

The biochemical TMPRSS2 assay we disseminate here is a simple and HTS-amenable approach to TMPRSS2 inhibitor therapeutic development. Clinical trials of nafamostat for COVID19 have been reported, and we believe it warrants evaluation given its superior activity over camostat, as demonstrated herein.

## Methods

### Reagents

Recombinant Human TMPRSS2 protein expressed from Yeast (human TMPRSS2 residues 106-492, N-terminal 6x His-tag) (Cat # TMPRSS2-1856H) was acquired from Creative BioMart (Shirley, NY). Peptides obtained from Bachem include: Boc-Leu-Gly-Arg-AMC. acetate (Cat#: I-1105), Boc-Gln-Ala-Arg-AMC. HCl (Cat#: I-1550), Ac-Val-Arg-Pro-Arg-AMC. TFA (Cat#: I-1965), Cbz-Gly-Gly-Arg-AMC. HCl (Cat#: I-1140). Peptides custom ordered from LifeTein (Somerset, NJ) include: Cbz-D-Arg-Gly-Arg-AMC, Cbz-D-Arg-Pro-Arg-AMC.

### Fluorogenic Peptide Screening Protocol – 384-well plate

To a 384-well black plate (Greiner 781900) was added Boc-Gln-Ala-Arg-AMC (62.5 nL) and inhibitor (62.5 nL) using an ECHO 655 acoustic dispenser (LabCyte). To that was added TMPRSS2 (750 nL) in assay buffer (50 mM Tris pH 8, 150 mM NaCl) to give total reaction volume of 25 µL. Following 1 hr incubation at RT, detection was done using the PHERAstar with excitation: 340 nm and emission: 440 nm.

### Fluorescence counter assay – 384-well plate

To a 384-well black plate (Greiner 781900) was added 7-amino-methylcoumarin (62.5 nL) and inhibitor or DMSO (62.5 nL) using an ECHO 655 acoustic dispenser (LabCyte). To that was added assay buffer (50 mM Tris pH 8, 150 mM NaCl) to give total reaction volume of 25 µL. Detection was done using the PHERAstar with excitation: 340 nm and emission: 440 nm. Fluorescence was normalized relative to a negative control containing DMSO-only wells (0% activity, low fluorescence) and a positive control containing AMC only (100% activity, high fluorescence). An inhibitor causing fluorescence quenching would be identified as having a concentration-dependent decrease on AMC fluorescence.

### Fluorogenic Peptide Screening Protocol – 1536-well plate

To a 1536-well black plate was added Boc-Gln-Ala-Arg-AMC substrate (20 nL) and inhibitor (20 nL) using an ECHO 655 acoustic dispenser (LabCyte). To that was dispensed TMPRSS2 (150 nL) in assay buffer (50 mM Tris pH 8, 150 mM NaCl) using a BioRAPTR (Beckman Coulter) to give total reaction volume of 5 µL. Following 1 hr incubation at RT detection was done using the PHERAstar with excitation: 340 nm and emission: 440 nm.

### TMPRSS2 Assay Protocol

The TMPRSS2 biochemical assay was performed according to the assay protocol shown in Table 1.

### Data Process and Analysis

To determine compound activity in the assay, the concentration-response data for each sample was plotted and modeled by a four-parameter logistic fit yielding IC_50_ and efficacy (maximal response) values. Raw plate reads for each titration point were first normalized relative to a positive control containing no enzyme (0% activity, full inhibition) and a negative control containing DMSO-only wells (100% activity, basal activity). Data normalization, visualization and curve fitting were performed using Prism (GraphPad, San Diego, CA).

### Protease profiling

Camostat, nafamostat and gabexate were assessed for inhibition against panels of recombinant human proteases by commercial services from Reaction Biology Corp and BPS Biosciences. The Reaction Biology Corp profile tested in a 10-dose IC_50_ with a 3-fold serial dilution starting at 10 μM against 65 proteases. The BPS Biosciences profile was against 48 proteases at a single concentration of 10 μM.

## Supporting information

Supplement BPS Bioscience Report

Supplement Reaction Biology Report

## Supporting Information

The Supporting Information is available free of charge at _.

Figure S1. Calculating Kiapp for gabexate and FOY-251 using the Morrison Equation

Figure S2. Camostat, nafamostat and gabexate inhibition of recombinant human proteases from a BPS Bioscience protease panel

Figure S3. LC-MS chromatogram of bromhexine (PDF)

Supplemental: Reaction Biology Report

Supplemental: BPS Bioscience Report

