## Supplement BPS Bioscience Report for "An Enzymatic TMPRSS2 Assay for Assessment of Clinical Candidates and Discovery of Inhibitors as Potential Treatment of COVID-19"

**Assay Report**

**Protease Inhibitor Assays**

Enzymatic Study of Three Compounds from NCATS/NIH

**NIH_Protease_200603**

**Protease Inhibitor Assays**

Study Sponsor:	NCATS/NIH

Attention:	Jonathan Shrimp

Address:	NCATS/NIH

9800 Medical Center Drive, Rm 1034B

Rockville, MD 20850

Study Director:	Henry Zhu, Ph.D.

Testing Facility:	BPS Bioscience Inc.

6042 Cornerstone Court West, Ste. B

San Diego, CA 92121

USA

Study Period:

Report Version:	1

Report Date:	June 3, 2020

**Study Director**

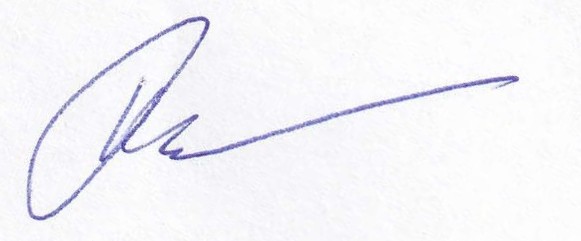

____________________

Pavel Shashkin, Ph.D.

Director

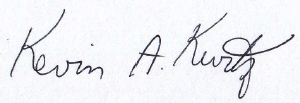

____________________

Kevin Kurtz

Sr. Scientist II.

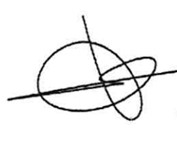

____________________

Ivan Caballero

Sr. Scientist II.

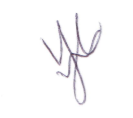

____________________

Victoria Castillo

Research Associate II.

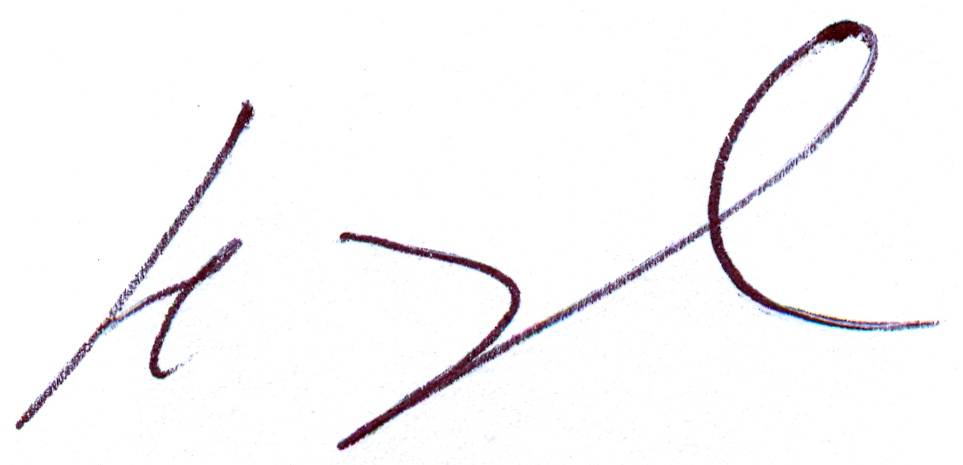

_____________________

Henry Zhu, Ph.D.

President

06-03-2020

______________________

Date

**CONTENTS**

[Protease Inhibitor Assays	1](../../../../S:/Reports/2020/NCATS-NIH%20Protease%2020200514.doc" \l "__RefHeading___Toc39743437)

[Enzymatic Study of Three Compounds from NCATS/NIH	1](../../../../S:/Reports/2020/NCATS-NIH%20Protease%2020200514.doc" \l "__RefHeading___Toc39743438)

[Protease Inhibitor Assays	2](../../../../S:/Reports/2020/NCATS-NIH%20Protease%2020200514.doc" \l "__RefHeading___Toc39743439)

[Study Director	3](../../../../S:/Reports/2020/NCATS-NIH%20Protease%2020200514.doc" \l "__RefHeading___Toc39743440)

1.	Purpose of the Study	6

2.	Materials and Methods	7

2.1	Materials	7

2.2	Compounds	8

2.3	Experimental Conditions	9

2.3.1	Enzymes and Substrates	9

2.3.2	Assay Conditions	11

2.3.3	Data Analysis	13

3.	Assay Results	14

3.1.	Summary of the Inhibitory Effects of the Compounds on Individual Protease Activities	14

3.2.	Results of the Effects of the Compounds on Individual Protease Activity	16

3.2.1.	BACE1	16

3.2.2.	Caspase-3	17

3.2.3.	Caspase-6	18

3.2.4.	Caspase-7	19

3.2.5.	Caspase-8	20

3.2.6.	Caspase-9	21

3.2.7.	Renin	22

3.2.8.	3CL Protease	23

3.2.9.	A20	24

3.2.10.	Ataxin-3	25

3.2.11.	Cathepsin B	26

3.2.12.	Cathepsin F	27

3.2.13.	Cathepsin L	28

3.2.14.	Cathepsin S	29

3.2.15.	Cathepsin V	30

3.2.16.	MALT1	31

3.2.17.	OTUD6Bc	32

3.2.18.	UCHL1	33

3.2.19.	UCHL3	34

3.2.20.	USP2	35

3.2.21.	USP5	36

3.2.22.	USP7	37

3.2.23.	USP8	38

3.2.24.	USP10	39

3.2.25.	USP14	40

3.2.26.	ADAM17	41

3.2.27.	MMP1	42

3.2.28.	MMP2	43

3.2.29.	MMP3	44

3.2.30.	MMP7	45

3.2.31.	MMP8	46

3.2.32.	MMP9 (Q279R)	47

3.2.33.	MMP10	48

3.2.34.	MMP13	49

3.2.35.	DPP3	50

3.2.36.	DPP4	51

3.2.37.	DPP7	52

3.2.38.	DPP8	53

3.2.39.	DPP9	54

3.2.40.	FAP	55

3.2.41.	HCV1a (D168V)	56

3.2.42.	HCV1b	57

3.2.43.	HCV1b (D168V)	58

3.2.44.	HCV1b (R155K)	59

3.2.45.	HCV1b (R155Q)	60

3.2.46.	HCV2a	61

3.2.47.	POP	62

3.2.48.	APC	63

4.	Quality Assurance Statement	64

1. Purpose of the Study

The purpose of the study is to determine the effects of three compounds from NCATS/NIH on the enzymatic activities of recombinant human BACE1, Caspase-3, Caspase-6, Caspase-7, Caspase-8, Caspase-9, Renin, 3CL, A20, Ataxin-3, Cathepsin B, Cathepsin F, Cathepsin L, Cathepsin S, Cathepsin V, MALT1, OTUD6Bc, UCHL1, UCHL3, USP2, USP5, USP7, USP8, USP10, USP14, ADAM17, MMP1, MMP2, MMP3, MMP7, MMP8, MMP9 (Q279R), MMP10, MMP13, DPP3, DPP4, DPP7, DPP8, DPP9, FAP, HCV1a (D168V), HCV1b, HCV1b (D168V), HCV1b (R155K), HCV1b (R155Q), HCV2a, POP, and APC using an in vitro enzymatic assay.

2. Materials and Methods
   2.1 Materials

Verubecestat is purchased from SelleckChem (catalog number S8173).

Z-Ac-DNLD-CHO is purchased from Millipore (catalog number 218832).

Ac-IETD-CHO is purchased from Millipore (catalog number 218773).

Aliskiren hydrochloride is from BPS Bioscience (BPS number 27303).

GC376 is purchased from Aobious (catalog number AOB36447).

Ubiquitin-Aldehyde is purchased from Boston Biochem (catalog number U201).

E-64 Santa is purchased from Cruz Biotechnology (catalog number 66701-25-5).

Cystatin C is purchased from Enzo Life science (catalog number BML-SE479).

Z-VRPR-FMK is purchased from Tocris (Catalog #4645).

TAPI-2 is purchased from Cayman Chemical (catalog number 14695).

NNGH is purchased from Sigma Aldrich (catalog number SML0584).

Batimastat is purchased from Cayman Chemical (catalog number 14742).

Spinorphin is purchased from Tocris (catalog number 2931).

Sitagliptin from BPS Bioscience (BPS number 27106).

KR62436 is purchased from Sigma-Aldrich (catalog number K4264).

SP-13786 is purchased from MedChemExpress (catalog number HY-100684).

Denoprevir is purchased from Selleckchem (catalog number S1183).

Z-Pro-Pro-CHO is purchased from Enzo Life science (catalog number BML-PI112).

Dabigatran is purchased from Selleckchem (catalog number S2196).

Z-LEHD-FMK is purchased from Selleckchem (catalog number S7313).

Boc-VPR-AMC Fluorogenic Peptide Substrate is purchased from R & D Systems (catalog number ES011).

BACE substrate is purchased from R & D Systems (catalog number ES004).

Caspase 3 substrate is purchased from Anaspec (catalog number 25273-5).

Ac-VEID-AFC is purchased from Enzo Lifesciences (catalog number ALX-260-111).

Ac-IETD-AFC is purchased from Enzo Lifesciences (catalog number ALX-260-110).

Ac-LEHD-AFC is purchased from Enzo Lifesciences (catalog number ALX-260-116).

Z-Leu-Arg-AMC is purchased from Enzo Lifesciences (catalog number BML-P229-0010).

Ubiquitin-AMC is purchased from Boston Biochem (catalog number U550).

Z-Arg-Arg-AMC is purchased from BACHEM (catalog number 4004789.0050).

Fluorogenic DPP substrate 1 (Ala-Pro-AMC) is from BPS Bioscience (BPS number 80305).

Ac-LRSR-AMC is purchased from Peptides International (catalog number MCA-3952-PI).

390 MMP FRET Substrate I is purchased from Anaspec (catalog number 27077).

Renin assay kit is from BPS Bioscience (BPS number 80211).

HCV Protease FRET Substrate (RET S1) is purchased from Anaspec (catalog number AS-22991).

2.2 Compounds

The test compound is supplied by NCATS/NIH.

| Compound I.D. | Compound Supplied | Stock Concentration | Dissolving Solvent | Test Range (μM) | Intermediate Dilution |
| --- | --- | --- | --- | --- | --- |
| NCGC00167526 (Camostat) | Solution | 10 mM | DMSO | 10 | 10 % DMSO in Assay Buffer |
| NCGC00160398 (Nafamostat) | Solution | 10 mM | DMSO | 10 | 10 % DMSO in Assay Buffer |
| NCGC00025297 (Gabexate) | Solution | 10 mM | DMSO | 10 | 10 % DMSO in Assay Buffer |
| Verubecestat* | Solid | 10 mM | DMSO | 0.3, 0.03, 0.003 | 10 % DMSO in Assay Buffer |
| Z-Ac-DNLD-CHO* | Solid | 10 mM | DMSO | 0.5, 0.1, 0.05, 0.01, 0.001, 0.005 | 10 % DMSO in Assay Buffer |
| Ac-IETD-CHO* | Solid | 10 mM | DMSO | 0.5, 0.1, 0.05, 0.01, 0.001, 0.005 | 10 % DMSO in Assay Buffer |
| Aliskiren* | Solid | 10 mM | DMSO | 0.01, 0.001, 0.0001 | 10 % DMSO in Assay Buffer |
| GC376* | Solid | 10 mM | DMSO | 5, 0.5, 0.05 | 10 % DMSO in Assay Buffer |
| Ubiquitin-Aldehyde* | Solution | 160 μM | Buffer | 100 – 0.0001 | 10 % DMSO in Assay Buffer |
| E-64* | Solid | 10 mM | DMSO | 0.5, 0.05, 0.005 | 10 % DMSO in Assay Buffer |
| Cystatin C* | Solid | 10 mM | DMSO | 100, 10, 1 | 10 % DMSO in Assay Buffer |
| Z-VRPR-FMK* | Solid | 10 mM | DMSO | 0.1, 0.01, 0.001 | 10 % DMSO in Assay Buffer |
| TAPI-2* | Solid | 10 mM | DMSO | 10, 1, 0.1 | 10 % DMSO in Assay Buffer |
| NNGH* | Solid | 10 mM | DMSO | 1, 0.1, 0.01, 0.001 | 10 % DMSO in Assay Buffer |
| Batimastat* | Solid | 10 mM | DMSO | 0.01, 0.001, 0.0001 | 10 % DMSO in Assay Buffer |
| Spinorphin* | Solid | 10 mM | DMSO | 10, 0.1, 0.01 | 10 % DMSO in Assay Buffer |
| Sitagliptin* | Solid | 10 mM | DMSO | 0.1, 0.01, 0.001 | 10 % DMSO in Assay Buffer |
| KR62436* | Solid | 10 mM | DMSO | 100, 10, 1, 0.1 | 10 % DMSO in Assay Buffer |
| SP-13786* | Solid | 10 mM | DMSO | 0.1, 0.01, 0.001 | 10 % DMSO in Assay Buffer |
| Denoprevir* | Solid | 10 mM | DMSO | 1, 0.1, 0.01, 0.001, 0.0001 | 10 % DMSO in Assay Buffer |
| Z-Pro-Pro-CHO* | Solid | 10 mM | DMSO | 0.1, 0.01, 0.001 | 10 % DMSO in Assay Buffer |
| Dabigatran* | Solution | 10 mM | DMSO | 20, 2, 0.2 | 10 % DMSO in Assay Buffer |
| Z-LEHD-FMK* | Solid | 10 mM | DMSO | 10, 1, 0.1 | 10 % DMSO in Assay Buffer |

*Reference compounds.

2.3 Experimental Conditions
    2.3.1 Enzymes and Substrates

| Assay | Catalog # | Enzyme Lot # | Enzyme Used (ng) / Reaction | Substrate |
| --- | --- | --- | --- | --- |
| BACE1 | 71657 | 170307 | 100 | 7.5 µM BACE1 Substrate |
| Caspase-3 | 80500 | 130226-G1 | 1 | 1 µM Caspase 3 Substrate |
| Caspase-6 | 80113 | 130222-2 | 15 | 1 µM Ac-VEID-AFC |
| Caspase-7 | 70000 | 130116 | 3 | 1 µM Caspase 3 Substrate |
| Caspase-8 | 80114 | 130215-2 | 5 | 1 µM Ac-LEHD-AFC |
| Caspase-9 | 80115 | 190306 | 2000 | 1 µM Ac-LEHD-AFC |
| Renin | 80200 | 161221 | 4 | 5 μM Renin Substrate |
| 3CL Protease | 100707 | 200416-1 | 150 | 50 μM 3CL Protease Substrate |
| A20 | 80408 | 171109 | 100 | 100 nM Ub-AMC |
| Ataxin3 | 80399 | 180817 | 2000 | 100 nM Ub-AMC |
| Cathepsin B | 80001 | 161019 | 0.4 | 5 μM Z-Leu-Arg-AMC |
| Cathepsin F | 80003 | 160921 | 100 | 5 μM Z-Leu-Arg-AMC |
| Cathepsin L | 80005 | 170620-1 | 0.4 | 5 μM Z-Leu-Arg-AMC |
| Cathepsin S | 80008 | 170214 | 20 | 5 μM Z-Leu-Arg-AMC |
| Cathepsin V | 80009 | 160823 | 2 | 5 μM Z-Leu-Arg-AMC |
| MALT1 | 100360 | 190722-1 | 150 | 10 μM Ac-LRSR-AMC |
| OTUD6B | 80407 | 170926-1 | 400 | 100 nM Ub-AMC |
| UCHL1 | 80351 | 100826 | 0.5 | 100 nM Ub-AMC |
| UCHL3 | 80353 | 190212-1 | 0.01 | 100 nM Ub-AMC |
| USP2 | 80352 | 141212 | 5 | 100 nM Ub-AMC |
| USP5 | 80355 | 120213 | 20 | 100 nM Ub-AMC |
| USP7 | 80395 | 141020-H2 | 5 | 100 nM Ub-AMC |
| USP8 | 80358 | 140521-A | 40 | 100 nM Ub-AMC |
| USP10 | 80360 | 140501-1 | 200 | 100 nM Ub-AMC |
| USP14 | 80364 | 120227 | 20 | 100 nM Ub-AMC |
| ADAM17 | - | 200529 | 20 | 1 μM ADAM17 Substrate |
| MMP1 | 80214 | 130730 | 10 | 10 μM  390 MMP FRET Substrate I |
| MMP2 | 80213 | 160720 | 4 | 2 μM  390 MMP FRET Substrate I |
| MMP3 | 11346 | 130730 | 100 | 2 μM  390 MMP FRET Substrate I |
| MMP7 | - | - | 10 | 2 μM  390 MMP FRET Substrate I |
| MMP8 | 100552 | 191217-1 | 4 | 1 μM  390 MMP FRET Substrate I |
| MMP9  (Q279R) | 80215 | 130730 | 2 | 1 μM  390 MMP FRET Substrate I |
| MMP10 | 100565 | 200115 | 100 | 2 μM  390 MMP FRET Substrate I |
| MMP13 | 11345 | 140219 | 100 | 2 μM  390 MMP FRET Substrate I |
| DPP3 | 80030 | 191211 | 10 | 5 μM DPP Substrate 2  (Arg-Arg-AMC) |
| DPP4 | 80040 | 100707 | 0.2 | 5 μM DPP Substrate 1  (Ala-Pro-AMC) |
| DPP7 | 80070 | 111207-1 | 10 | 5 μM DPP Substrate 1  (Ala-Pro-AMC) |
| DPP8 | 80080 | 190312-G2 | 10 | 5 μM DPP Substrate 1  (Ala-Pro-AMC) |
| DPP9 | 80090 | 2000 | 2.5 | 5 μM DPP Substrate 1  (Ala-Pro-AMC) |
| FAP | 80100 | 190620 | 250 | 5 μM DPP Substrate 1  (Ala-Pro-AMC) |
| HCV1a (D168V) | 80103 | 130920-G2 | 50 | 5 μM HCV Substrate |
| HCV1b | 80107 | 120817-GC | 20 | 5 μM HCV Substrate |
| HCV1b (D168V) | 80110 | 171101 | 20 | 5 μM HCV Substrate |
| HCV1b (R155K) | 80109 | 120928 | 100 | 5 μM HCV Substrate |
| HCV1b (R155Q) | 80104 | 170118C | 400 | 5 μM HCV Substrate |
| HCV2a | 80108 | 120928 | 5 | 5 μM HCV Substrate |
| POP | 80105 | 160513-E | 150 | 5 μM DPP Substrate 1  (Ala-Pro-AMC) |
| APC | 70019 | 130620-E1D | 20 | 5 µM Boc-VPR-AMC |

2.3.2 Assay Conditions

A compound solution ten-fold higher than the final concentration was prepared with 10 % DMSO in assay buffer and 5 µl of the dilution was added to a 50 µl reaction so that the final concentration of DMSO is 1 % in all of reactions. All of control samples, including background and no compound controls, also contain 1 % DMSO.

For activated protein C assay, the enzymatic reactions were conducted in duplicate at room temperature for 30 minutes in a 50 µl mixture containing 50 mM Tris-HCl, pH 8.5, 150 mM NaCl, 10 mM CaCl2, 0.01 % Brij-35, 10 µM Boc-VPR-AMC (see 2.3.1), activated Protein C enzyme (see 2.3.1) and a test compound (see 2.2). Fluorescence intensity was measured at an excitation of 380 nm and an emission of 460 nm using a Tecan Infinite M1000 microplate reader.

For BACE assay, BACE1 FRET Assay Kit (BPS catalogue number 71656) was used according to the kit manual.

For caspase assays, the enzymatic reactions were conducted in duplicate at room temperature for 30 minutes in a 50 µl mixture containing 10 mM HEPES buffer, pH 7.4, 10 mM EDTA, 0.05 % Chaps, 5 mM DTT, 1 µM substrate (see 2.3.1), a caspase enzyme (see 2.3.1) and a test compound (see 2.2). Fluorescence intensity was measured at an excitation of 380 nm and an emission of 505 nm using a Tecan Infinite M1000 microplate reader.

For Cathepsin assays, the enzymatic reactions were conducted in duplicate at room temperature for 30 minutes in a 100 µl mixture containing 50 mM MES buffer, pH 5.0, 100 mM NaCl, 5 mM DTT, a Cathepsin substrate (see 2.3.1), a Cathepsin enzyme (see 2.3.1) and a test compound (see 2.2). Fluorescence intensity was measured at an excitation of 360 nm and an emission of 460 nm using a Tecan Infinite M1000 microplate reader.

For deubiquitinase assays, the enzymatic reactions were conducted in duplicate at room temperature for 30 minutes in a 50 µl mixture containing 50 mM Tris-HCl, pH 7.4, 0.5 mM EDTA, 0.05 % Tween 20, 1 mM DTT, 100 nM Ubiquitin-AMC substrate (see 2.3.1), a deubiquitinase enzyme (see 2.3.1) and a test compound (see 2.2). Fluorescence intensity was measured at an excitation of 360 nm and an emission of 460 nm using a Tecan Infinite M1000 microplate reader.

For DPP assays, the enzymatic reactions were conducted in duplicate at room temperature for 30 minutes in a 50 µl mixture containing DPP assay buffer, 5 µM substrate (see 2.3.1), a DPP enzyme (see 2.3.1) and a test compound (see 2.2). Fluorescence intensity was measured at an excitation of 360 nm and an emission of 460 nm using a Tecan Infinite M1000 microplate reader.

For HCV assay, the enzymatic reactions were conducted in duplicates at room temperature for 30 minutes in a 100 μl mixture containing 50 mM Tris-HCl, pH 7.4, 150 mM NaCl, 10 % Glycerol, 5 mM DTT, an HCV enzyme (see 2.3.1) and a test compound (see 2.2). Fluorescence intensity was measured at an excitation of 355 nm and an emission of 485 nm using a Tecan Infinite M1000 microplate reader.

For MMP assays, the enzyme is diluted to 200 µg/ml in 50 mM HEPES buffer, pH 7.4, 10 mM CaCl_2_, 0.05 % Brij-35, and 1 mM APMA, and activated at 37ºC for 2 hours. The enzymatic reactions were conducted in duplicate at room temperature for 30 minutes in a 50 µl mixture containing 50 mM HEPES buffer, pH 7.4, 10 mM CaCl_2_, 0.05 % Brij-35, an MMP substrate (see 2.3.1), an MMP enzyme (see 2.3.1) and a test compound (see 2.2). Fluorescence intensity was measured at an excitation of 328 nm and an emission of 393 nm using a Tecan Infinite M1000 microplate reader.

For Renin assay, the enzymatic reactions were conducted in duplicate at room temperature for 30 minutes in a 50 µl mixture containing 50 mM Tris-HCl, pH 7.4, 100 mM NaCl, 10 mM MgCl_2_, 0.05 % Tween 20, 5 µM Renin substrate (see 2.3.1), Renin enzyme, and a test compound (see 2.2). Fluorescence intensity was measured at an excitation of 490 nm and an emission of 520 nm using a Tecan Infinite M1000 microplate reader.

For MALT1 assay, the enzymatic reactions were conducted in duplicate at room temperature for 60 minutes in a 50µl mixture containing 50 mM HEPES, pH 6.8, 150 mM NaCl, 1 M Na Citrate, 0.05 % CHAPS, 10 µM Ac-LRSR-AMC (see 2.3.1), MALT1 enzyme, and a test compound (see 2.2). Fluorescence intensity was measured at an excitation of 360 nm and an emission of 460 nm using a Tecan Infinite M1000 microplate reader.

For 3CL Protease assay, 30 μl of 3CL Protease, and 10 μl of compound diluted in assay buffer were pre-mixed in assay well. Mixture was incubated for 30 min at room temperature with slow shaking. The reaction was initiated by adding 10 μl of 50 μM 3CL Protease substrate (50 μM final, see **2.3.1**). Compounds were originally diluted in DMSO, then in 3CL Protease assay buffer. Final concentration of DMSO was 1%. Total volume was 50 μl. Fluorescence was measured using a M1000 Tecan microplate reader (λ_exc_=360 nm, λ_em_=460 nm) after overnight reaction. Fluorescence was also measured before the substrate addition to obtain data for the intrinsic compound fluorescence. These data were subtracted from data after overnight incubation to get net values corrected to the compound fluorescence.

2.3.3 Data Analysis

All of the enzyme activity assays were performed in duplicates at each concentration. The fluorescent intensity data were analyzed using the computer software, Graphpad Prism. In the absence of the compound, the fluorescent intensity (F_t_) in each data set was defined as 100 % activity. In the absence of the enzyme, the fluorescent intensity (F_b_) in each data set was defined as 0 % activity. The percent activity in the presence of each compound was calculated according to the following equation: % activity = (F-F_b_)/(F_t_-F_b_), where F = the fluorescent intensity in the presence of the compound.

The values of percentage activity were plotted on a bar graph.

3. Assay Results
   3.1. Summary of the Inhibitory Effects of the Compounds on Individual Protease Activities

The percentage inhibition of the three compounds against Proteases is summarized on Table 3.1. Compound NCGC00160398 was fluorescent for most assays and background was subtracted.

**Table 3.1	Inhibitory Effects of the Compounds on Protease Activities**

| **Enzyme** | **% Inhibition** | | | | | |
| --- | --- | --- | --- | --- | --- | --- |
|  | **NCGC00167526 (Camostat), 10 μM** | **NCGC00160398 (Nafamostat), 10 μM*** | **NCGC00025297 (Gabexate), 10 μM** | **Reference**  **(~0.1 x IC_50_)** | **Reference**  **(~1 x IC_50_)** | **Reference**  **(~10 x IC_50_)** |
| **BACE1** | 4 | 0 | 0 | 6 | 52 | 95 |
| **Caspase-3** | 0 | 0 | 2 | 10 | 25 | 80 |
| **Caspase-6** | 3 | 4 | 0 | 8 | 34 | 93 |
| **Caspase-7** | 0 | 1 | 1 | 8 | 43 | 89 |
| **Caspase-8** | 0 | 0 | 1 | 0 | 38 | 78 |
| **Caspase-9** | 0 | 0 | 7 | 7 | 50 | 86 |
| **Renin** | 1 | 45 | 0 | 0 | 11 | 70 |
| **3CL** | 6 | 13 | 3 | 11 | 62 | 96 |
| **A20** | 4 | 13 | 4 | 11 | 54 | 96 |
| **Ataxin-3** | 3 | 0 | 3 | 30 | 67 | 96 |
| **Cathepsin B** | 0 | 48 | 0 | 32 | 77 | 97 |
| **Cathepsin F** | 0 | 0 | 2 | 20 | 73 | 89 |
| **Cathepsin L** | 2 | 57 | 0 | 13 | 39 | 84 |
| **Cathepsin S** | 3 | 8 | 4 | 18 | 81 | 97 |
| **Cathepsin V** | 1 | 27 | 3 | 17 | 57 | 94 |
| **MALT1** | 48 | 49 | 46 | 9 | 56 | 97 |
| **OTUD6Bc** | 1 | 9 | 3 | 6 | 24 | 100 |
| **UCHL1** | 7 | 0 | 5 | 7 | 20 | 71 |
| **UCHL3** | 3 | 6 | 4 | 4 | 33 | 89 |
| **USP2** | 1 | 0 | 2 | 5 | 55 | 96 |
| **USP5** | 1 | 0 | 1 | 2 | 57 | 97 |
| **USP7** | 0 | 0 | 0 | 12 | 56 | 97 |
| **USP8** | 0 | 5 | 0 | 11 | 35 | 85 |
| **USP10** | 2 | 0 | 3 | 5 | 43 | 83 |
| **USP14** | 0 | 0 | 1 | 19 | 72 | 100 |
| **ADAM17** | 6 | 5 | 7 | 0 | 13 | 73 |
| **MMP1** | 3 | 0 | 2 | 2 | 26 | 77 |
| **MMP2** | 83 | 0 | 26 | 24 | 71 | 100 |
| **MMP3** | 27 | 0 | 18 | 20 | 80 | 100 |
| **MMP7** | 2 | 5 | 3 | 4 | 31 | 94 |
| **MMP8** | 33 | 0 | 7 | 5 | 43 | 91 |
| **MMP9 (Q279R)** | 8 | 11 | 13 | 27 | 74 | 96 |
| **MMP10** | 29 | 0 | 12 | 25 | 63 | 91 |
| **MMP13** | 24 | 75 | 13 | 11 | 42 | 95 |
| **DPP3** | 8 | 3 | 4 | 0 | 82 | 99 |
| **DPP4** | 17 | 18 | 21 | 13 | 53 | 93 |
| **DPP7** | 20 | 29 | 83 | 8 | 30 | 77 |
| **DPP8** | 8 | 8 | 13 | 0 | 15 | 81 |
| **DPP9** | 18 | 13 | 14 | 0 | 49 | 87 |
| **FAP** | 30 | 25 | 29 | 1 | 35 | 100 |
| **HCV1a (D168V)** | 5 | 66 | 1 | 18 | 59 | 92 |
| **HCV1b** | 11 | 46 | 29 | 26 | 58 | 93 |
| **HCV1b (D168V)** | 5 | 44 | 13 | 30 | 78 | 98 |
| **HCV1b (R155K)** | 1 | 41 | 0 | 18 | 57 | 94 |
| **HCV1b (R155Q)** | 5 | 58 | 12 | 22 | 51 | 99 |
| **HCV2a** | 1 | 20 | 0 | 11 | 55 | 97 |
| **POP** | 31 | 29 | 27 | 7 | 29 | 100 |
| **APC** | 77 | 98 | 99 | 16 | 43 | 95 |

*Compound was fluorescent for most assays.

3.2. Results of the Effects of the Compounds on Individual Protease Activity
     3.2.1. BACE1

| Compounds | BACE1 Activity (Fluorescence count) | | Background  (Fluorescence count) | | % Activity | | % Inhibition |
| --- | --- | --- | --- | --- | --- | --- | --- |
|  | Repeat1 | Repeat2 | Repeat1 | Repeat2 | Repeat1 | Repeat2 |  |
| No Compound | 543 | 542 | 62 | 65 | 100 | 100 | 0 |
| NCGC00167526, 10 μM | 501 | 530 | 60 | 53 | 93 | 99 | 4 |
| NCGC00160398, 10 μM | 845 | 788 | 334 | 325 | 108 | 96 | 0 |
| NCGC00025297, 10 μM | 522 | 553 | 61 | 53 | 97 | 103 | 0 |
| Verubecestat, 0.003 μM | 501 | 517 | 58 | 54 | 93 | 96 | 6 |
| Verubecestat, 0.03 μM | 290 | 287 | 60 | 53 | 48 | 48 | 52 |
| Verubecestat, 0.3 μM | 86 | 84 | 59 | 53 | 5 | 5 | 95 |
| Background | 67 | 65 | 62 | 63 |  |  |  |

Table 3.2.1. Data for the Effect of the Compounds on BACE1 Activity

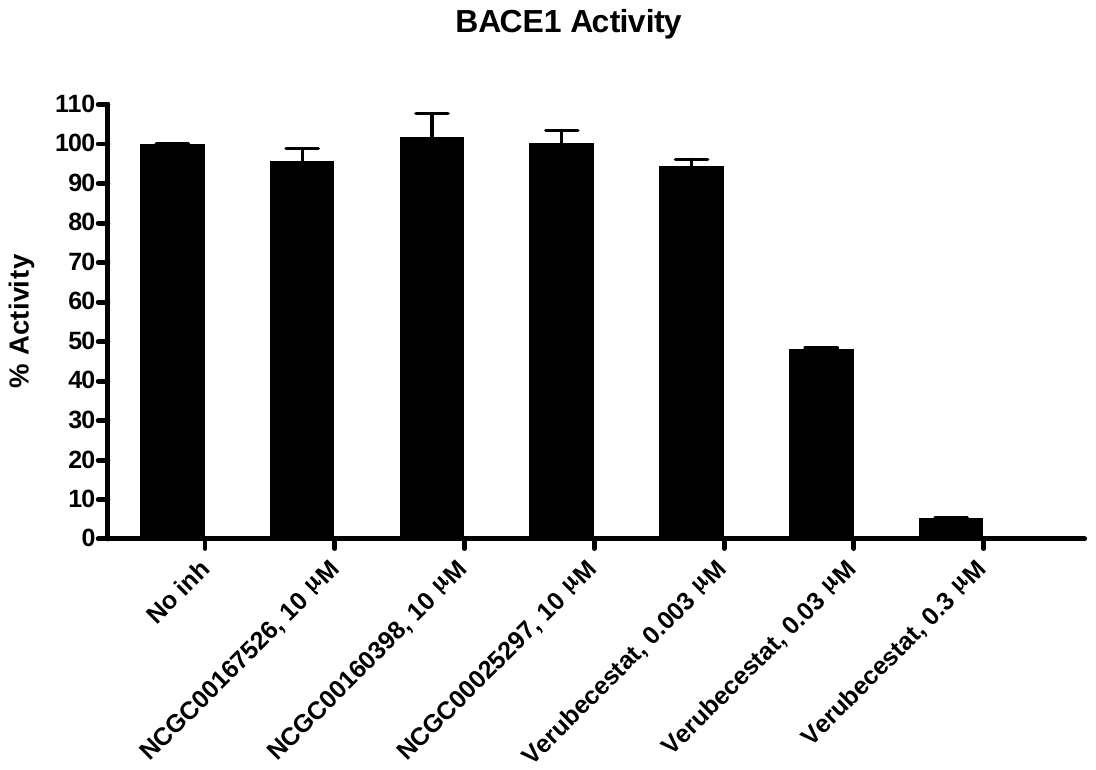

3.2.2. Caspase-3

| Compounds | Caspase-3 Activity (Fluorescence count) | | Background  (Fluorescence count) | | % Activity | | % Inhibition |
| --- | --- | --- | --- | --- | --- | --- | --- |
|  | Repeat1 | Repeat2 | Repeat1 | Repeat2 | Repeat1 | Repeat2 |  |
| No Compound | 733 | 724 | 62 | 56 | 100 | 100 | 0 |
| NCGC00167526, 10 μM | 728 | 732 | 61 | 59 | 100 | 101 | 0 |
| NCGC00160398, 10 μM | 1304 | 1244 | 636 | 559 | 100 | 102 | 0 |
| NCGC00025297, 10 μM | 715 | 725 | 69 | 60 | 96 | 99 | 2 |
| Ac-DNLD-CHO, 0.001 μM | 677 | 664 | 62 | 70 | 92 | 89 | 10 |
| Ac-DNLD-CHO, 0.01 μM | 566 | 568 | 58 | 65 | 76 | 75 | 25 |
| Ac-DNLD-CHO, 0.1 μM | 195 | 195 | 55 | 68 | 21 | 19 | 80 |
| Background | 60 | 64 | 58 | 62 |  |  |  |

Table 3.2.2. Data for the Effect of the Compounds on Caspase-3 Activity

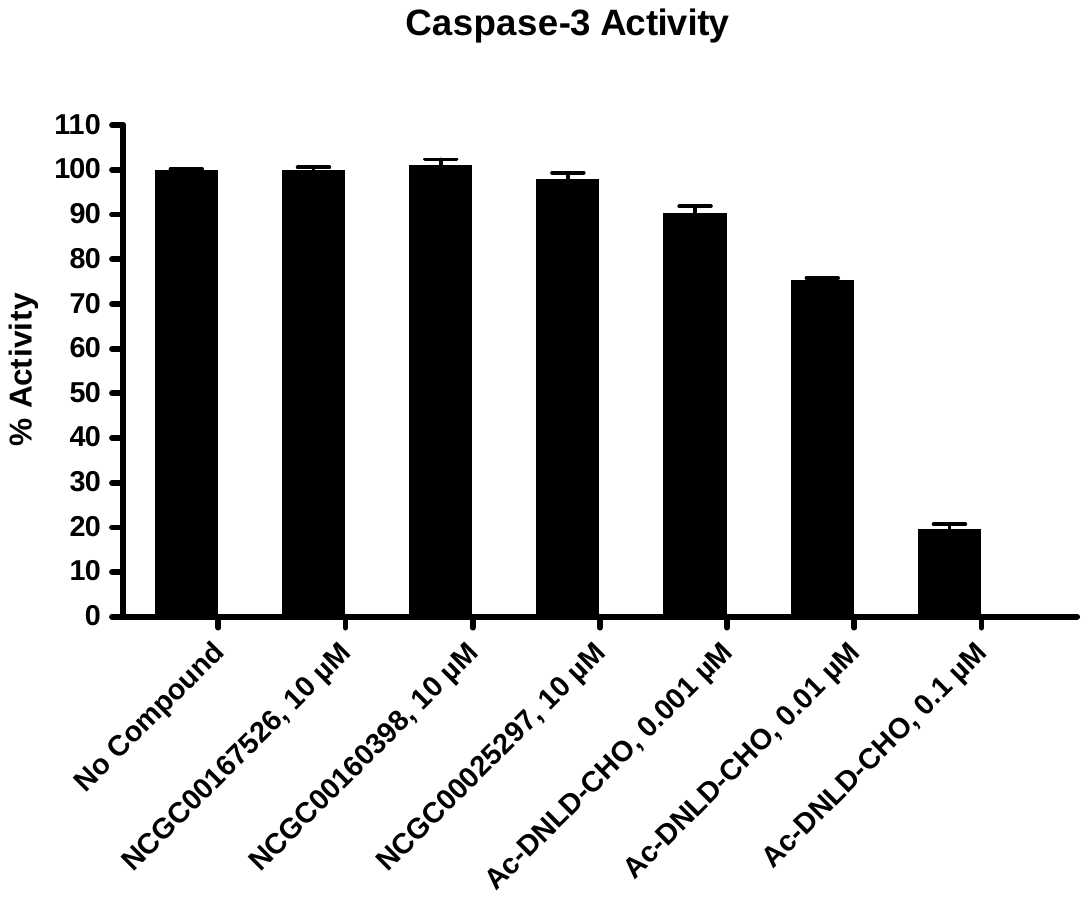

3.2.3. Caspase-6

| Compounds | Caspase-6 Activity (Fluorescence count) | | Background  (Fluorescence count) | | % Activity | | % Inhibition |
| --- | --- | --- | --- | --- | --- | --- | --- |
|  | Repeat1 | Repeat2 | Repeat1 | Repeat2 | Repeat1 | Repeat2 |  |
| No Compound | 491 | 477 | 26 | 20 | 101 | 99 | 0 |
| NCGC00167526, 10 μM | 474 | 478 | 32 | 25 | 96 | 98 | 3 |
| NCGC00160398, 10 μM | 1077 | 1059 | 629 | 626 | 97 | 94 | 4 |
| NCGC00025297, 10 μM | 485 | 487 | 27 | 24 | 99 | 100 | 0 |
| Ac-IETD-CHO, 0.05 μM | 453 | 461 | 33 | 28 | 91 | 94 | 8 |
| Ac-IETD-CHO, 0.5 μM | 333 | 328 | 22 | 31 | 67 | 64 | 34 |
| Ac-IETD-CHO, 5 μM | 61 | 62 | 28 | 25 | 7 | 8 | 93 |
| Background | 22 | 23 | 21 | 22 |  |  |  |

**Table 3.2.3. Data for the Effect of the Compounds on Caspase-6 Activity**

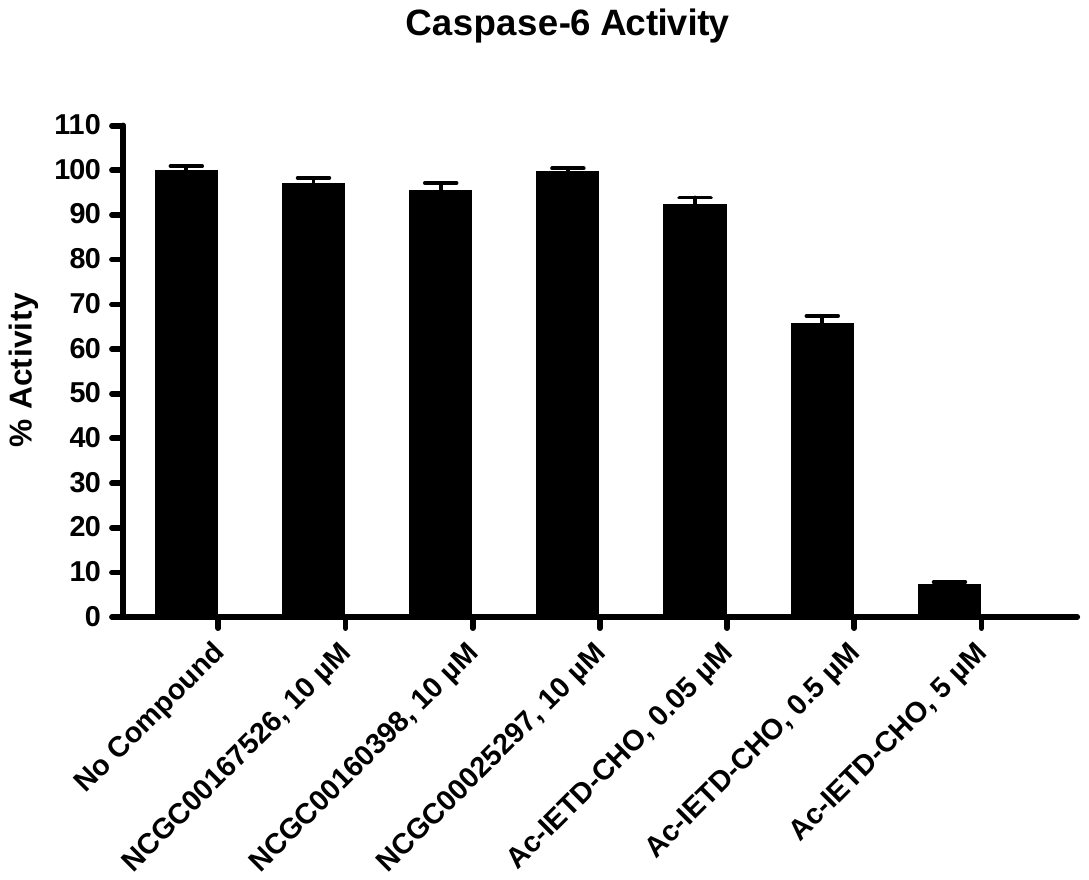

3.2.4. Caspase-7

| Compounds | Caspase-7 Activity (Fluorescence count) | | Background  (Fluorescence count) | | % Activity | | % Inhibition |
| --- | --- | --- | --- | --- | --- | --- | --- |
|  | Repeat1 | Repeat2 | Repeat1 | Repeat2 | Repeat1 | Repeat2 |  |
| No Compound | 731 | 702 | 62 | 57 | 102 | 98 | 0 |
| NCGC00167526, 10 μM | 718 | 731 | 59 | 64 | 100 | 102 | 0 |
| NCGC00160398, 10 μM | 1251 | 1257 | 596 | 614 | 100 | 98 | 1 |
| NCGC00025297, 10 μM | 700 | 713 | 55 | 59 | 98 | 100 | 1 |
| Ac-DNLD-CHO, 0.005 μM | 676 | 659 | 58 | 63 | 94 | 91 | 8 |
| Ac-DNLD-CHO, 0.05 μM | 433 | 439 | 60 | 54 | 57 | 58 | 43 |
| Ac-DNLD-CHO, 0.5 μM | 129 | 130 | 61 | 52 | 10 | 11 | 89 |
| Background | 57 | 58 | 53 | 56 |  |  |  |

**Table 3.2.4. Data for the Effect of the Compounds on Caspase-7 Activity**

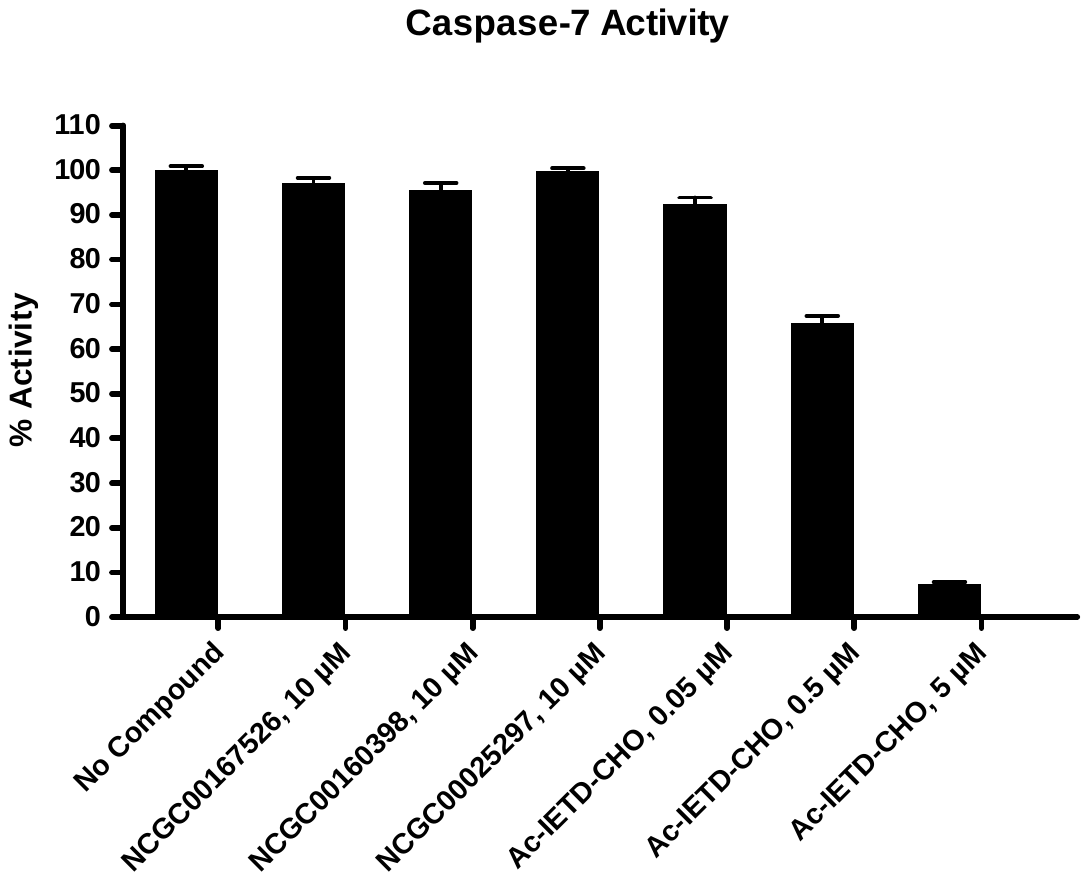

3.2.5. Caspase-8

| Compounds | Caspase-8 Activity (Fluorescence count) | | Background  (Fluorescence count) | | % Activity | | % Inhibition |
| --- | --- | --- | --- | --- | --- | --- | --- |
|  | Repeat1 | Repeat2 | Repeat1 | Repeat2 | Repeat1 | Repeat2 |  |
| No Compound | 434 | 420 | 32 | 25 | 101 | 99 | 0 |
| NCGC00167526, 10 μM | 430 | 425 | 26 | 26 | 101 | 100 | 0 |
| NCGC00160398, 10 μM | 1009 | 981 | 602 | 592 | 102 | 98 | 0 |
| NCGC00025297, 10 μM | 425 | 420 | 28 | 30 | 100 | 98 | 1 |
| Ac-IETD-CHO, 0.001 μM | 417 | 428 | 25 | 23 | 98 | 102 | 0 |
| Ac-IETD-CHO, 0.01 μM | 271 | 277 | 29 | 24 | 60 | 63 | 38 |
| Ac-IETD-CHO, 0.1 μM | 120 | 123 | 29 | 33 | 22 | 22 | 78 |
| Background | 32 | 31 | 30 | 28 |  |  |  |

**Table 3.2.5. Data for the Effect of the Compounds on Caspase-8 Activity**

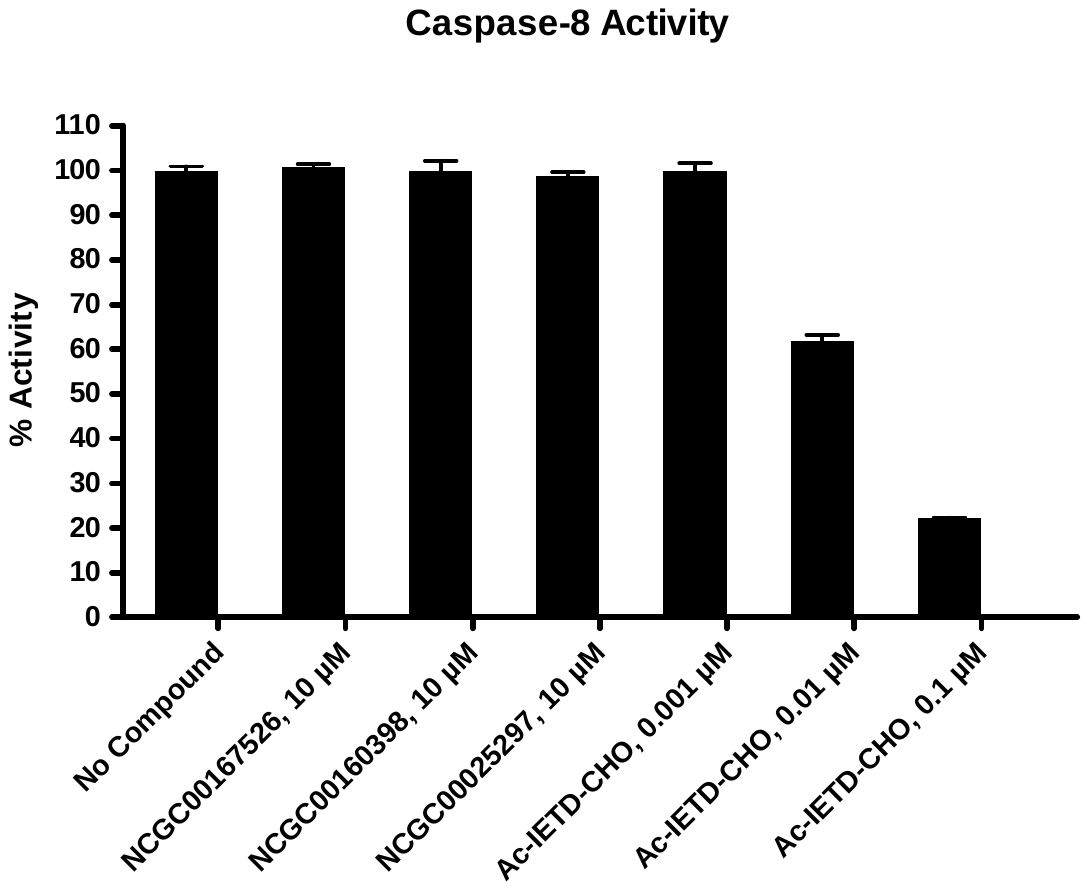

3.2.6. Caspase-9

| Compounds | Caspase-9 Activity (Fluorescence count) | | Background  (Fluorescence count) | | % Activity | | % Inhibition |
| --- | --- | --- | --- | --- | --- | --- | --- |
|  | Repeat1 | Repeat2 | Repeat1 | Repeat2 | Repeat1 | Repeat2 |  |
| No Compound | 6584 | 6534 | 2048 | 1656 | 96 | 104 | 0 |
| NCGC00167526, 10 μM | 7008 | 7517 | 1490 | 1561 | 118 | 127 | 0 |
| NCGC00160398, 10 μM | 16405 | 16946 | 5338 | 5404 | 238 | 248 | 0 |
| NCGC00025297, 10 μM | 7544 | 6002 | 3235 | 1528 | 91 | 95 | 7 |
| Z-LEHD-FMK, 0.1 μM | 6160 | 5736 | 1612 | 1527 | 97 | 89 | 7 |
| Z-LEHD-FMK, 1 μM | 3618 | 4616 | 1555 | 1864 | 43 | 58 | 50 |
| Z-LEHD-FMK, 10 μM | 2372 | 2308 | 1577 | 1578 | 15 | 14 | 86 |
| Background | 2115 | 2049 | 1968 | 1989 |  |  |  |

**Table 3.2.6. Data for the Effect of the Compounds on Caspase-9 Activity**

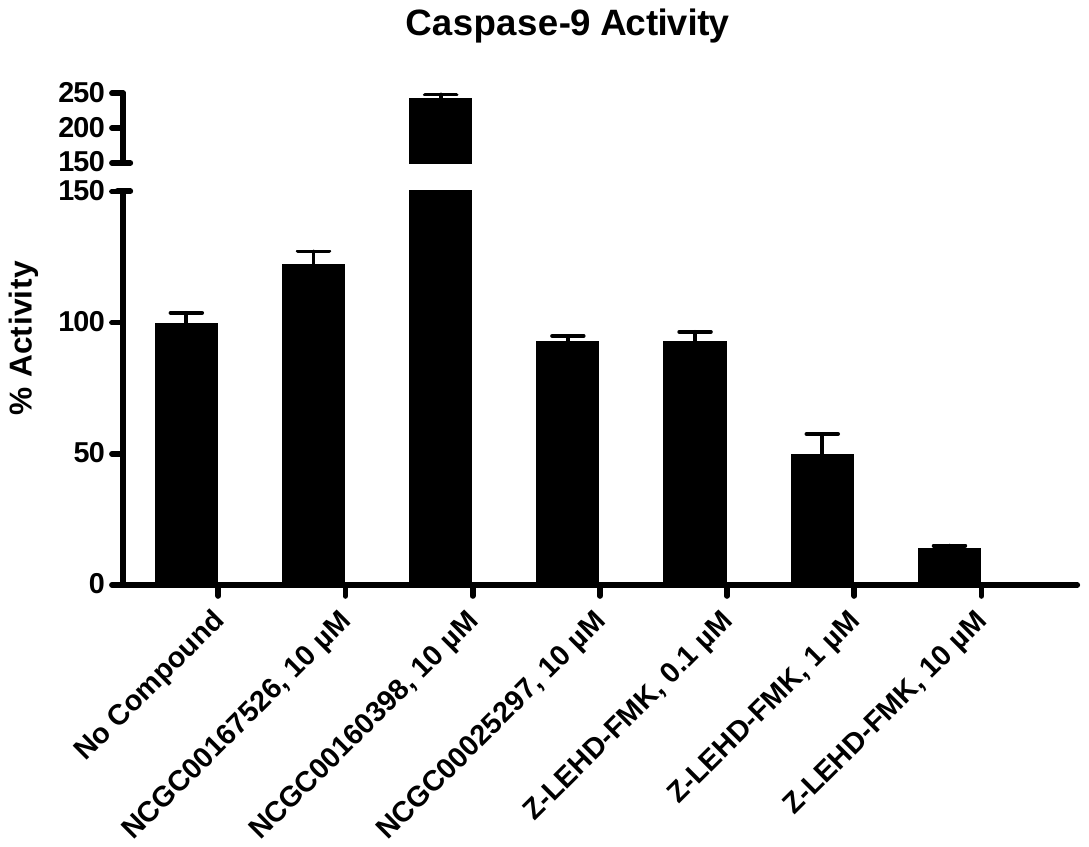

3.2.7. Renin

**Table 3.2.7. Data for the Effect of the Compounds on Renin Activity**

| Compounds | Renin Activity (Fluorescence count) | | % Activity | | % Inhibition |
| --- | --- | --- | --- | --- | --- |
|  | Repeat1 | Repeat2 | Repeat1 | Repeat2 |  |
| No Compound | 12253 | 12252 | 100 | 100 | 0 |
| NCGC00167526, 10 μM | 12115 | 12221 | 98 | 100 | 1 |
| NCGC00160398, 10 μM | 8395 | 8621 | 53 | 56 | 45 |
| NCGC00025297, 10 μM | 12390 | 12284 | 102 | 100 | 0 |
| Aliskiren, 0.0001 μM | 12335 | 12289 | 101 | 100 | 0 |
| Aliskiren, 0.001 μM | 11289 | 11463 | 88 | 90 | 11 |
| Aliskiren, 0.01 μM | 6377 | 6625 | 29 | 32 | 70 |
| Background | 3983 | 3999 |  |  |  |

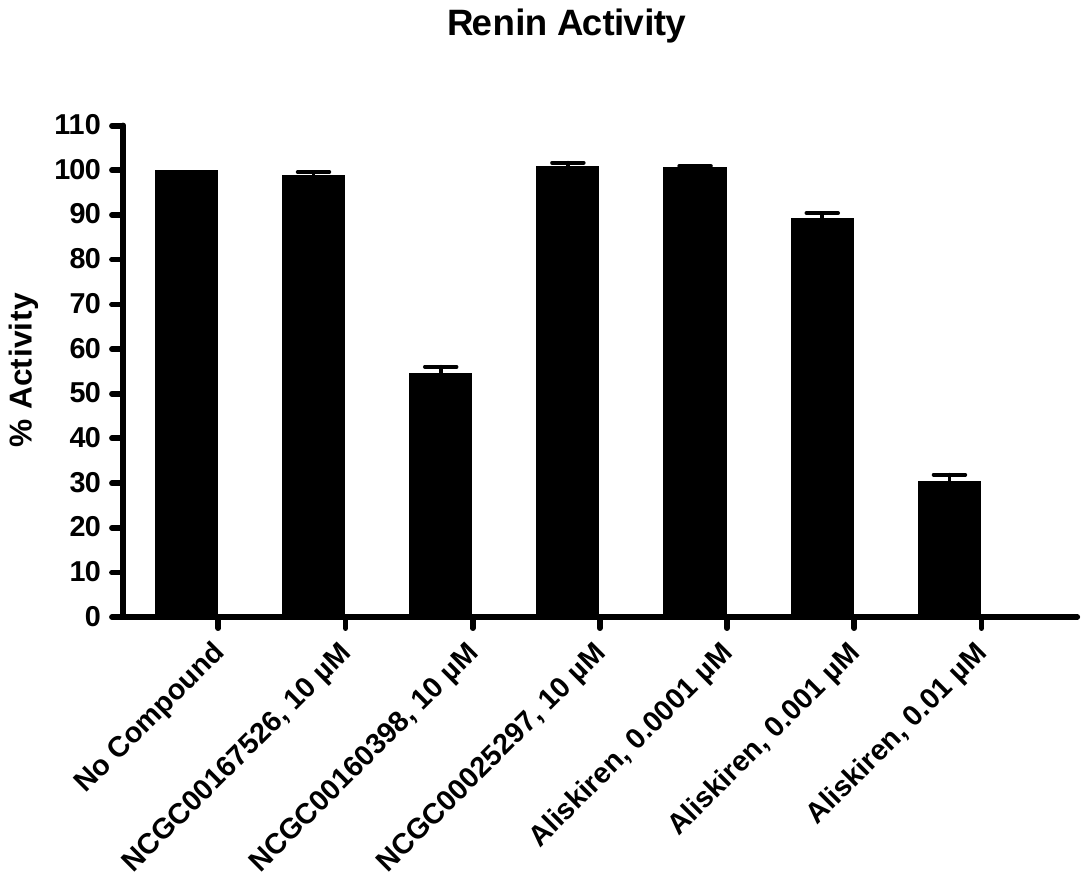

3.2.8. 3CL Protease

| Compounds | 3CL Proteae Activity (Fluorescence count) | | Background  (Fluorescence count) | | % Activity | | % Inhibition |
| --- | --- | --- | --- | --- | --- | --- | --- |
|  | Repeat1 | Repeat2 | Repeat1 | Repeat2 | Repeat1 | Repeat2 |  |
| No Compound | 1171 | 1226 | 53 | 54 | 97 | 103 | 0 |
| NCGC00167526, 10 μM | 1123 | 1150 | 47 | 53 | 93 | 95 | 6 |
| NCGC00160398, 10 μM | 1198 | 1219 | 189 | 191 | 86 | 88 | 13 |
| NCGC00025297, 10 μM | 1171 | 1172 | 51 | 51 | 97 | 97 | 3 |
| GC376, 0.05 μM | 1125 | 1171 | 73 | 150 | 90 | 87 | 11 |
| GC376, 0.5 μM | 676 | 605 | 79 | 84 | 42 | 34 | 62 |
| GC376, 5 μM | 310 | 300 | 77 | 56 | 4 | 5 | 96 |
| Background | 259 | 242 | 54 | 55 |  |  |  |

**Table 3.2.8. Data for the Effect of the Compounds on 3CL Protease Activity**

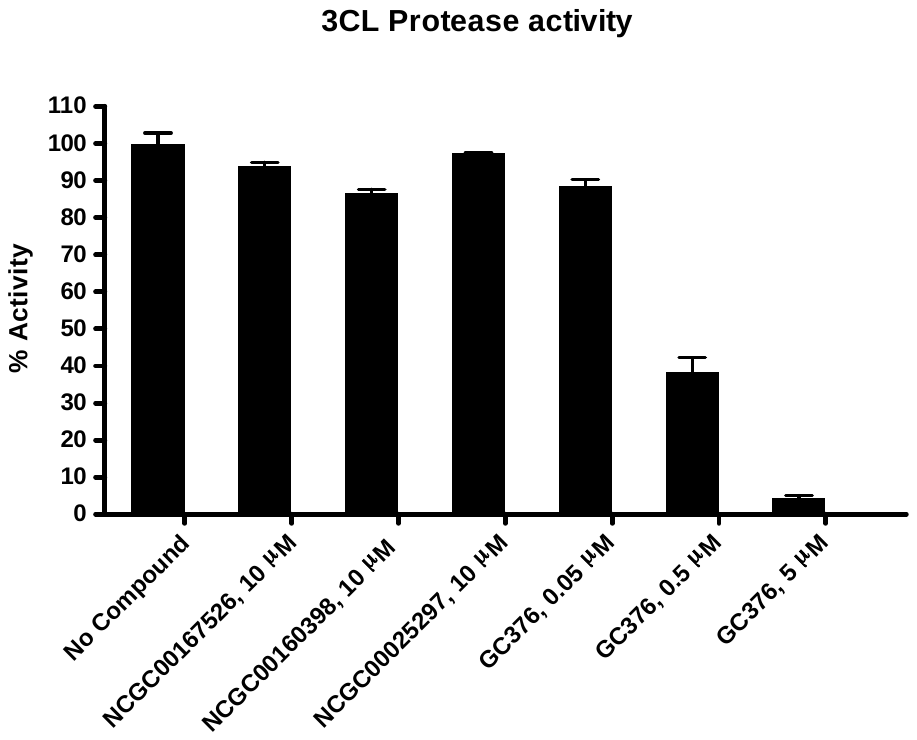

3.2.9. A20

| Compounds | A20 Activity (Fluorescence count) | | Background  (Fluorescence count) | | % Activity | | % Inhibition |
| --- | --- | --- | --- | --- | --- | --- | --- |
|  | Repeat1 | Repeat2 | Repeat1 | Repeat2 | Repeat1 | Repeat2 |  |
| No Compound | 192 | 189 | 22 | 22 | 101 | 99 | 0 |
| NCGC00167526, 10 μM | 183 | 183 | 22 | 21 | 95 | 96 | 4 |
| NCGC00160398, 10 μM | 221 | 221 | 75 | 74 | 86 | 87 | 13 |
| NCGC00025297, 10 μM | 183 | 183 | 21 | 20 | 96 | 97 | 4 |
| Ub-Aldehyde, 0.001 μM | 170 | 172 | 19 | 22 | 89 | 89 | 11 |
| Ub-Aldehyde, 0.01 μM | 101 | 99 | 21 | 21 | 47 | 45 | 54 |
| Ub-Aldehyde, 0.1 μM | 28 | 28 | 20 | 19 | 3 | 4 | 96 |
| Background | 15 | 15 | 12 | 13 |  |  |  |

**Table 3.2.9. Data for the Effect of the Compounds on A20 Activity**

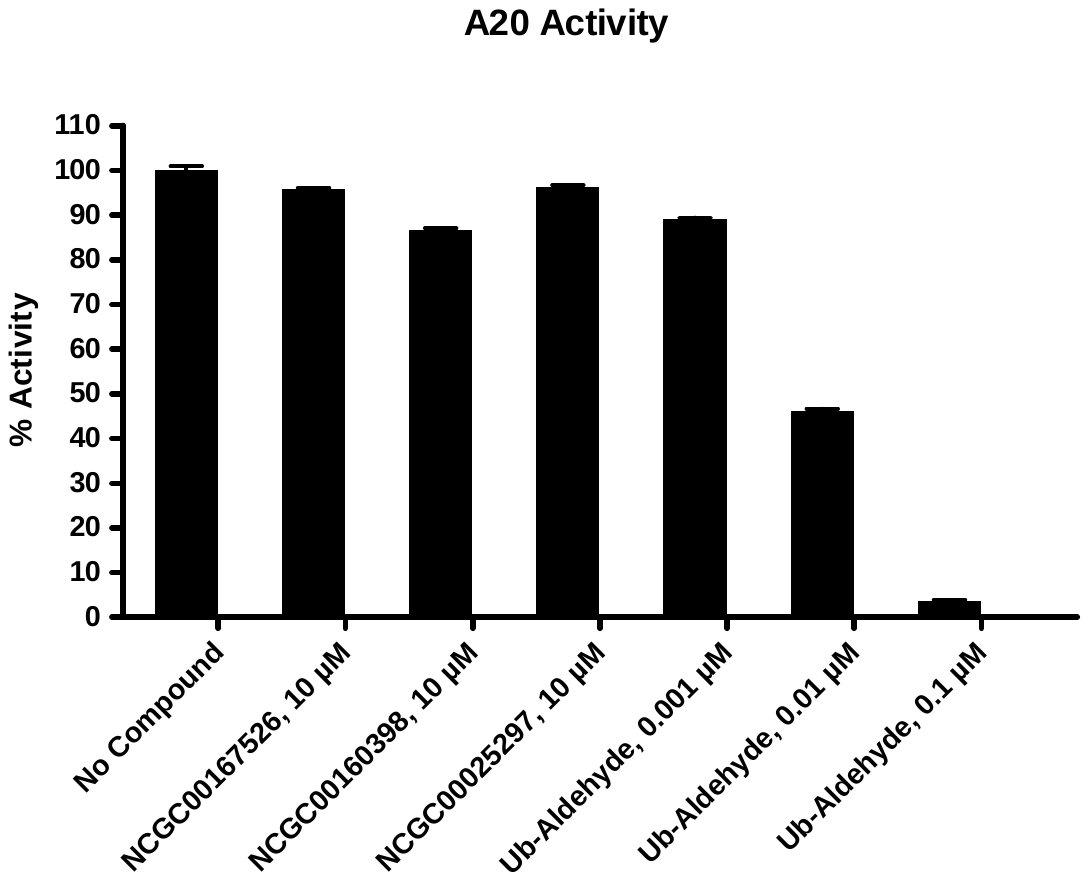

3.2.10. Ataxin-3

| Compounds | Ataxin-3 Activity (Fluorescence count) | | Background  (Fluorescence count) | | % Activity | | % Inhibition |
| --- | --- | --- | --- | --- | --- | --- | --- |
|  | Repeat1 | Repeat2 | Repeat1 | Repeat2 | Repeat1 | Repeat2 |  |
| No Compound | 93 | 92 | 29 | 30 | 102 | 98 | 0 |
| NCGC00167526, 10 μM | 90 | 93 | 30 | 31 | 95 | 98 | 3 |
| NCGC00160398, 10 μM | 212 | 214 | 68 | 67 | 230 | 234 | 0 |
| NCGC00025297, 10 μM | 91 | 88 | 27 | 30 | 102 | 92 | 3 |
| Ub-Aldehyde, 0.1 μM | 73 | 73 | 28 | 30 | 71 | 68 | 30 |
| Ub-Aldehyde, 1 μM | 49 | 50 | 29 | 28 | 31 | 34 | 67 |
| Ub-Aldehyde, 10 μM | 26 | 26 | 29 | 29 | 4 | 4 | 96 |
| Background | 15 | 16 | 16 | 16 |  |  |  |

**Table 3.2.10. Data for the Effect of the Compounds on Ataxin-3 Activity**

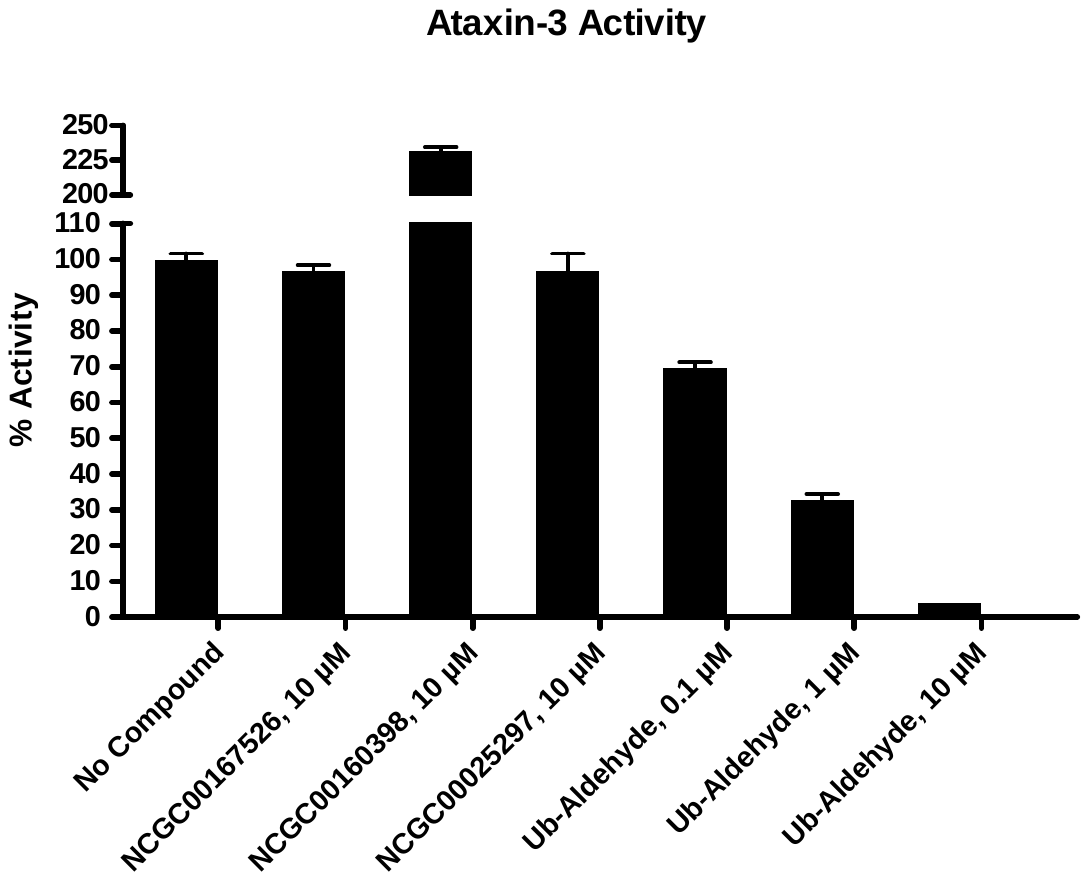

3.2.11. Cathepsin B

**Table 3.2.11. Data for the Effect of the Compounds on Cathepsin B Activity**

| Compounds | Cathepsin B Activity (Fluorescence count) | | % Activity | | % Inhibition |
| --- | --- | --- | --- | --- | --- |
|  | Repeat1 | Repeat2 | Repeat1 | Repeat2 |  |
| No Compound | 1818 | 2012 | 95 | 105 | 0 |
| NCGC00167526, 10 μM | 2032 | 2004 | 106 | 105 | 0 |
| NCGC00160398, 10 μM | 994 | 1056 | 50 | 53 | 48 |
| NCGC00025297, 10 μM | 1948 | 2033 | 102 | 106 | 0 |
| E-64, 0.0005 μM | 1274 | 1388 | 65 | 71 | 32 |
| E-64, 0.005 μM | 479 | 507 | 22 | 24 | 77 |
| E-64, 0.05 μM | 119 | 121 | 3 | 3 | 97 |
| Background | 64 | 72 |  |  |  |

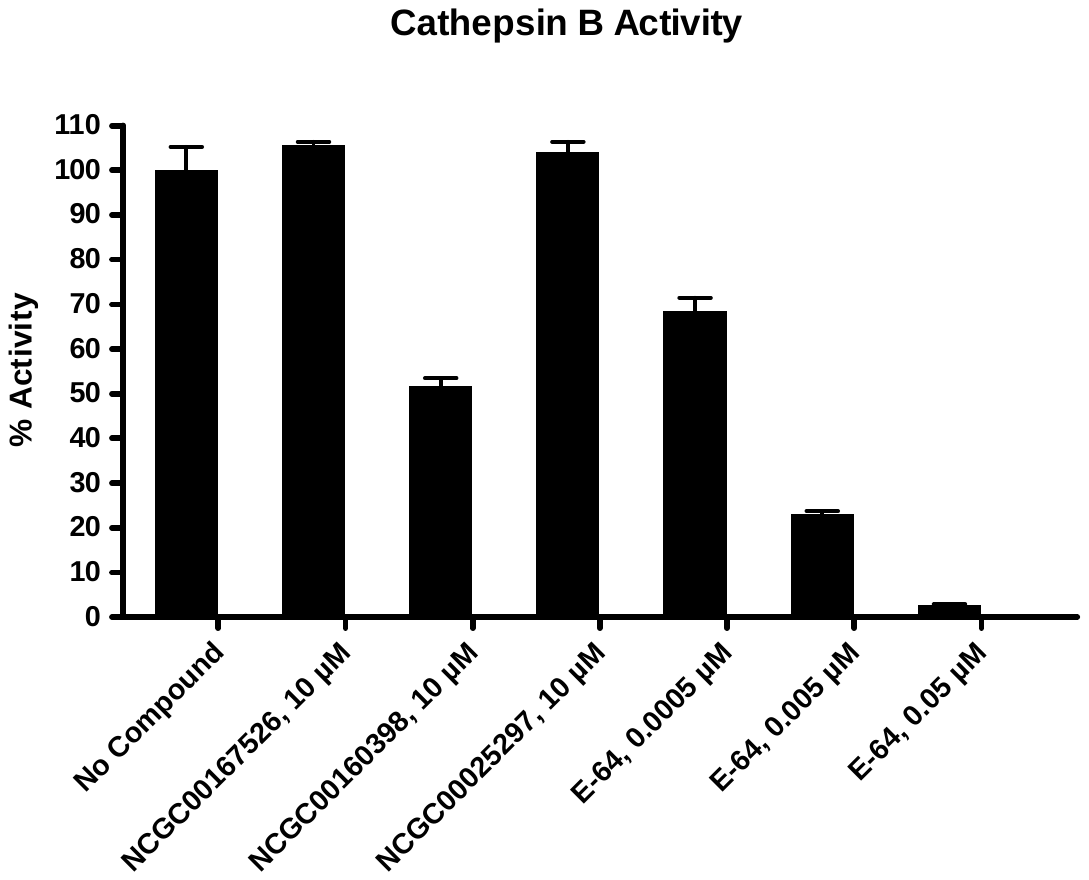

3.2.12. Cathepsin F

**Table 3.2.12. Data for the Effect of the Compounds on Cathepsin F Activity**

| Compounds | Cathepsin F Activity (Fluorescence count) | | % Activity | | % Inhibition |
| --- | --- | --- | --- | --- | --- |
|  | Repeat1 | Repeat2 | Repeat1 | Repeat2 |  |
| No Compound | 857 | 852 | 100 | 100 | 0 |
| NCGC00167526, 10 μM | 870 | 858 | 102 | 101 | 0 |
| NCGC00160398, 10 μM | 855 | 868 | 100 | 102 | 0 |
| NCGC00025297, 10 μM | 843 | 839 | 98 | 98 | 2 |
| Cystatin C, 0.1 μM | 724 | 705 | 81 | 78 | 20 |
| Cystatin C, 1 μM | 370 | 320 | 30 | 23 | 73 |
| Cystatin C, 10 μM | 232 | 247 | 10 | 12 | 89 |
| Background | 164 | 158 |  |  |  |

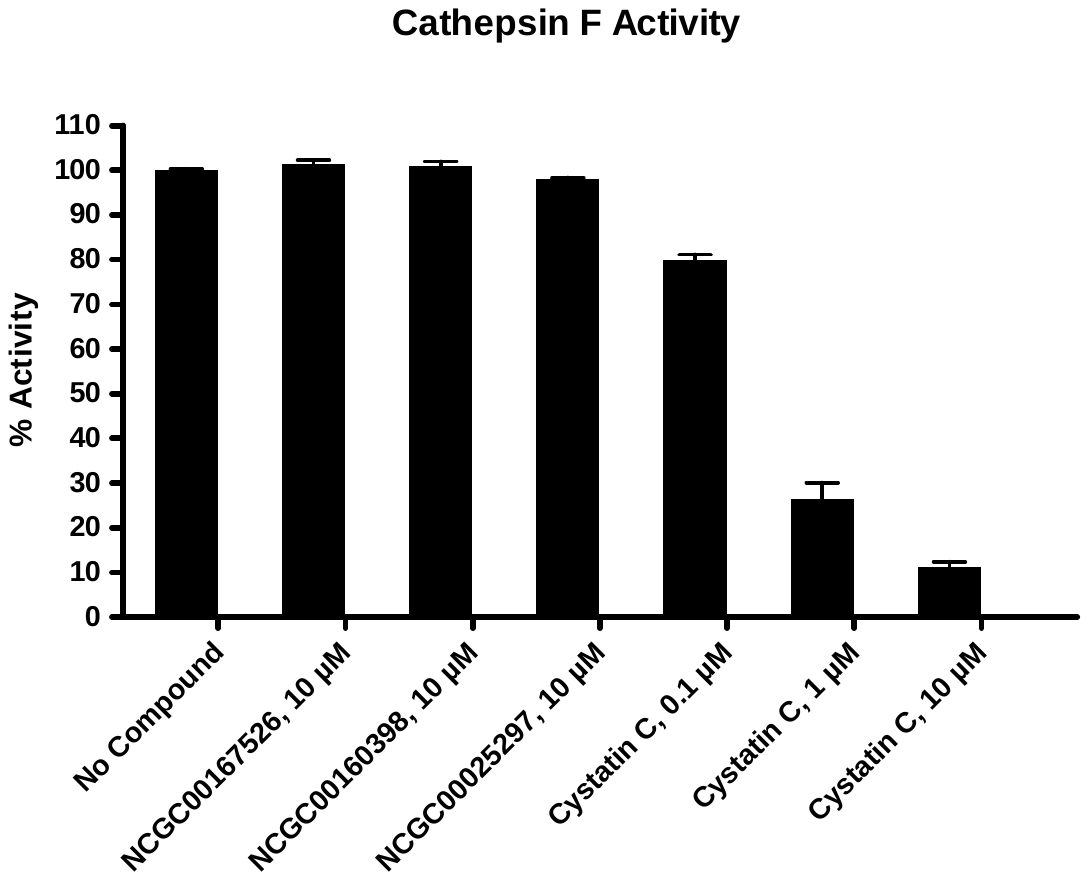

3.2.13. Cathepsin L

**Table 3.2.13. Data for the Effect of the Compounds on Cathepsin L Activity**

| Compounds | Cathepsin L Activity (Fluorescence count) | | % Activity | | % Inhibition |
| --- | --- | --- | --- | --- | --- |
|  | Repeat1 | Repeat2 | Repeat1 | Repeat2 |  |
| No Compound | 3581 | 3625 | 99 | 101 | 0 |
| NCGC00167526, 10 μM | 3571 | 3486 | 99 | 97 | 2 |
| NCGC00160398, 10 μM | 1563 | 1606 | 42 | 44 | 57 |
| NCGC00025297, 10 μM | 3693 | 3583 | 103 | 99 | 0 |
| E-64, 0.0005 μM | 3128 | 3141 | 87 | 87 | 13 |
| E-64, 0.005 μM | 2209 | 2271 | 61 | 62 | 39 |
| E-64, 0.05 μM | 629 | 663 | 16 | 17 | 84 |
| Background | 64 | 69 |  |  |  |

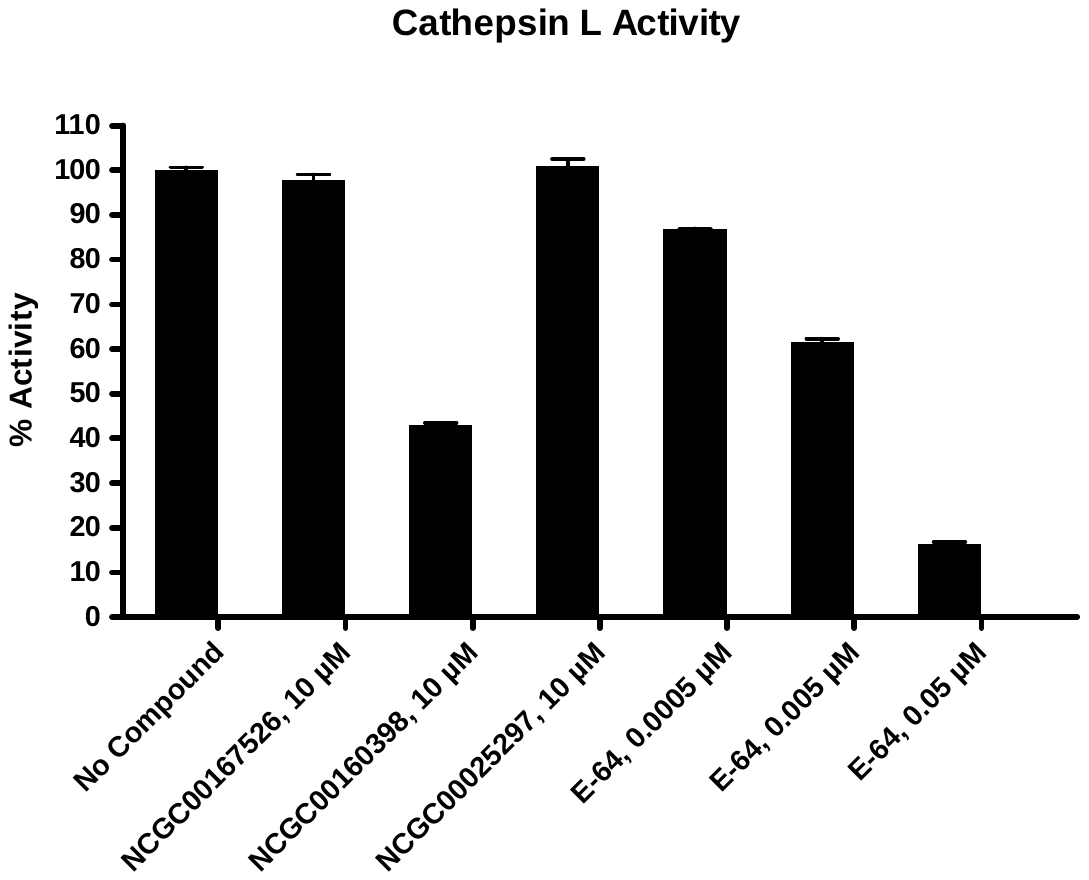

3.2.14. Cathepsin S

**Table 3.2.14. Data for the Effect of the Compounds on Cathepsin S Activity**

| Compounds | Cathepsin S Activity (Fluorescence count) | | % Activity | | % Inhibition |
| --- | --- | --- | --- | --- | --- |
|  | Repeat1 | Repeat2 | Repeat1 | Repeat2 |  |
| No Compound | 15416 | 15602 | 99 | 101 | 0 |
| NCGC00167526, 10 μM | 15033 | 15041 | 97 | 97 | 3 |
| NCGC00160398, 10 μM | 14129 | 14453 | 91 | 93 | 8 |
| NCGC00025297, 10 μM | 14932 | 14929 | 96 | 96 | 4 |
| E-64, 0.005 μM | 12846 | 12674 | 83 | 82 | 18 |
| E-64, 0.05 μM | 3066 | 3005 | 19 | 19 | 81 |
| E-64, 0.5 μM | 515 | 575 | 3 | 3 | 97 |
| Background | 64 | 70 |  |  |  |

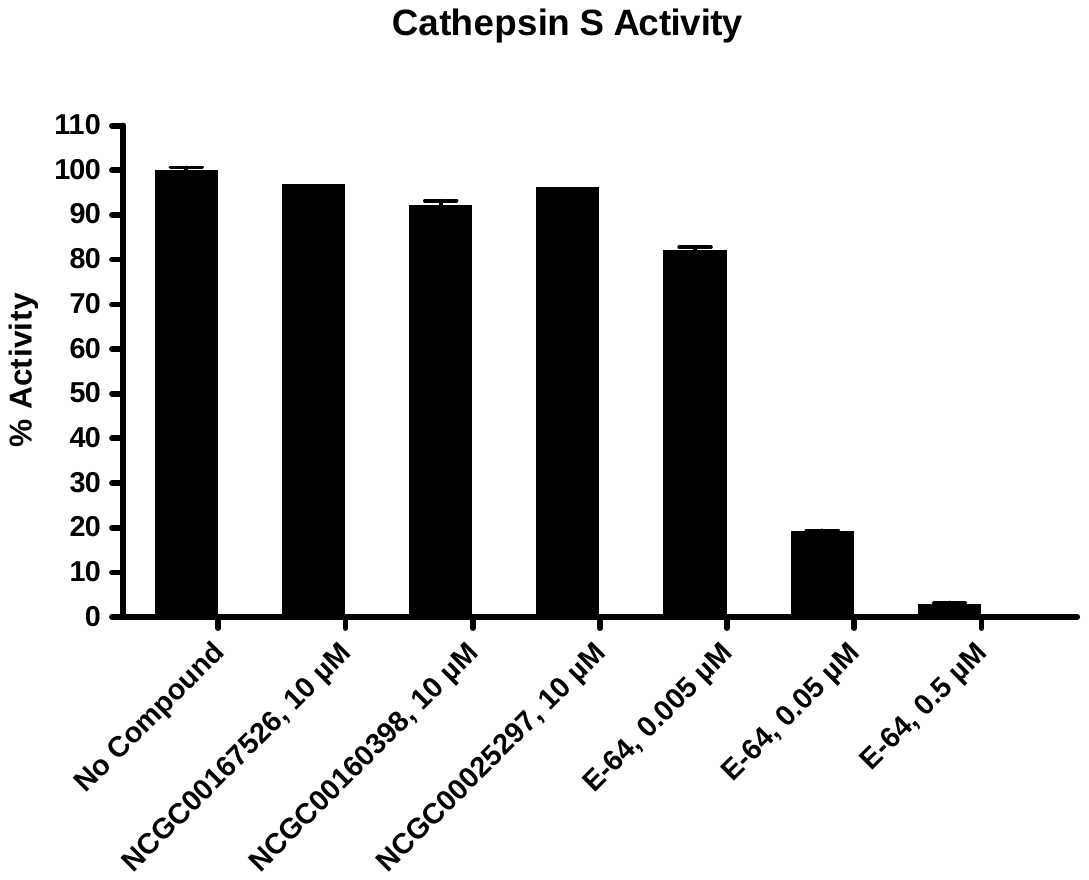

3.2.15. Cathepsin V

**Table 3.2.15. Data for the Effect of the Compounds on Cathepsin V Activity**

| Compounds | Cathepsin V Activity (Fluorescence count) | | % Activity | | % Inhibition |
| --- | --- | --- | --- | --- | --- |
|  | Repeat1 | Repeat2 | Repeat1 | Repeat2 |  |
| No Compound | 6640 | 6458 | 101 | 99 | 0 |
| NCGC00167526, 10 μM | 6463 | 6533 | 99 | 100 | 1 |
| NCGC00160398, 10 μM | 4748 | 4883 | 72 | 74 | 27 |
| NCGC00025297, 10 μM | 6423 | 6337 | 98 | 97 | 3 |
| E-64, 0.005 μM | 5488 | 5430 | 84 | 83 | 17 |
| E-64, 0.05 μM | 2947 | 2798 | 45 | 42 | 57 |
| E-64, 0.5 μM | 518 | 407 | 7 | 6 | 94 |
| Background | 45 | 43 |  |  |  |

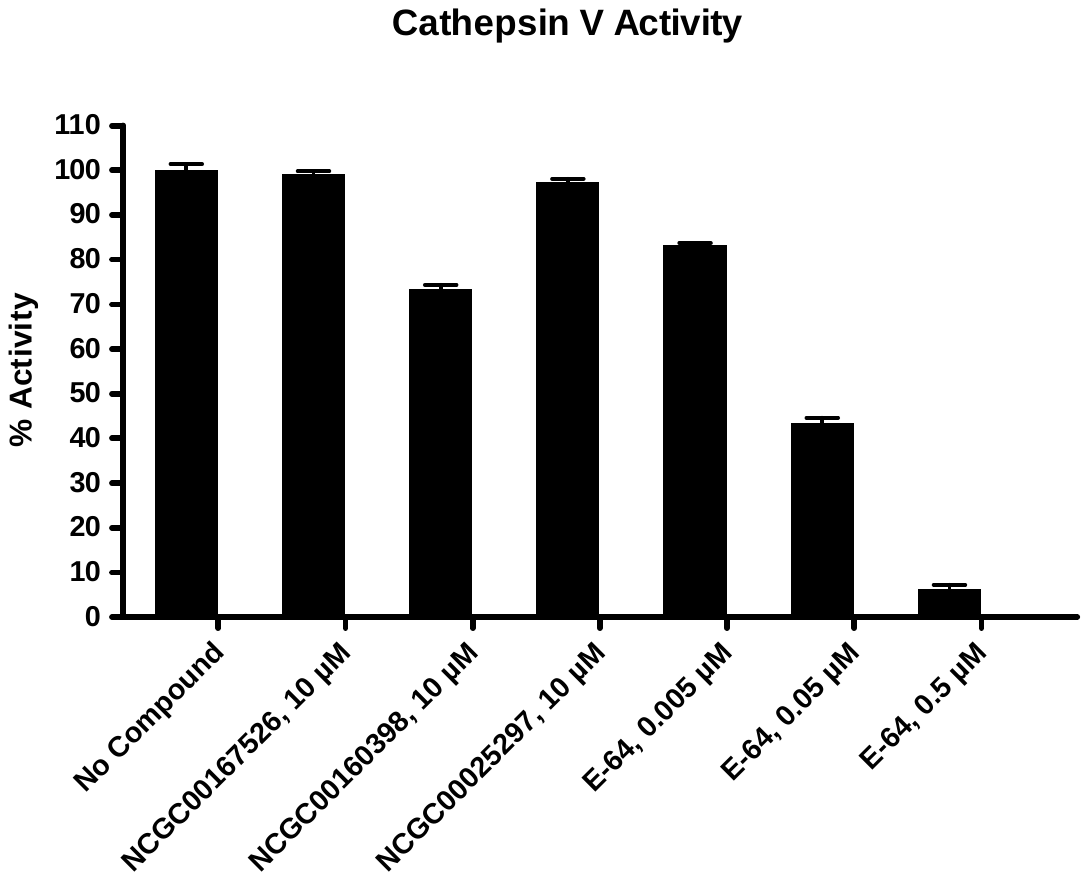

3.2.16. MALT1

| Compounds | MALT1 Activity (Fluorescence count) | | Background  (Fluorescence count) | | % Activity | | % Inhibition |
| --- | --- | --- | --- | --- | --- | --- | --- |
|  | Repeat1 | Repeat2 | Repeat1 | Repeat2 | Repeat1 | Repeat2 |  |
| No Compound | 7256 | 7345 | 450 | 544 | 100 | 100 | 0 |
| NCGC00167526, 10 μM | 3957 | 3934 | 361 | 325 | 52 | 52 | 48 |
| NCGC00160398, 10 μM | 4041 | 4049 | 500 | 546 | 51 | 50 | 49 |
| NCGC00025297, 10 μM | 4120 | 4121 | 382 | 348 | 54 | 55 | 46 |
| Z-VRPR-FMK, 0.001 μM | 6564 | 6756 | 400 | 514 | 90 | 92 | 9 |
| Z-VRPR-FMK, 0.01 μM | 3396 | 3379 | 295 | 344 | 44 | 43 | 56 |
| Z-VRPR-FMK, 0.1 μM | 668 | 669 | 278 | 369 | 4 | 2 | 97 |
| Background | 404 | 405 | 264 | 272 |  |  |  |

**Table 3.2.16. Data for the Effect of the Compounds on MALT1 Activity**

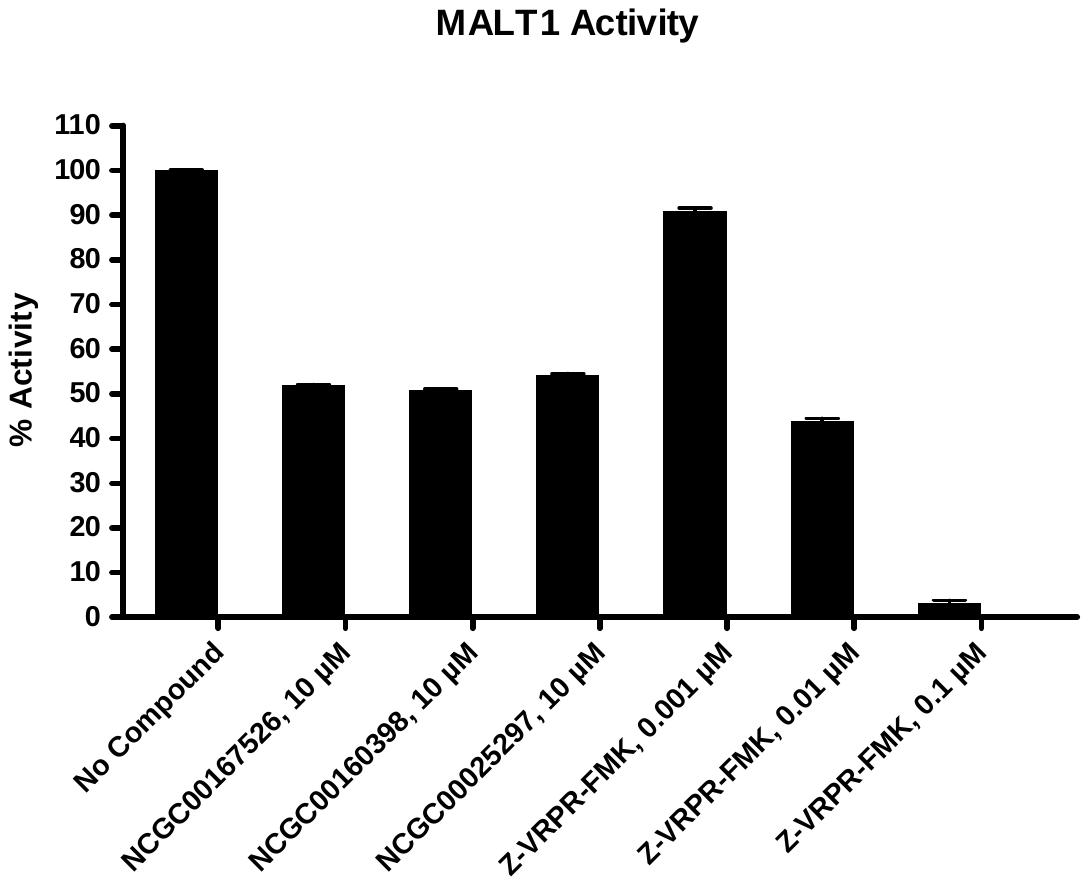

3.2.17. OTUD6Bc

| Compounds | OTUD6Bc Activity (Fluorescence count) | | Background  (Fluorescence count) | | % Activity | | % Inhibition |
| --- | --- | --- | --- | --- | --- | --- | --- |
|  | Repeat1 | Repeat2 | Repeat1 | Repeat2 | Repeat1 | Repeat2 |  |
| No Compound | 224 | 225 | 22 | 22 | 100 | 100 | 0 |
| NCGC00167526, 10 μM | 222 | 220 | 21 | 20 | 99 | 99 | 1 |
| NCGC00160398, 10 μM | 265 | 260 | 80 | 77 | 91 | 90 | 9 |
| NCGC00025297, 10 μM | 218 | 217 | 20 | 21 | 98 | 97 | 3 |
| Ub-Aldehyde, 0.01 μM | 218 | 205 | 22 | 21 | 97 | 91 | 6 |
| Ub-Aldehyde, 0.1 μM | 176 | 176 | 20 | 21 | 77 | 76 | 24 |
| Ub-Aldehyde, 1 μM | 17 | 18 | 16 | 18 | 0 | 0 | 100 |
| Background | 16 | 16 | 14 | 13 |  |  |  |

**Table 3.2.17. Data for the Effect of the Compounds on OTUD6Bc Activity**

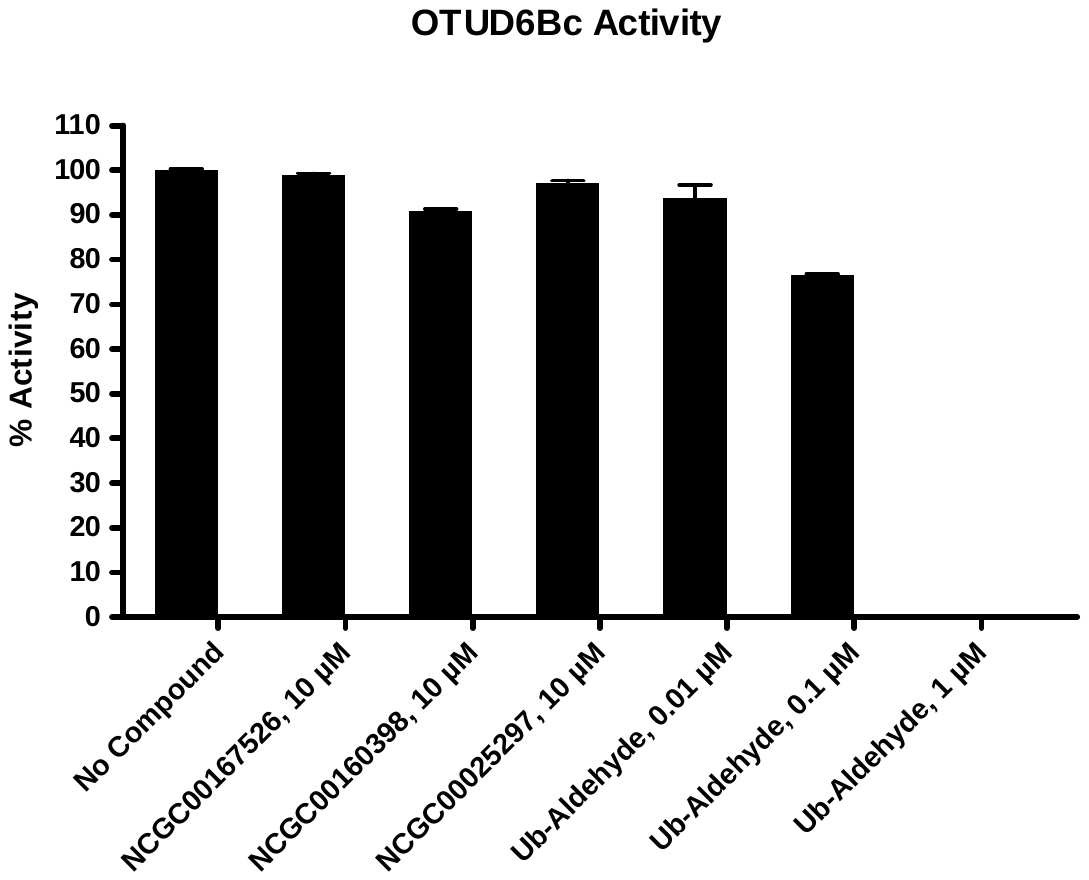

3.2.18. UCHL1

| Compounds | OCHL1 Activity (Fluorescence count) | | Background  (Fluorescence count) | | % Activity | | % Inhibition |
| --- | --- | --- | --- | --- | --- | --- | --- |
|  | Repeat1 | Repeat2 | Repeat1 | Repeat2 | Repeat1 | Repeat2 |  |
| No Compound | 230 | 230 | 23 | 23 | 100 | 100 | 0 |
| NCGC00167526, 10 μM | 214 | 217 | 22 | 23 | 93 | 94 | 7 |
| NCGC00160398, 10 μM | 296 | 295 | 64 | 62 | 112 | 113 | 0 |
| NCGC00025297, 10 μM | 218 | 219 | 23 | 22 | 94 | 95 | 5 |
| Ub-Aldehyde, 0.001 μM | 213 | 217 | 21 | 22 | 93 | 94 | 7 |
| Ub-Aldehyde, 0.01 μM | 187 | 185 | 21 | 20 | 80 | 80 | 20 |
| Ub-Aldehyde, 0.1 μM | 80 | 81 | 21 | 21 | 29 | 29 | 71 |
| Background | 14 | 14 | 14 | 14 |  |  |  |

**Table 3.2.18. Data for the Effect of the Compounds on UCHL1 Activity**

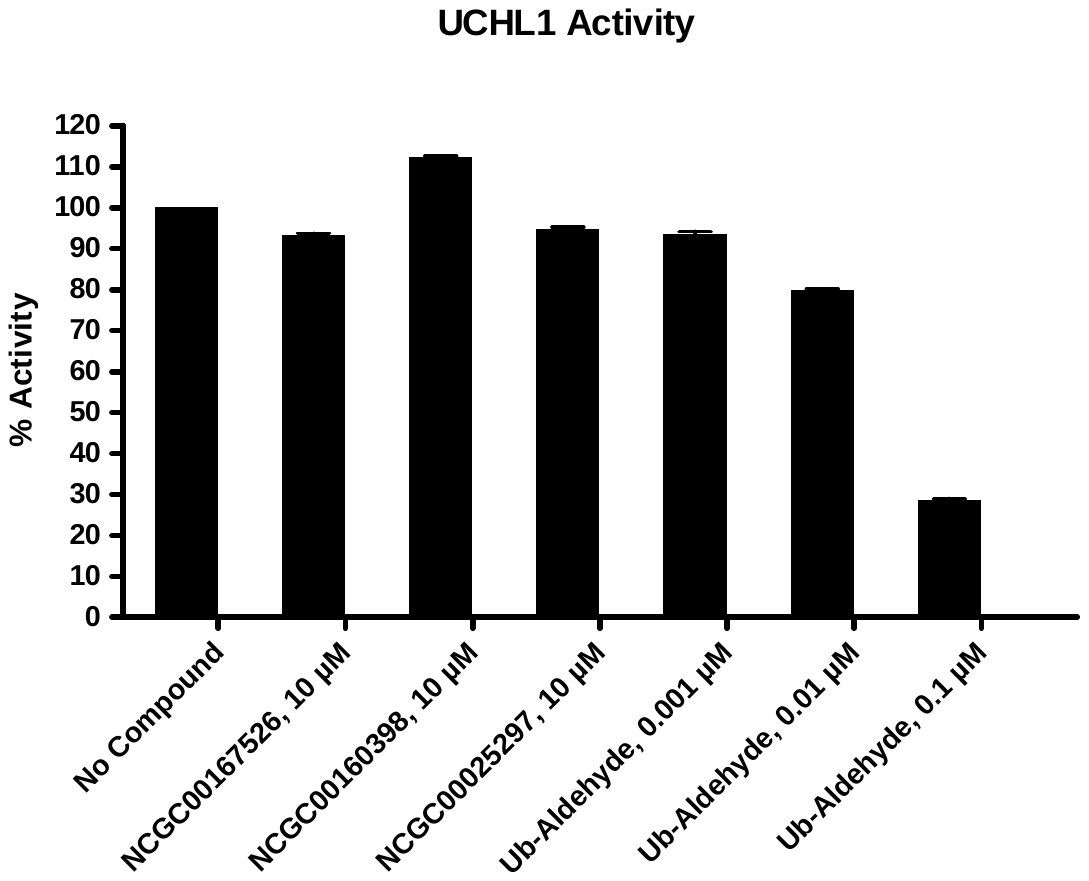

3.2.19. UCHL3

| Compounds | OCHL3 Activity (Fluorescence count) | | Background  (Fluorescence count) | | % Activity | | % Inhibition |
| --- | --- | --- | --- | --- | --- | --- | --- |
|  | Repeat1 | Repeat2 | Repeat1 | Repeat2 | Repeat1 | Repeat2 |  |
| No Compound | 634 | 626 | 49 | 48 | 101 | 99 | 0 |
| NCGC00167526, 10 μM | 615 | 611 | 48 | 46 | 97 | 97 | 3 |
| NCGC00160398, 10 μM | 647 | 647 | 98 | 97 | 94 | 95 | 6 |
| NCGC00025297, 10 μM | 607 | 610 | 48 | 49 | 96 | 96 | 4 |
| Ub-Aldehyde, 0.0001 μM | 600 | 608 | 44 | 46 | 96 | 97 | 4 |
| Ub-Aldehyde, 0.001 μM | 440 | 440 | 46 | 49 | 68 | 67 | 33 |
| Ub-Aldehyde, 0.01 μM | 111 | 112 | 40 | 42 | 12 | 11 | 89 |
| Background | 19 | 18 | 14 | 14 |  |  |  |

**Table 3.2.19. Data for the Effect of the Compounds on UCHL3 Activity**

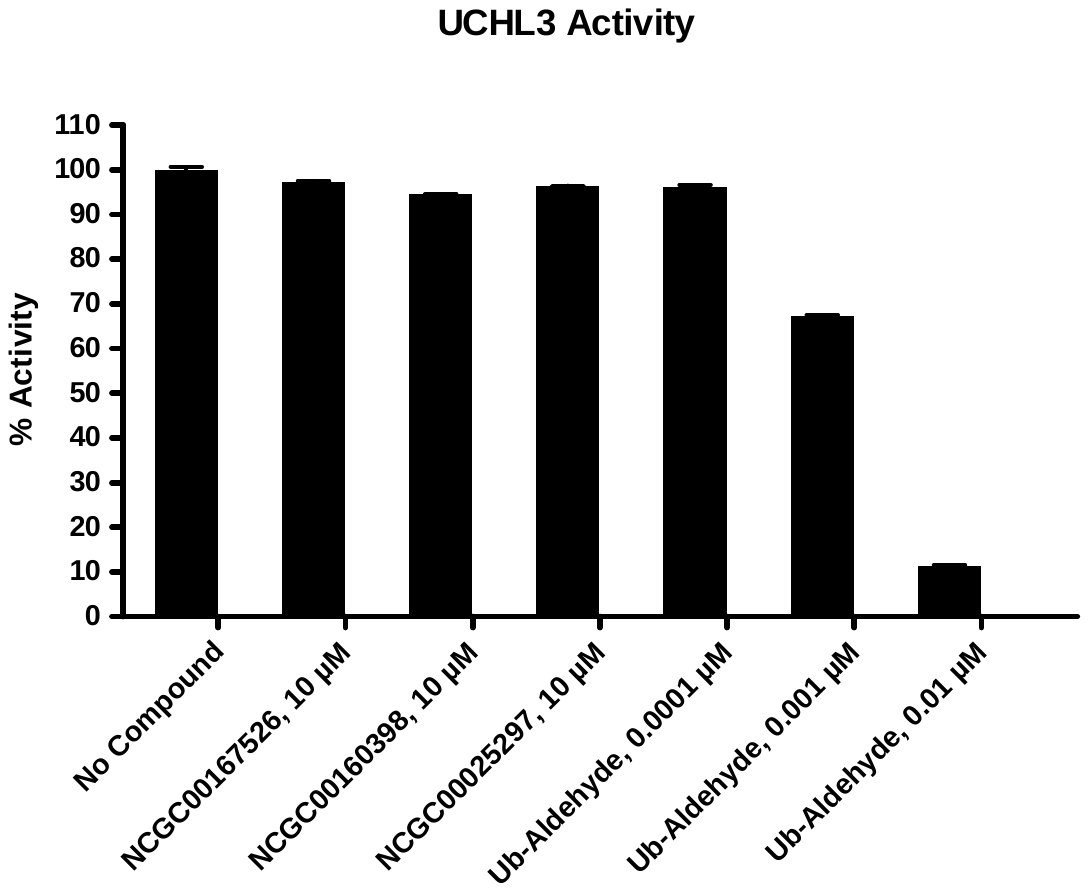

3.2.20. USP2

| Compounds | USP2 Activity (Fluorescence count) | | Background  (Fluorescence count) | | % Activity | | % Inhibition |
| --- | --- | --- | --- | --- | --- | --- | --- |
|  | Repeat1 | Repeat2 | Repeat1 | Repeat2 | Repeat1 | Repeat2 |  |
| No Compound | 343 | 330 | 23 | 21 | 102 | 98 | 0 |
| NCGC00167526, 10 μM | 332 | 337 | 22 | 23 | 99 | 100 | 1 |
| NCGC00160398, 10 μM | 404 | 400 | 69 | 67 | 107 | 106 | 0 |
| NCGC00025297, 10 μM | 331 | 333 | 23 | 22 | 98 | 99 | 2 |
| Ub-Aldehyde, 0.01 μM | 324 | 325 | 24 | 27 | 95 | 95 | 5 |
| Ub-Aldehyde, 0.1 μM | 163 | 166 | 23 | 23 | 44 | 45 | 55 |
| Ub-Aldehyde, 1 μM | 32 | 33 | 20 | 21 | 4 | 4 | 96 |
| Background | 15 | 14 | 14 | 13 |  |  |  |

**Table 3.2.20. Data for the Effect of the Compounds on USP2 Activity**

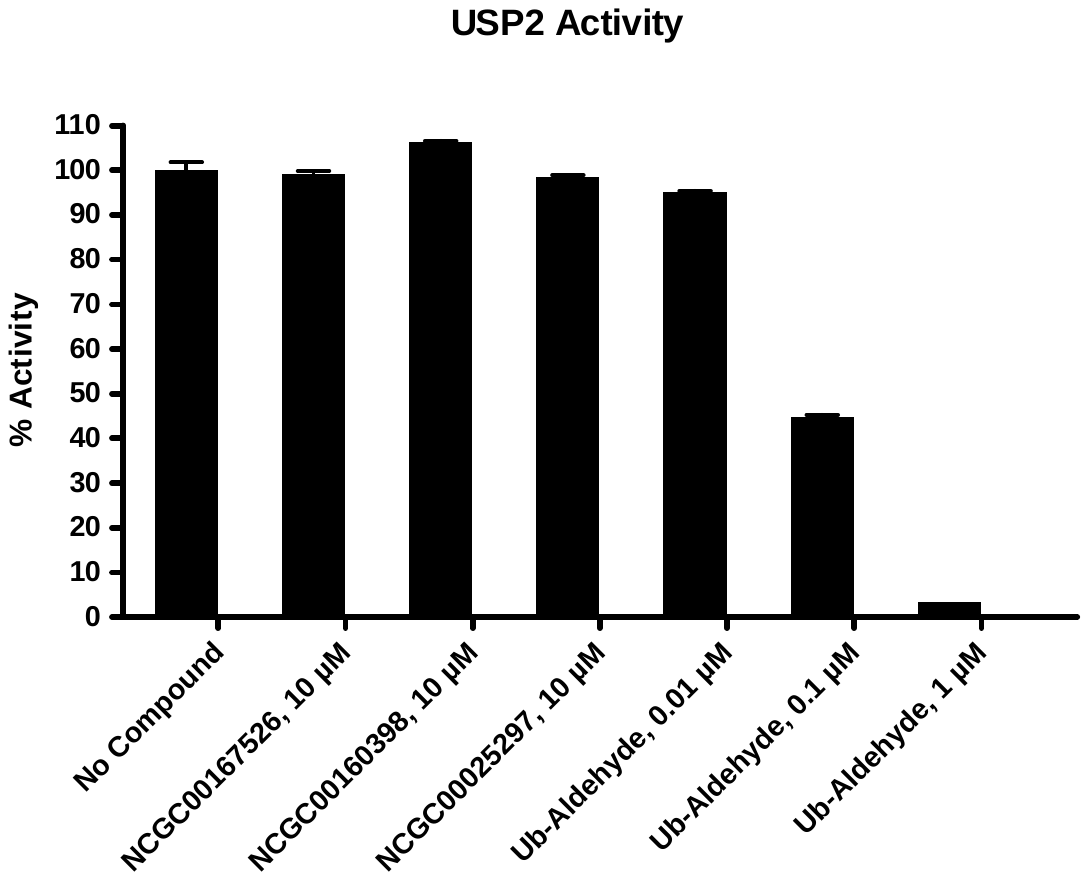

3.2.21. USP5

| Compounds | USP5 Activity (Fluorescence count) | | Background  (Fluorescence count) | | % Activity | | % Inhibition |
| --- | --- | --- | --- | --- | --- | --- | --- |
|  | Repeat1 | Repeat2 | Repeat1 | Repeat2 | Repeat1 | Repeat2 |  |
| No Compound | 854 | 867 | 21 | 22 | 99 | 101 | 0 |
| NCGC00167526, 10 μM | 835 | 862 | 20 | 22 | 97 | 100 | 1 |
| NCGC00160398, 10 μM | 927 | 953 | 67 | 72 | 103 | 105 | 0 |
| NCGC00025297, 10 μM | 854 | 856 | 23 | 23 | 99 | 99 | 1 |
| Ub-Aldehyde, 0.01 μM | 847 | 845 | 26 | 21 | 98 | 98 | 2 |
| Ub-Aldehyde, 0.1 μM | 388 | 391 | 23 | 24 | 43 | 43 | 57 |
| Ub-Aldehyde, 1 μM | 58 | 58 | 28 | 29 | 3 | 3 | 97 |
| Background | 29 | 13 | 14 | 14 |  |  |  |

**Table 3.2.21. Data for the Effect of the Compounds on USP5 Activity**

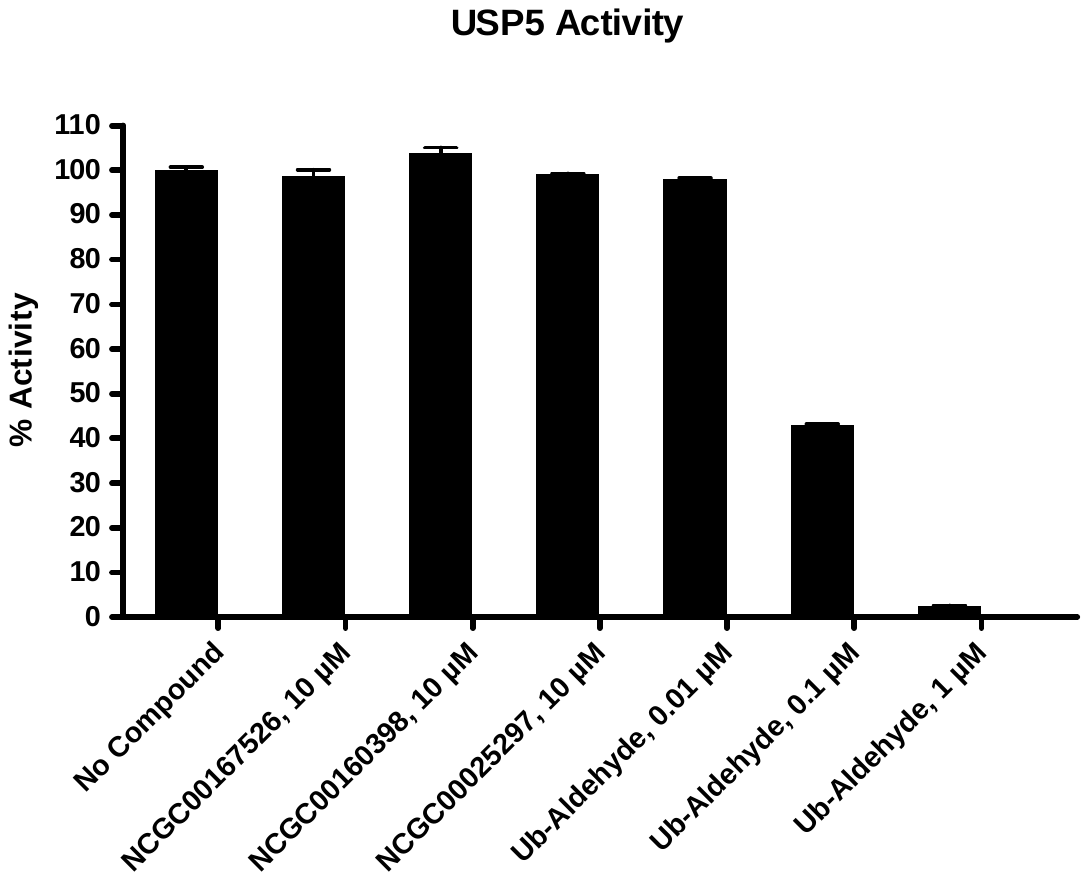

3.2.22. USP7

| Compounds | USP7 Activity (Fluorescence count) | | Background  (Fluorescence count) | | % Activity | | % Inhibition |
| --- | --- | --- | --- | --- | --- | --- | --- |
|  | Repeat1 | Repeat2 | Repeat1 | Repeat2 | Repeat1 | Repeat2 |  |
| No Compound | 524 | 523 | 28 | 29 | 100 | 100 | 0 |
| NCGC00167526, 10 μM | 535 | 512 | 30 | 27 | 102 | 98 | 0 |
| NCGC00160398, 10 μM | 583 | 593 | 73 | 79 | 103 | 104 | 0 |
| NCGC00025297, 10 μM | 529 | 522 | 29 | 29 | 101 | 100 | 0 |
| Ub-Aldehyde, 0.001 μM | 460 | 466 | 27 | 28 | 87 | 88 | 12 |
| Ub-Aldehyde, 0.01 μM | 250 | 243 | 28 | 26 | 45 | 44 | 56 |
| Ub-Aldehyde, 0.1 μM | 40 | 41 | 23 | 24 | 3 | 3 | 97 |
| Background | 13 | 11 | 12 | 10 |  |  |  |

**Table 3.2.22. Data for the Effect of the Compounds on USP7 Activity**

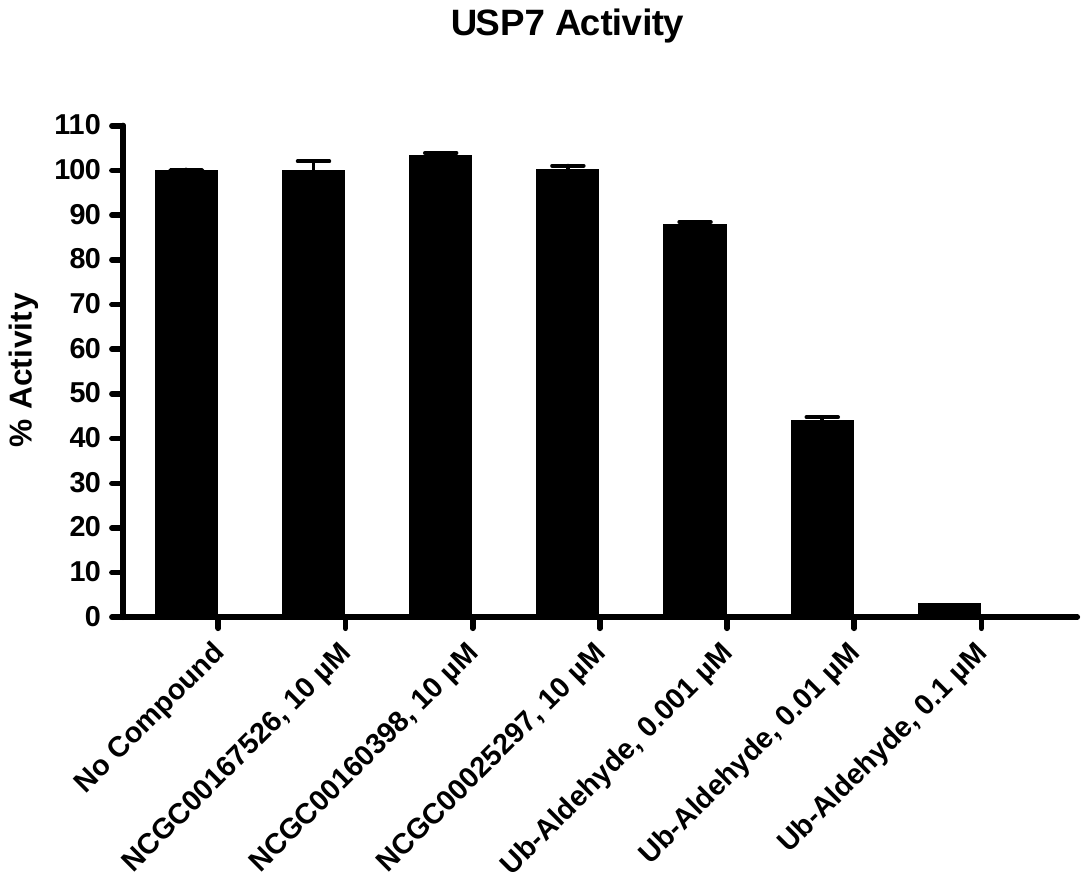

3.2.23. USP8

| Compounds | USP8 Activity (Fluorescence count) | | Background  (Fluorescence count) | | % Activity | | % Inhibition |
| --- | --- | --- | --- | --- | --- | --- | --- |
|  | Repeat1 | Repeat2 | Repeat1 | Repeat2 | Repeat1 | Repeat2 |  |
| No Compound | 210 | 213 | 19 | 20 | 99 | 101 | 0 |
| NCGC00167526, 10 μM | 220 | 213 | 20 | 19 | 104 | 101 | 0 |
| NCGC00160398, 10 μM | 247 | 254 | 64 | 71 | 95 | 95 | 5 |
| NCGC00025297, 10 μM | 212 | 219 | 20 | 20 | 100 | 104 | 0 |
| Ub-Aldehyde, 0.01 μM | 192 | 188 | 19 | 19 | 90 | 88 | 11 |
| Ub-Aldehyde, 0.1 μM | 143 | 143 | 18 | 19 | 65 | 65 | 35 |
| Ub-Aldehyde, 1 μM | 47 | 47 | 18 | 17 | 15 | 16 | 85 |
| Background | 13 | 13 | 13 | 13 |  |  |  |

**Table 3.2.23. Data for the Effect of the Compounds on USP8 Activity**

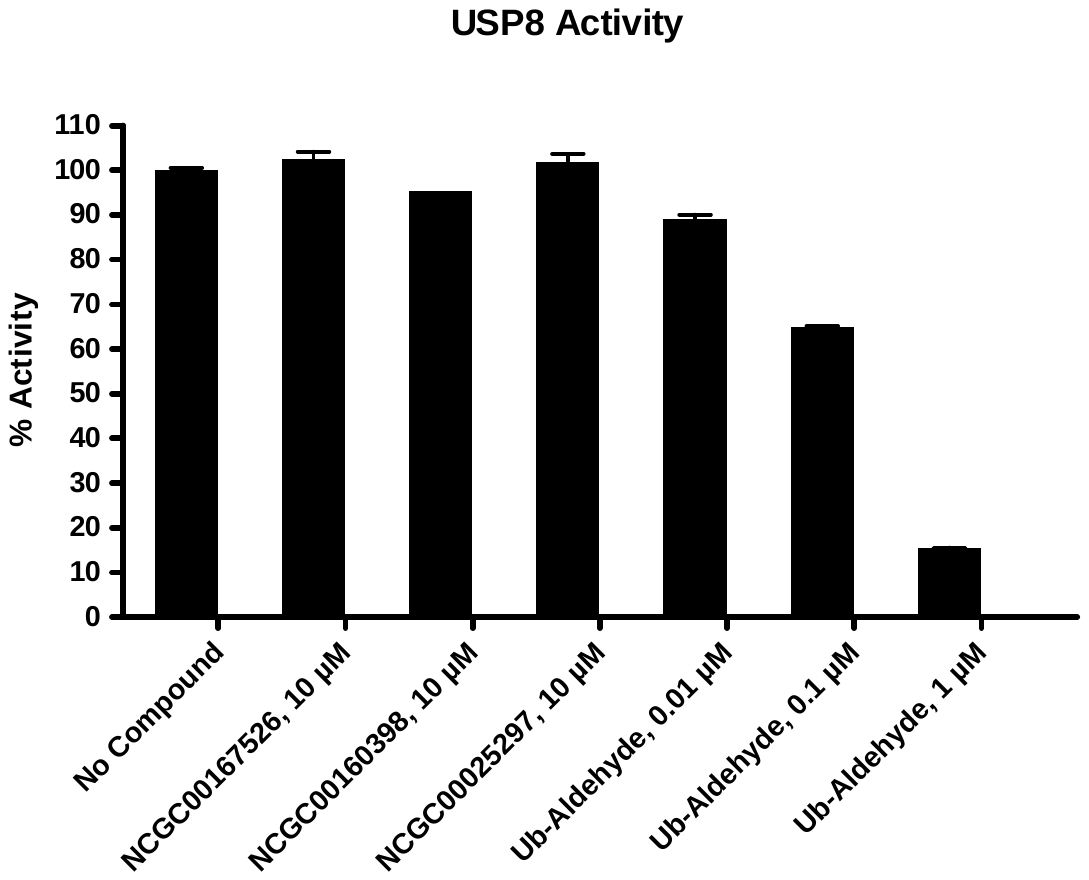

3.2.24. USP10

| Compounds | USP10 Activity (Fluorescence count) | | Background  (Fluorescence count) | | % Activity | | % Inhibition |
| --- | --- | --- | --- | --- | --- | --- | --- |
|  | Repeat1 | Repeat2 | Repeat1 | Repeat2 | Repeat1 | Repeat2 |  |
| No Compound | 331 | 353 | 28 | 33 | 97 | 103 | 0 |
| NCGC00167526, 10 μM | 321 | 349 | 28 | 31 | 94 | 102 | 2 |
| NCGC00160398, 10 μM | 395 | 399 | 77 | 86 | 102 | 100 | 0 |
| NCGC00025297, 10 μM | 325 | 347 | 31 | 34 | 94 | 100 | 3 |
| Ub-Aldehyde, 0.01 μM | 327 | 325 | 35 | 27 | 94 | 96 | 5 |
| Ub-Aldehyde, 0.1 μM | 207 | 217 | 28 | 34 | 57 | 58 | 43 |
| Ub-Aldehyde, 1 μM | 86 | 97 | 29 | 41 | 17 | 17 | 83 |
| Background | 9 | 9 | 14 | 14 |  |  |  |

**Table 3.2.24. Data for the Effect of the Compounds on USP10 Activity**

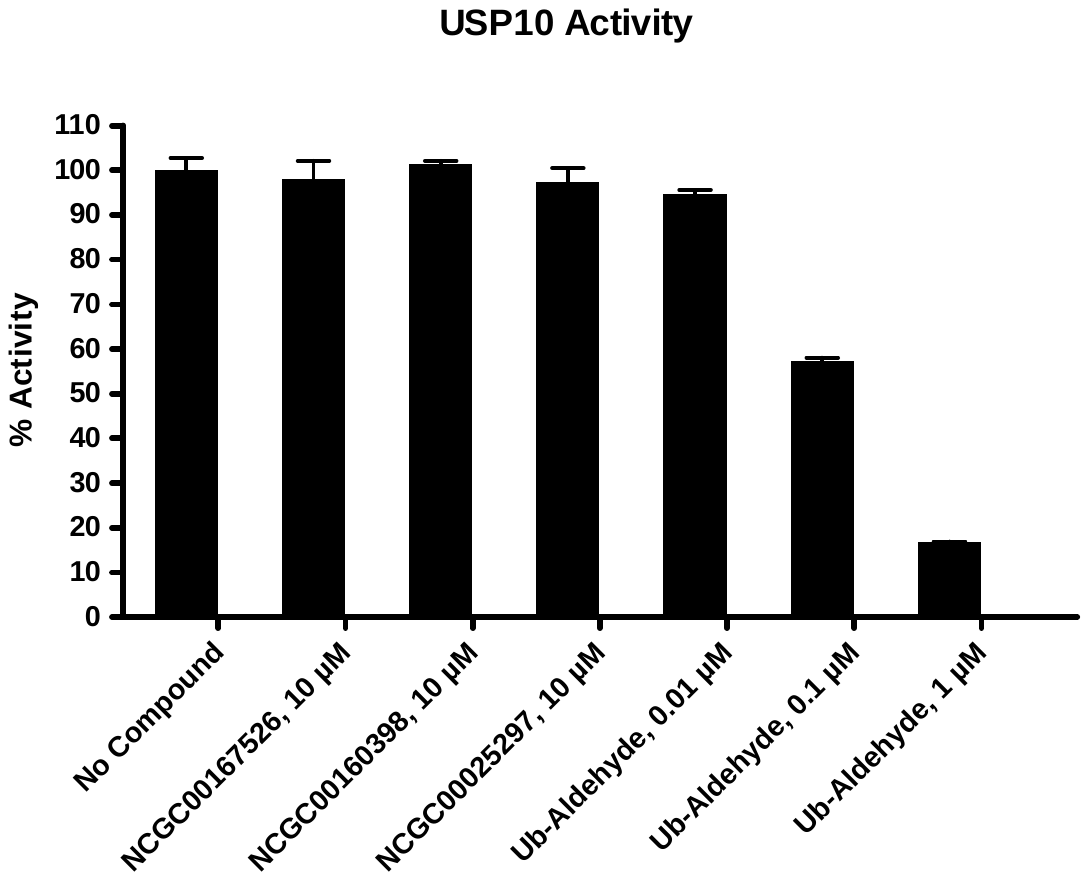

3.2.25. USP14

| Compounds | USP14 Activity (Fluorescence count) | | Background  (Fluorescence count) | | % Activity | | % Inhibition |
| --- | --- | --- | --- | --- | --- | --- | --- |
|  | Repeat1 | Repeat2 | Repeat1 | Repeat2 | Repeat1 | Repeat2 |  |
| No Compound | 57 | 66 | 17 | 18 | 90 | 110 | 0 |
| NCGC00167526, 10 μM | 58 | 66 | 16 | 17 | 95 | 113 | 0 |
| NCGC00160398, 10 μM | 139 | 136 | 62 | 57 | 184 | 189 | 0 |
| NCGC00025297, 10 μM | 58 | 65 | 18 | 18 | 90 | 108 | 1 |
| Ub-Aldehyde, 0.001 μM | 57 | 52 | 19 | 17 | 85 | 77 | 19 |
| Ub-Aldehyde, 0.01 μM | 34 | 32 | 18 | 17 | 29 | 27 | 72 |
| Ub-Aldehyde, 0.1 μM | 17 | 18 | 17 | 18 | 0 | 0 | 100 |
| Background | 11 | 11 | 16 | 15 |  |  |  |

**Table 3.2.25. Data for the Effect of the Compounds on USP14 Activity**

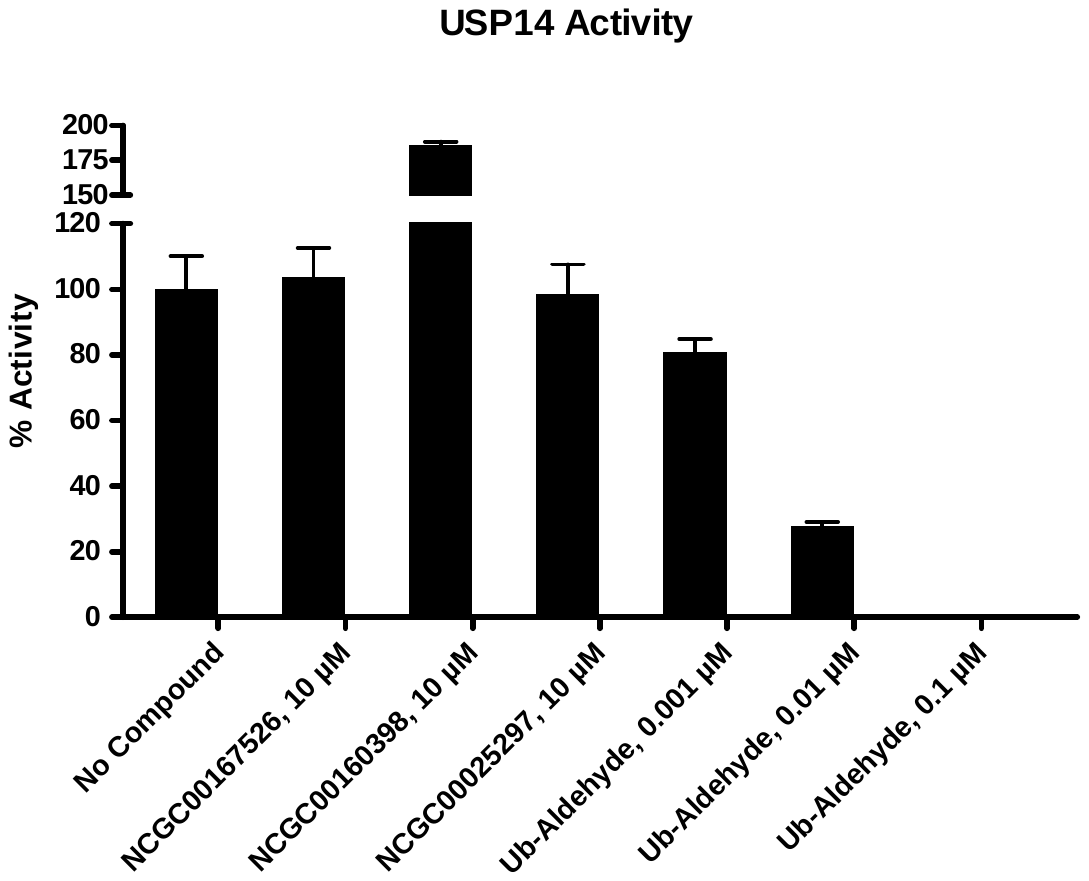

3.2.26. ADAM17

| Compounds | ADAM17 Activity (Fluorescence count) | | Background  (Fluorescence count) | | % Activity | | % Inhibition |
| --- | --- | --- | --- | --- | --- | --- | --- |
|  | Repeat1 | Repeat2 | Repeat1 | Repeat2 | Repeat1 | Repeat2 |  |
| No Compound | 42801 | 42329 | 6838 | 6676 | 100 | 100 | 0 |
| NCGC00167526, 10 μM | 40520 | 41117 | 7128 | 7511 | 93 | 94 | 6 |
| NCGC00160398, 10 μM | 40625 | 41349 | 6852 | 7156 | 94 | 95 | 5 |
| NCGC00025297, 10 μM | 40489 | 39926 | 6784 | 6885 | 94 | 92 | 7 |
| TAPI-2, 0.1 μM | 42733 | 42649 | 7037 | 6796 | 100 | 100 | 0 |
| TAPI-2, 1 μM | 39241 | 36415 | 6823 | 6600 | 90 | 83 | 13 |
| TAPI-2, 10 μM | 17326 | 15501 | 6635 | 6396 | 30 | 25 | 73 |
| Background | 5504 | 5482 | 5308 | 5350 |  |  |  |

**Table 3.2.26. Data for the Effect of the Compounds on ADAM17 Activity**

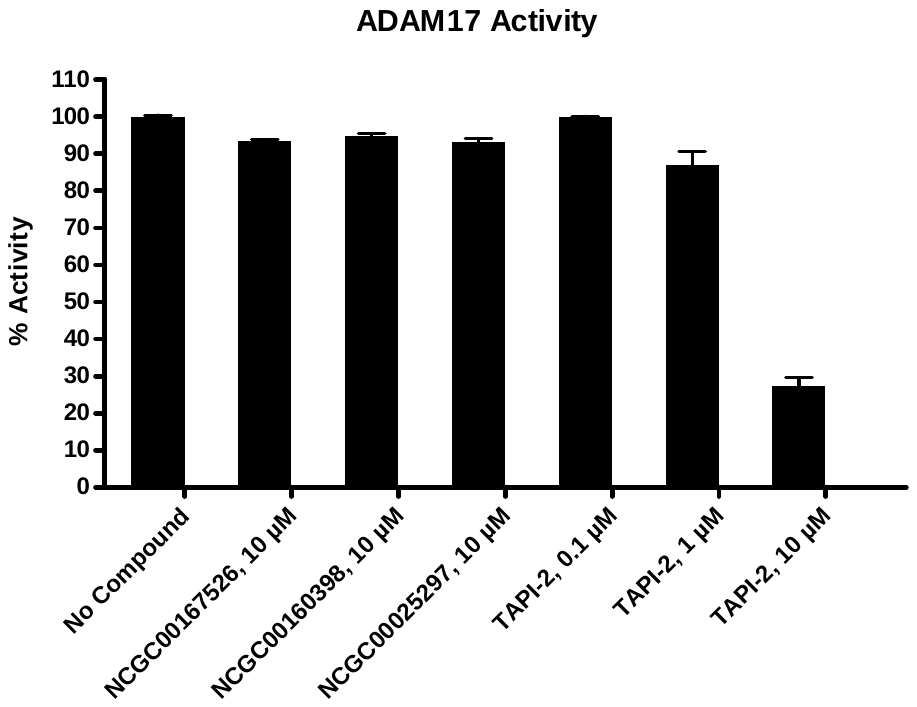

3.2.27. MMP1

| Compounds | MMP1 Activity (Fluorescence count) | | Background  (Fluorescence count) | | % Activity | | % Inhibition |
| --- | --- | --- | --- | --- | --- | --- | --- |
|  | Repeat1 | Repeat2 | Repeat1 | Repeat2 | Repeat1 | Repeat2 |  |
| No Compound | 2709 | 2726 | 587 | 591 | 100 | 100 | 0 |
| NCGC00167526, 10 μM | 2638 | 2631 | 569 | 564 | 97 | 97 | 3 |
| NCGC00160398, 10 μM | 3274 | 3220 | 1040 | 1015 | 105 | 104 | 0 |
| NCGC00025297, 10 μM | 2649 | 2660 | 560 | 562 | 98 | 99 | 2 |
| NNGH, 0.1 μM | 2639 | 2645 | 548 | 548 | 98 | 98 | 2 |
| NNGH, 1 μM | 2138 | 2130 | 558 | 550 | 74 | 74 | 26 |
| NNGH, 10 μM | 1066 | 1039 | 537 | 511 | 23 | 23 | 77 |
| Background | 533 | 536 | 486 | 491 |  |  |  |

**Table 3.2.27. Data for the Effect of the Compounds on MMP1 Activity**

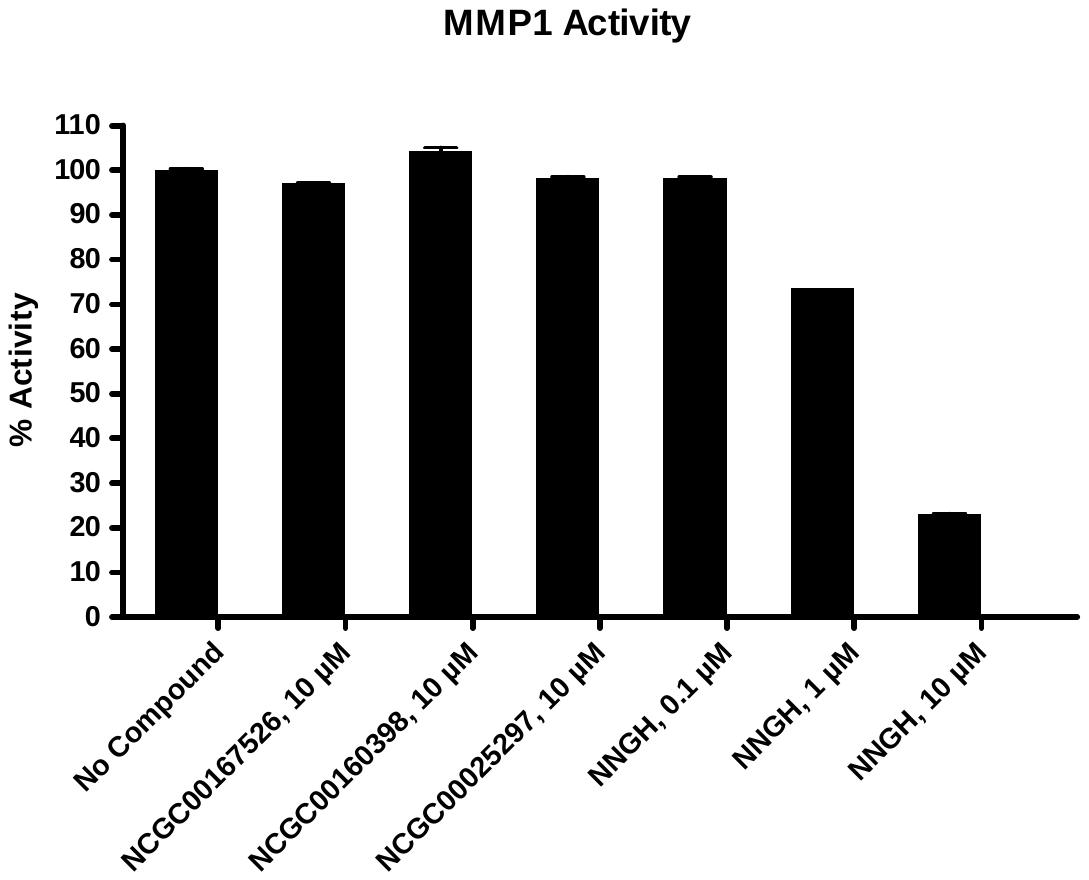

3.2.28. MMP2

| Compounds | MMP2 Activity (Fluorescence count) | | Background  (Fluorescence count) | | % Activity | | % Inhibition |
| --- | --- | --- | --- | --- | --- | --- | --- |
|  | Repeat1 | Repeat2 | Repeat1 | Repeat2 | Repeat1 | Repeat2 |  |
| No Compound | 337 | 335 | 158 | 158 | 101 | 99 | 0 |
| NCGC00167526, 10 μM | 210 | 198 | 160 | 161 | 21 | 13 | 83 |
| NCGC00160398, 10 μM | 985 | 997 | 627 | 618 | 211 | 224 | 0 |
| NCGC00025297, 10 μM | 303 | 286 | 166 | 153 | 75 | 72 | 26 |
| NNGH, 0.01 μM | 290 | 285 | 154 | 143 | 74 | 78 | 24 |
| NNGH, 0.1 μM | 207 | 205 | 143 | 144 | 30 | 28 | 71 |
| NNGH, 1 μM | 160 | 160 | 151 | 146 | 0 | 0 | 100 |
| Background | 163 | 163 | 147 | 148 |  |  |  |

**Table 3.2.28. Data for the Effect of the Compounds on MMP2 Activity**

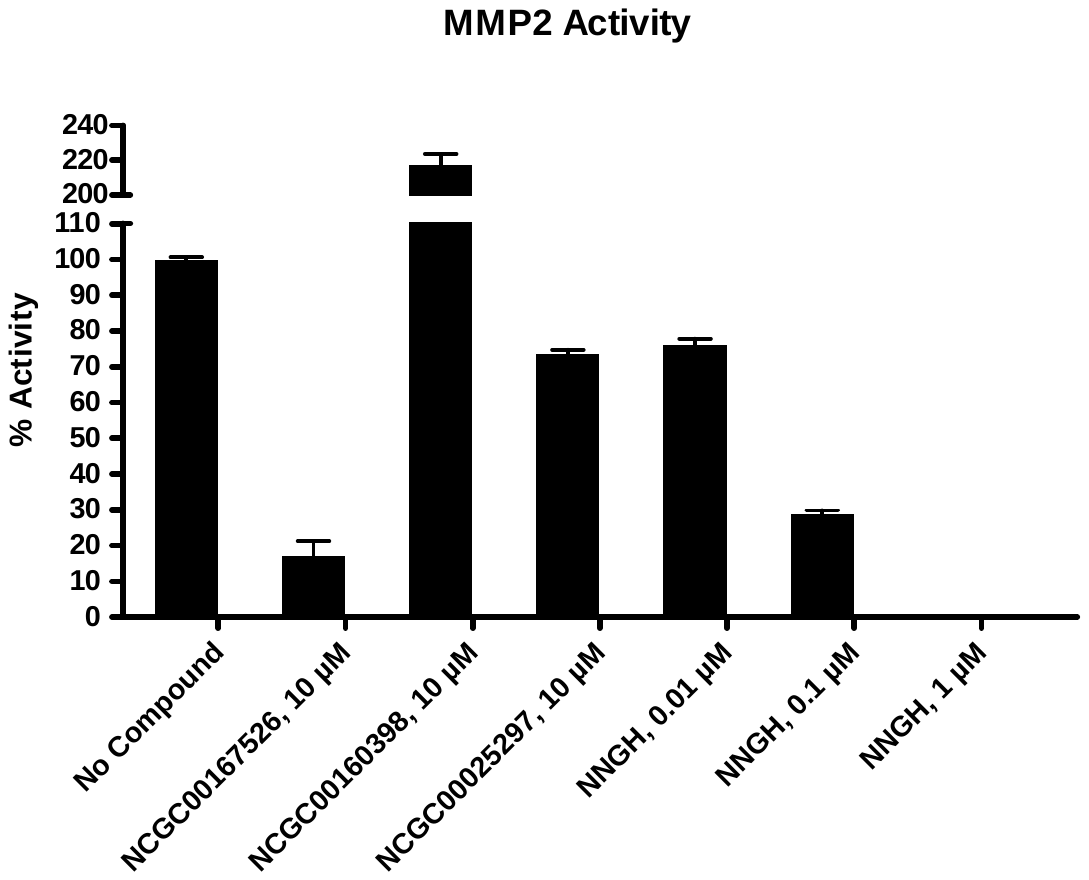

3.2.29. MMP3

| Compounds | MMP3 Activity (Fluorescence count) | | Background  (Fluorescence count) | | % Activity | | % Inhibition |
| --- | --- | --- | --- | --- | --- | --- | --- |
|  | Repeat1 | Repeat2 | Repeat1 | Repeat2 | Repeat1 | Repeat2 |  |
| No Compound | 470 | 480 | 258 | 268 | 100 | 100 | 0 |
| NCGC00167526, 10 μM | 421 | 422 | 266 | 265 | 73 | 74 | 27 |
| NCGC00160398, 10 μM | 1054 | 1028 | 688 | 684 | 173 | 163 | 0 |
| NCGC00025297, 10 μM | 417 | 428 | 245 | 253 | 81 | 82 | 18 |
| NNGH, 0.01 μM | 426 | 422 | 256 | 253 | 80 | 80 | 20 |
| NNGH, 0.1 μM | 298 | 298 | 255 | 253 | 20 | 20 | 80 |
| NNGH, 1 μM | 242 | 241 | 242 | 245 | 0 | 1 | 100 |
| Background | 144 | 149 | 144 | 145 |  |  |  |

**Table 3.2.29. Data for the Effect of the Compounds on MMP3 Activity**

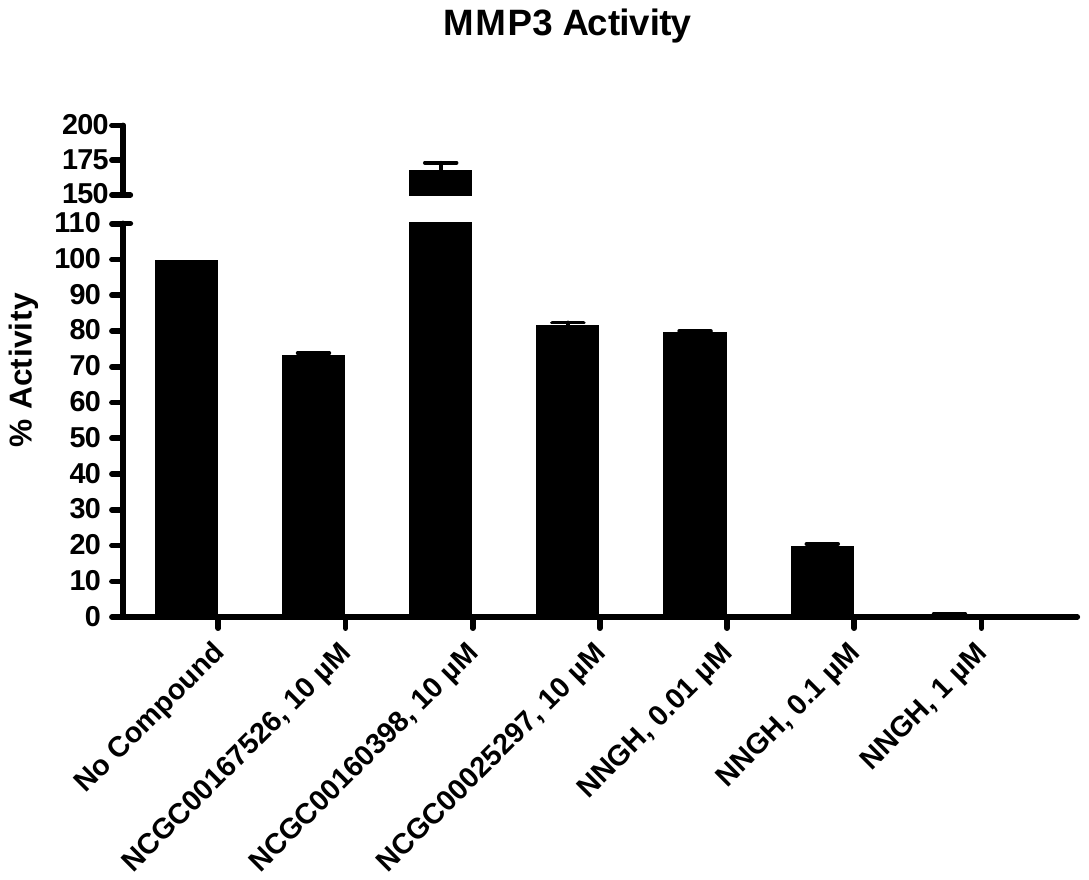

3.2.30. MMP7

| Compounds | MMP7 Activity (Fluorescence count) | | Background  (Fluorescence count) | | % Activity | | % Inhibition |
| --- | --- | --- | --- | --- | --- | --- | --- |
|  | Repeat1 | Repeat2 | Repeat1 | Repeat2 | Repeat1 | Repeat2 |  |
| No Compound | 13151 | 13336 | 721 | 821 | 100 | 100 | 0 |
| NCGC00167526, 10 μM | 13024 | 12955 | 731 | 811 | 99 | 97 | 2 |
| NCGC00160398, 10 μM | 13050 | 13055 | 1193 | 1214 | 95 | 95 | 5 |
| NCGC00025297, 10 μM | 12727 | 12789 | 661 | 651 | 97 | 97 | 3 |
| Batimastat, 0.0001 μM | 12628 | 12882 | 792 | 827 | 95 | 97 | 4 |
| Batimastat, 0.001 μM | 9330 | 9338 | 707 | 679 | 69 | 69 | 31 |
| Batimastat, 0.01 μM | 1369 | 1398 | 587 | 624 | 6 | 6 | 94 |
| Background | 161 | 159 | 138 | 135 |  |  |  |

**Table 3.2.30. Data for the Effect of the Compounds on MMP7 Activity**

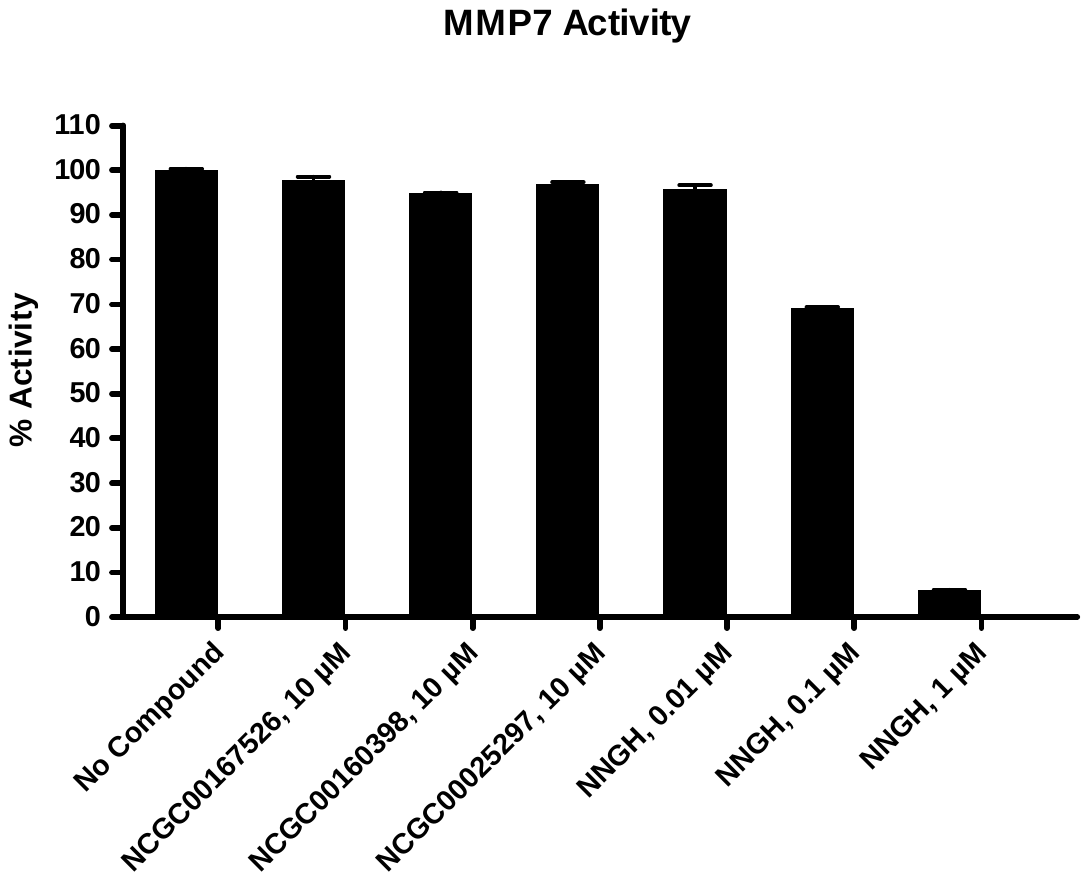

3.2.31. MMP8

| Compounds | MMP8 Activity (Fluorescence count) | | Background  (Fluorescence count) | | % Activity | | % Inhibition |
| --- | --- | --- | --- | --- | --- | --- | --- |
|  | Repeat1 | Repeat2 | Repeat1 | Repeat2 | Repeat1 | Repeat2 |  |
| No Compound | 268 | 264 | 92 | 86 | 99 | 101 | 0 |
| NCGC00167526, 10 μM | 216 | 214 | 99 | 91 | 66 | 69 | 33 |
| NCGC00160398, 10 μM | 951 | 952 | 510 | 514 | 251 | 249 | 0 |
| NCGC00025297, 10 μM | 247 | 254 | 83 | 89 | 93 | 93 | 7 |
| NNGH, 0.01 μM | 256 | 251 | 90 | 81 | 94 | 96 | 5 |
| NNGH, 0.1 μM | 189 | 187 | 88 | 83 | 57 | 58 | 43 |
| NNGH, 1 μM | 101 | 105 | 82 | 87 | 10 | 9 | 91 |
| Background | 85 | 80 | 87 | 82 |  |  |  |

**Table 3.2.31. Data for the Effect of the Compounds on MMP8 Activity**

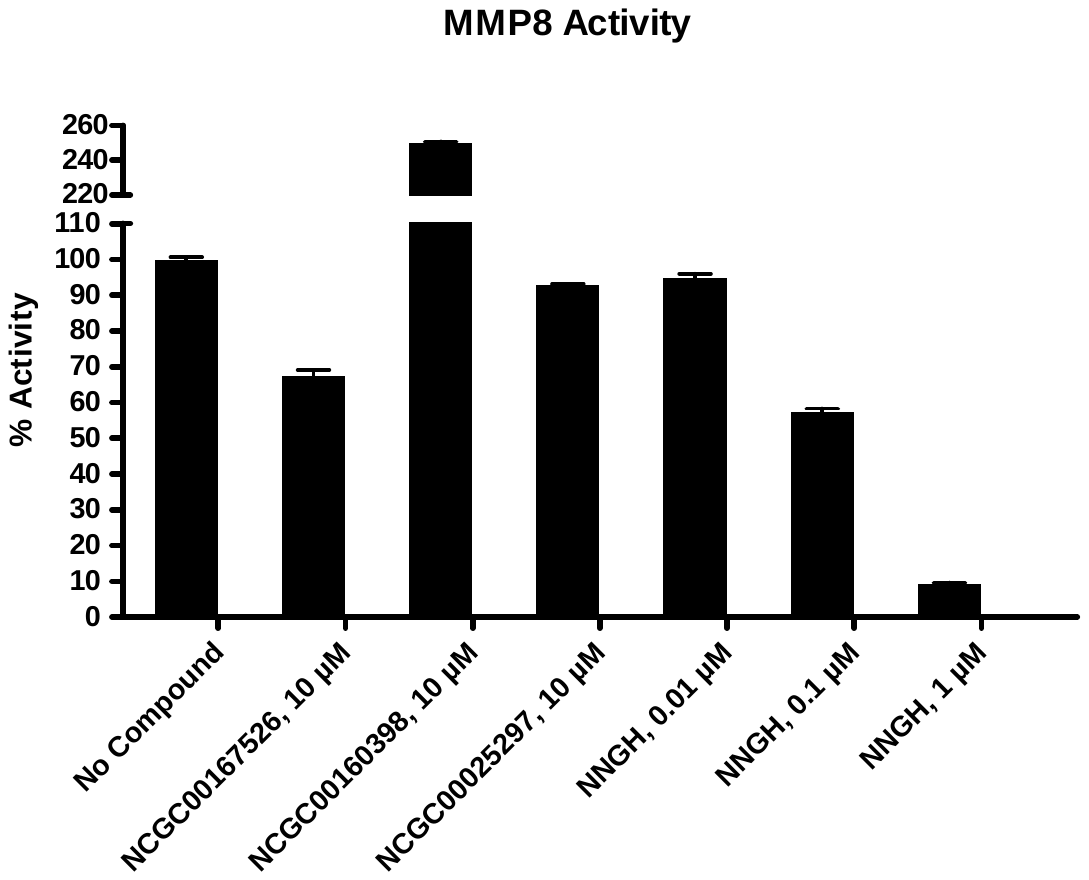

3.2.32. MMP9 (Q279R)

| Compounds | MMP9 (Q279R) Activity (Fluorescence count) | | Background  (Fluorescence count) | | % Activity | | % Inhibition |
| --- | --- | --- | --- | --- | --- | --- | --- |
|  | Repeat1 | Repeat2 | Repeat1 | Repeat2 | Repeat1 | Repeat2 |  |
| No Compound | 731 | 718 | 93 | 85 | 100 | 100 | 0 |
| NCGC00167526, 10 μM | 682 | 673 | 93 | 87 | 93 | 92 | 8 |
| NCGC00160398, 10 μM | 854 | 868 | 286 | 306 | 89 | 88 | 11 |
| NCGC00025297, 10 μM | 639 | 648 | 84 | 93 | 87 | 87 | 13 |
| NNGH, 0.001 μM | 558 | 540 | 85 | 83 | 74 | 72 | 27 |
| NNGH, 0.01 μM | 245 | 258 | 82 | 85 | 26 | 27 | 74 |
| NNGH, 0.1 μM | 110 | 105 | 81 | 79 | 4 | 4 | 96 |
| Background | 84 | 85 | 84 | 84 |  |  |  |

**Table 3.2.32. Data for the Effect of the Compounds on MMP9 (Q279R) Activity**

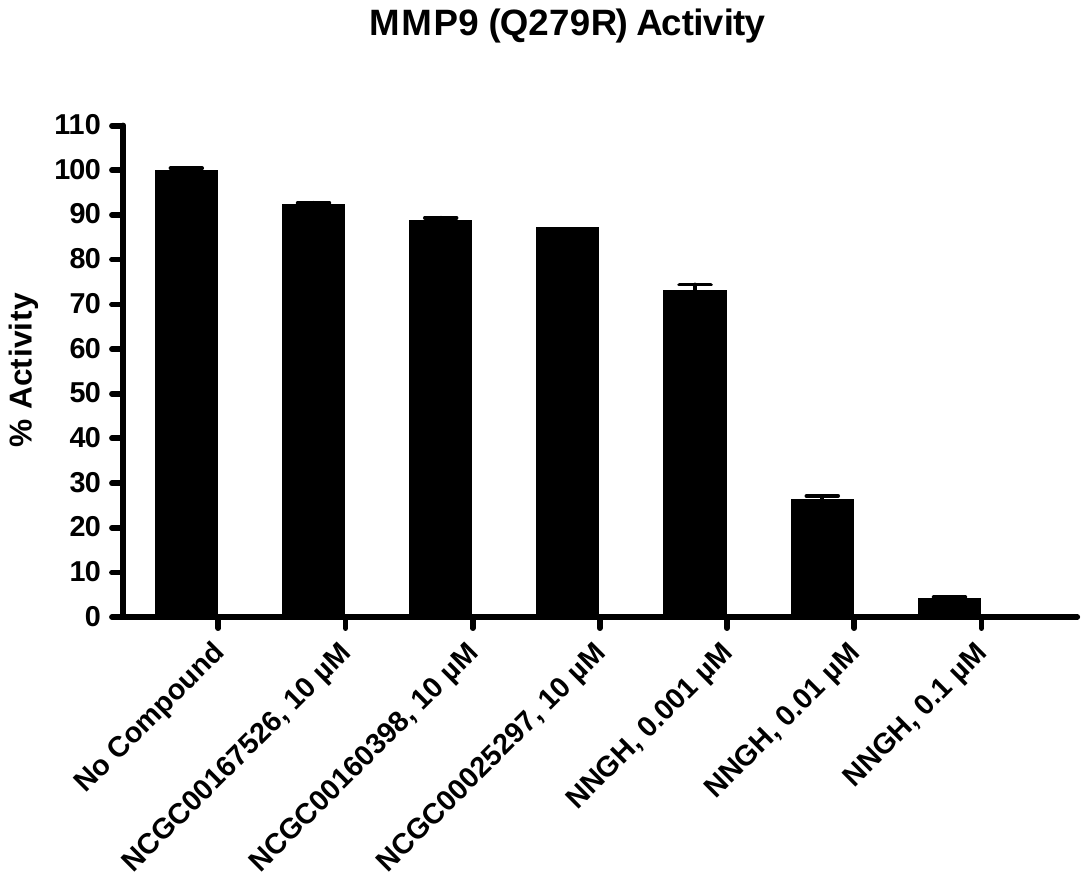

3.2.33. MMP10

| Compounds | MMP10 Activity (Fluorescence count) | | Background  (Fluorescence count) | | % Activity | | % Inhibition |
| --- | --- | --- | --- | --- | --- | --- | --- |
|  | Repeat1 | Repeat2 | Repeat1 | Repeat2 | Repeat1 | Repeat2 |  |
| No Compound | 312 | 318 | 141 | 141 | 98 | 102 | 0 |
| NCGC00167526, 10 μM | 266 | 278 | 138 | 156 | 73 | 70 | 29 |
| NCGC00160398, 10 μM | 1038 | 1049 | 517 | 551 | 304 | 290 | 0 |
| NCGC00025297, 10 μM | 295 | 298 | 140 | 147 | 89 | 87 | 12 |
| NNGH, 0.1 μM | 275 | 277 | 141 | 149 | 77 | 73 | 25 |
| NNGH, 1 μM | 213 | 212 | 142 | 150 | 40 | 34 | 63 |
| NNGH, 10 μM | 152 | 154 | 132 | 135 | 10 | 9 | 91 |
| Background | 152 | 155 | 148 | 152 |  |  |  |

**Table 3.2.33. Data for the Effect of the Compounds on MMP10 Activity**

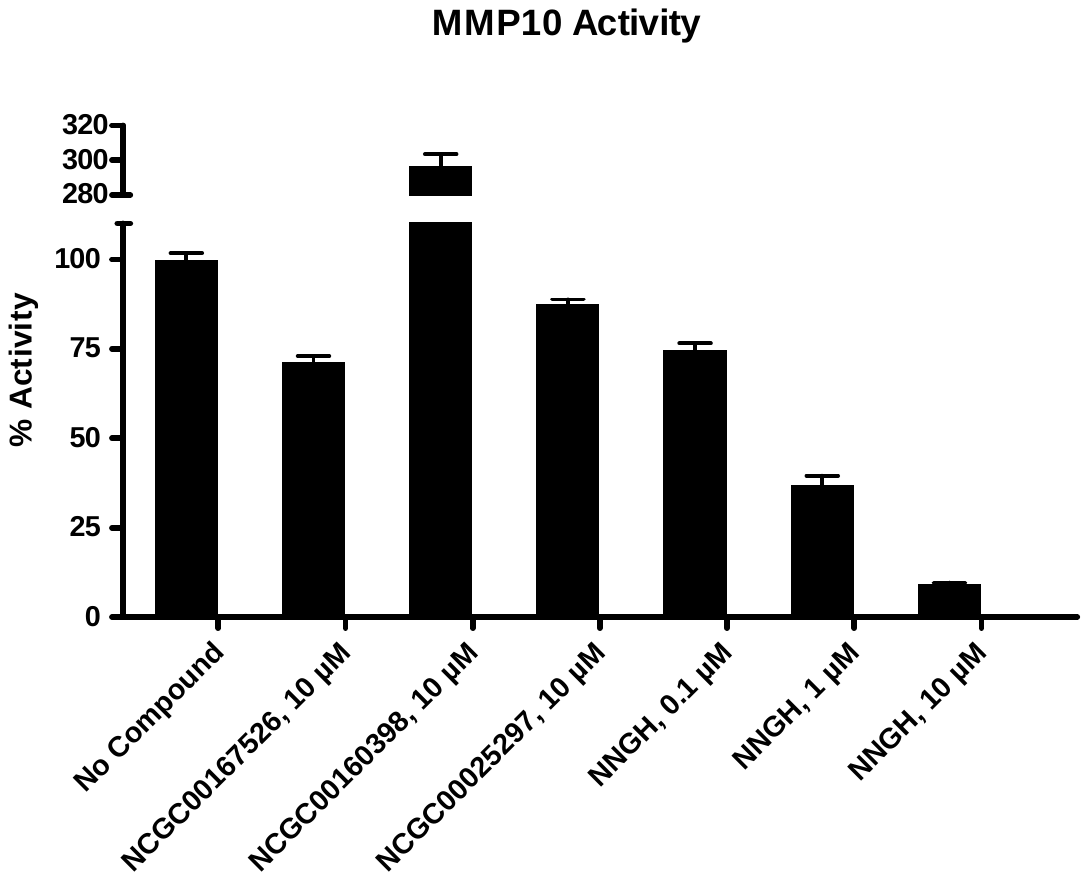

3.2.34. MMP13

| Compounds | MMP13 Activity (Fluorescence count) | | Background  (Fluorescence count) | | % Activity | | % Inhibition |
| --- | --- | --- | --- | --- | --- | --- | --- |
|  | Repeat1 | Repeat2 | Repeat1 | Repeat2 | Repeat1 | Repeat2 |  |
| No Compound | 9961 | 10322 | 705 | 985 | 100 | 100 | 0 |
| NCGC00167526, 10 μM | 7717 | 7857 | 653 | 773 | 76 | 76 | 24 |
| NCGC00160398, 10 μM | 3327 | 3360 | 999 | 1095 | 25 | 24 | 75 |
| NCGC00025297, 10 μM | 8927 | 8870 | 786 | 792 | 88 | 87 | 13 |
| Batimastat, 0.01 μM | 9094 | 9008 | 846 | 793 | 89 | 88 | 11 |
| Batimastat, 0.1 μM | 6273 | 6230 | 823 | 809 | 59 | 58 | 42 |
| Batimastat, 1 μM | 1221 | 1221 | 705 | 713 | 5 | 5 | 95 |
| Background | 155 | 157 | 150 | 149 |  |  |  |

**Table 3.2.34. Data for the Effect of the Compounds on MMP13 Activity**

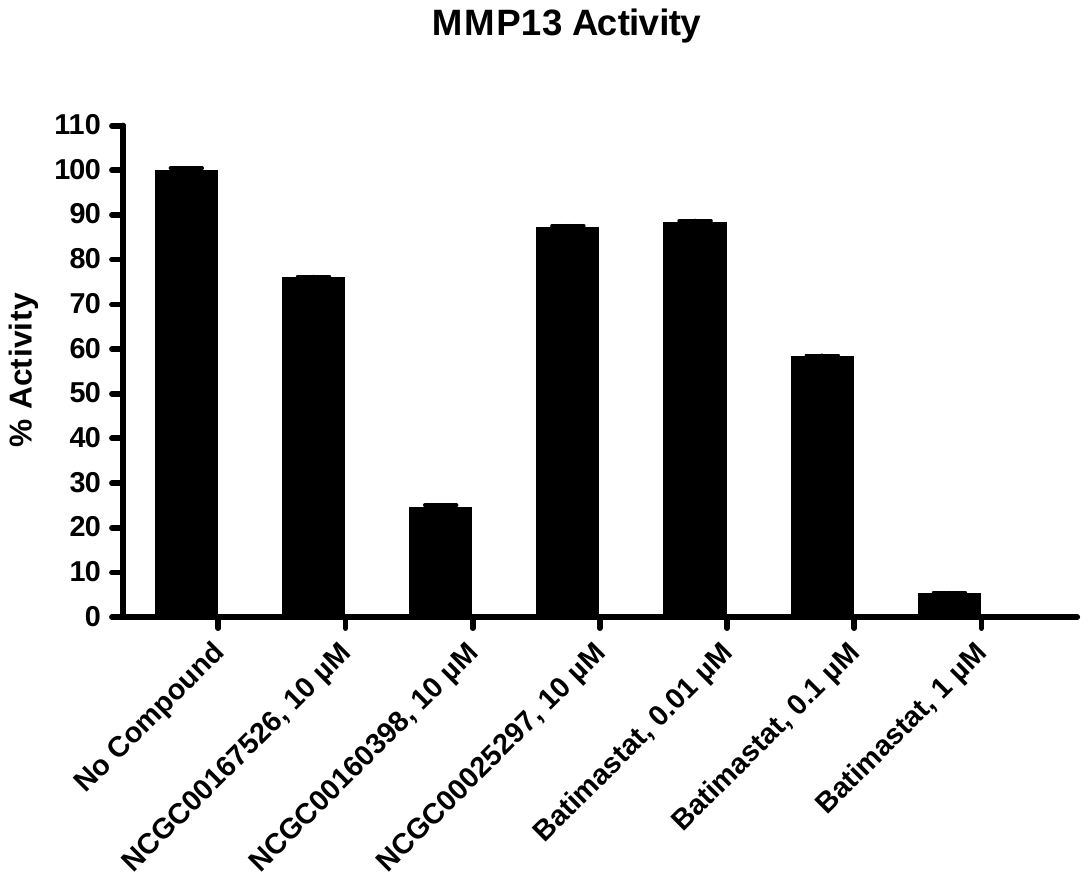

3.2.35. DPP3

| Compounds | DPP3 Activity (Fluorescence count) | | Background  (Fluorescence count) | | % Activity | | % Inhibition |
| --- | --- | --- | --- | --- | --- | --- | --- |
|  | Repeat1 | Repeat2 | Repeat1 | Repeat2 | Repeat1 | Repeat2 |  |
| No Compound | 44056 | 46143 | 3900 | 3999 | 98 | 102 | 0 |
| NCGC00167526, 10 μM | 42024 | 41828 | 4409 | 3715 | 91 | 93 | 8 |
| NCGC00160398, 10 μM | 43746 | 44531 | 3950 | 4271 | 97 | 98 | 3 |
| NCGC00025297, 10 μM | 44179 | 43330 | 4097 | 4267 | 97 | 95 | 4 |
| Spinorphine, 0.1 μM | 45714 | 50360 | 3738 | 3658 | 102 | 114 | 0 |
| Spinorphine, 1 μM | 9183 | 8460 | 1211 | 1335 | 19 | 17 | 82 |
| Spinorphine, 10 μM | 1716 | 1819 | 1139 | 1116 | 1 | 1 | 99 |
| Background | 1313 | 1356 | 1058 | 1078 |  |  |  |

**Table 3.2.35. Data for the Effect of the Compounds on DPP3 Activity**

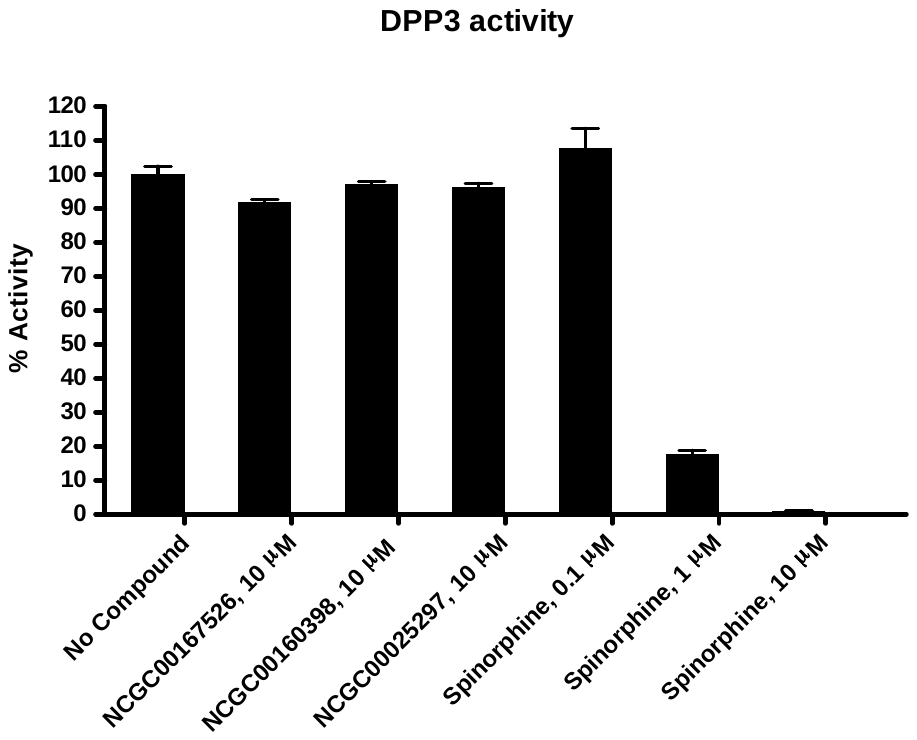

3.2.36. DPP4

| Compounds | DPP4 Activity (Fluorescence count) | | Background  (Fluorescence count) | | % Activity | | % Inhibition |
| --- | --- | --- | --- | --- | --- | --- | --- |
|  | Repeat1 | Repeat2 | Repeat1 | Repeat2 | Repeat1 | Repeat2 |  |
| No Compound | 18664 | 18230 | 3486 | 3255 | 101 | 99 | 0 |
| NCGC00167526, 10 μM | 14545 | 15960 | 2756 | 2799 | 78 | 87 | 17 |
| NCGC00160398, 10 μM | 15176 | 15667 | 2921 | 3110 | 81 | 83 | 18 |
| NCGC00025297, 10 μM | 14687 | 14879 | 2760 | 2848 | 79 | 80 | 21 |
| Sitagliptin, 0.001 μM | 14862 | 15705 | 2196 | 2265 | 84 | 89 | 13 |
| Sitagliptin, 0.01 μM | 8929 | 8882 | 1830 | 1723 | 47 | 47 | 53 |
| Sitagliptin, 0.1 μM | 2437 | 2454 | 1318 | 1365 | 7 | 7 | 93 |
| Background | 1276 | 1839 | 1195 | 1755 |  |  |  |

**Table 3.2.36. Data for the Effect of the Compounds on DPP4 Activity**

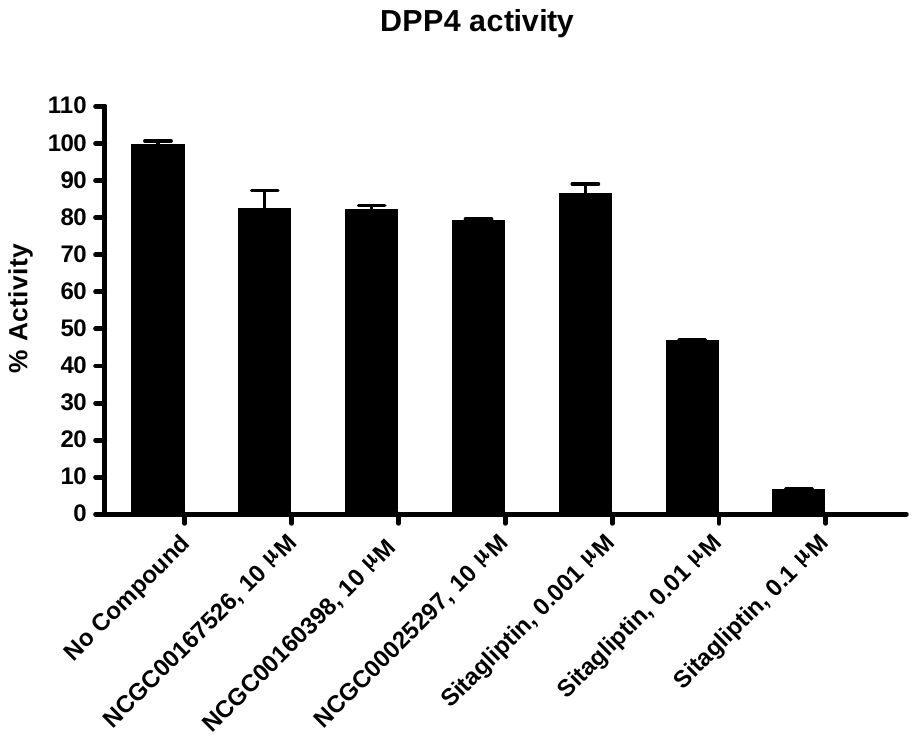

3.2.37. DPP7

| Compounds | DPP7 Activity (Fluorescence count) | | Background  (Fluorescence count) | | % Activity | | % Inhibition |
| --- | --- | --- | --- | --- | --- | --- | --- |
|  | Repeat1 | Repeat2 | Repeat1 | Repeat2 | Repeat1 | Repeat2 |  |
| No Compound | 4790 | 4678 | 566 | 485 | 100 | 100 | 0 |
| NCGC00167526, 10 μM | 3729 | 3792 | 380 | 388 | 80 | 81 | 20 |
| NCGC00160398, 10 μM | 3241 | 3507 | 375 | 403 | 68 | 74 | 29 |
| NCGC00025297, 10 μM | 660 | 1307 | 259 | 292 | 9 | 24 | 83 |
| KR62436, 1 μM | 4110 | 4599 | 472 | 469 | 86 | 98 | 8 |
| KR62436, 10 μM | 3018 | 3620 | 323 | 393 | 64 | 77 | 30 |
| KR62436, 100 μM | 1206 | 1304 | 229 | 295 | 23 | 24 | 77 |
| Background | 209 | 252 | 195 | 242 |  |  |  |

**Table 3.2.37. Data for the Effect of the Compounds on DPP7 Activity**

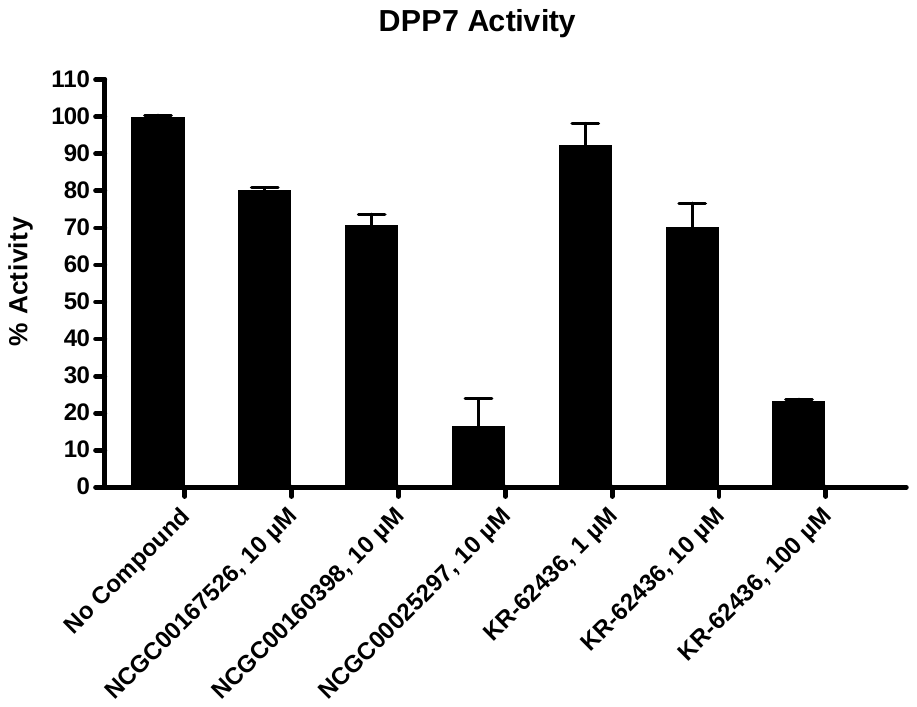

3.2.38. DPP8

| Compounds | DPP8 Activity (Fluorescence count) | | Background  (Fluorescence count) | | % Activity | | % Inhibition |
| --- | --- | --- | --- | --- | --- | --- | --- |
|  | Repeat1 | Repeat2 | Repeat1 | Repeat2 | Repeat1 | Repeat2 |  |
| No Compound | 12015 | 10890 | 1835 | 1616 | 105 | 95 | 0 |
| NCGC00167526, 10 μM | 10224 | 10376 | 1094 | 1602 | 94 | 90 | 8 |
| NCGC00160398, 10 μM | 10514 | 10167 | 1364 | 1390 | 94 | 90 | 8 |
| NCGC00025297, 10 μM | 9190 | 10396 | 1049 | 1665 | 84 | 90 | 13 |
| KR62436, 1 μM | 11007 | 11277 | 1348 | 1374 | 99 | 102 | 0 |
| KR62436, 10 μM | 10133 | 8763 | 1530 | 804 | 88 | 82 | 15 |
| KR62436, 100 μM | 2112 | 2332 | 310 | 336 | 18 | 20 | 81 |
| Background | 247 | 216 | 230 | 210 |  |  |  |

**Table 3.2.38. Data for the Effect of the Compounds on DPP8 Activity**

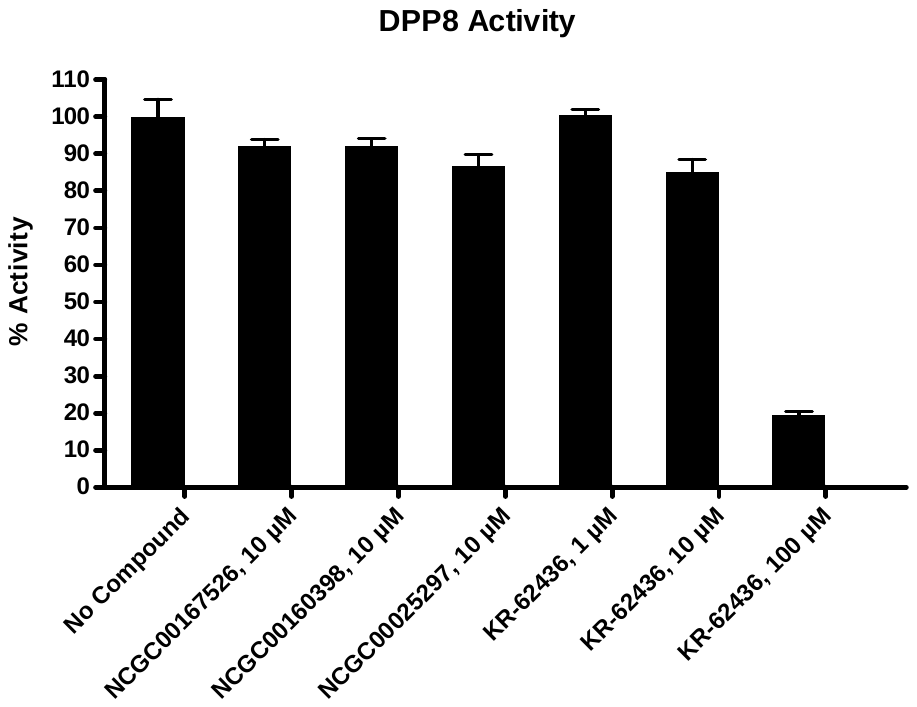

3.2.39. DPP9

| Compounds | DPP9 Activity (Fluorescence count) | | Background  (Fluorescence count) | | % Activity | | % Inhibition |
| --- | --- | --- | --- | --- | --- | --- | --- |
|  | Repeat1 | Repeat2 | Repeat1 | Repeat2 | Repeat1 | Repeat2 |  |
| No Compound | 6728 | 6726 | 1082 | 1102 | 100 | 100 | 0 |
| NCGC00167526, 10 μM | 5462 | 5396 | 829 | 819 | 82 | 81 | 18 |
| NCGC00160398, 10 μM | 5499 | 6006 | 808 | 923 | 83 | 90 | 13 |
| NCGC00025297, 10 μM | 5644 | 5849 | 838 | 914 | 85 | 88 | 14 |
| KR62436, 0.1 μM | 6633 | 7122 | 999 | 1125 | 100 | 106 | 0 |
| KR62436, 1 μM | 3343 | 3659 | 373 | 877 | 53 | 49 | 49 |
| KR62436, 10 μM | 1007 | 1018 | 269 | 249 | 13 | 14 | 87 |
| Background | 220 | 225 | 230 | 228 |  |  |  |

**Table 3.2.39. Data for the Effect of the Compounds on DPP9 Activity**

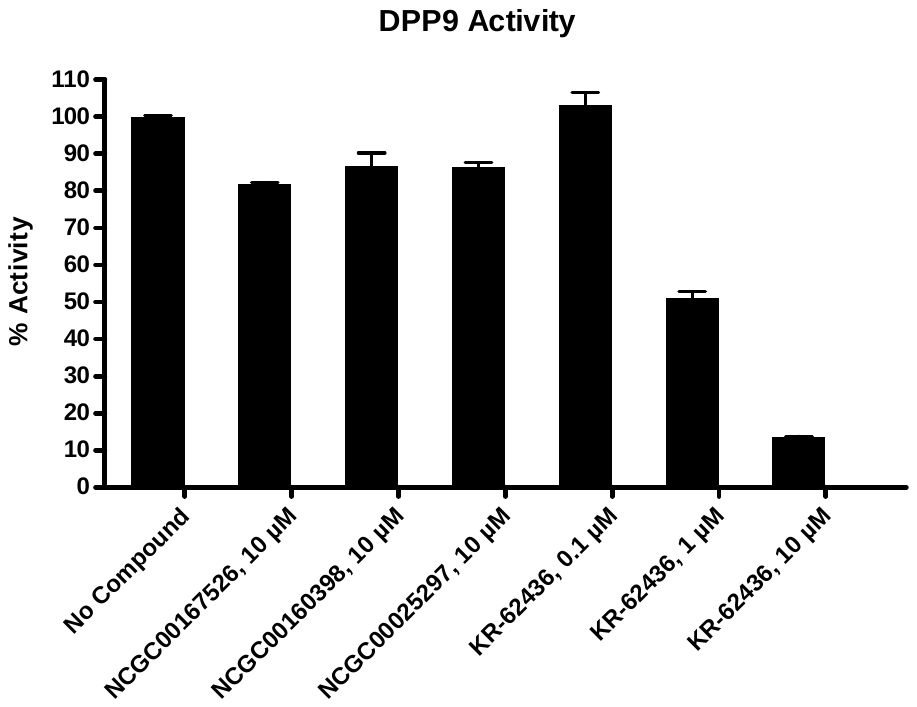

3.2.40. FAP

| Compounds | FAP Activity (Fluorescence count) | | Background  (Fluorescence count) | | % Activity | | % Inhibition |
| --- | --- | --- | --- | --- | --- | --- | --- |
|  | Repeat1 | Repeat2 | Repeat1 | Repeat2 | Repeat1 | Repeat2 |  |
| No Compound | 31645 | 31041 | 4753 | 4762 | 101 | 99 | 0 |
| NCGC00167526, 10 μM | 21101 | 21989 | 2816 | 3061 | 69 | 71 | 30 |
| NCGC00160398, 10 μM | 23833 | 23249 | 3786 | 3620 | 75 | 74 | 25 |
| NCGC00025297, 10 μM | 22699 | 22836 | 3896 | 3649 | 71 | 72 | 29 |
| SP-13786, 0.001 μM | 29765 | 30336 | 3599 | 3675 | 98 | 100 | 1 |
| SP-13786, 0.01 μM | 20566 | 20174 | 3256 | 2970 | 65 | 65 | 35 |
| SP-13786, 0.1 μM | 1365 | 1323 | 1258 | 1190 | 0 | 0 | 100 |
| Background | 1242 | 1400 | 1191 | 1369 |  |  |  |

**Table 3.2.40. Data for the Effect of the Compounds on FAP Activity**

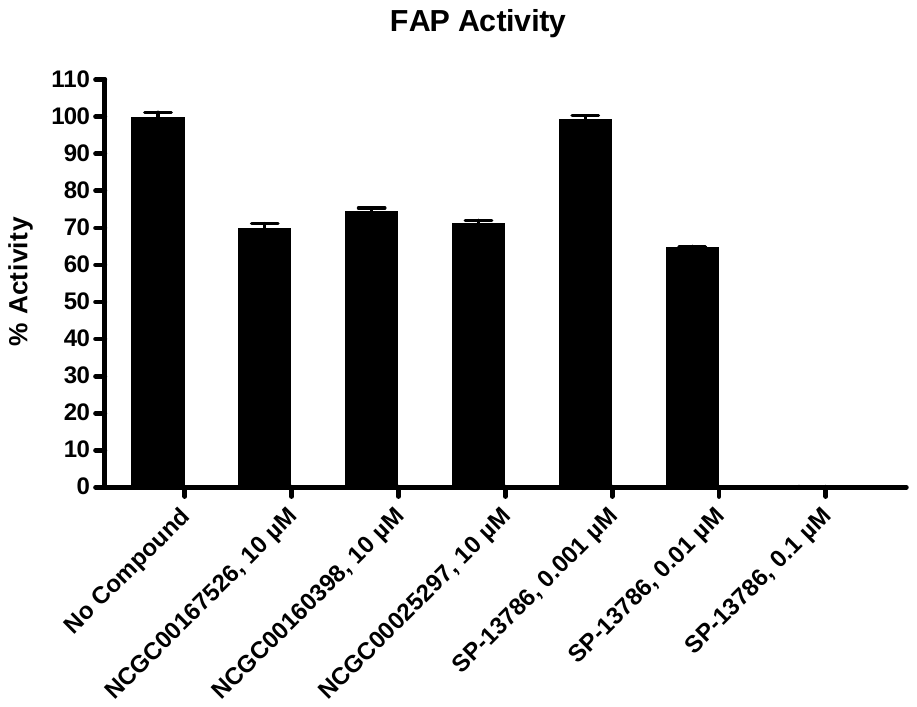

3.2.41. HCV1a (D168V)

| Compounds | HCV1a (D168V) Activity (Fluorescence count) | | Background  (Fluorescence count) | | % Activity | | % Inhibition |
| --- | --- | --- | --- | --- | --- | --- | --- |
|  | Repeat1 | Repeat2 | Repeat1 | Repeat2 | Repeat1 | Repeat2 |  |
| No Compound | 7562 | 7826 | 1245 | 1252 | 98 | 102 | 0 |
| NCGC00167526, 10 μM | 7711 | 7444 | 1432 | 1431 | 97 | 93 | 5 |
| NCGC00160398, 10 μM | 36331 | 36122 | 33942 | 34067 | 36 | 33 | 66 |
| NCGC00025297, 10 μM | 7889 | 7604 | 1353 | 1424 | 101 | 96 | 1 |
| Danoprevir, 0.001 μM | 6378 | 6687 | 1267 | 1274 | 79 | 84 | 18 |
| Danoprevir, 0.01 μM | 4164 | 3648 | 1306 | 1231 | 45 | 37 | 59 |
| Danoprevir, 0.1 μM | 1804 | 1819 | 1254 | 1275 | 8 | 8 | 92 |
| Background | 1245 | 1289 | 1243 | 1245 |  |  |  |

**Table 3.2.41. Data for the Effect of the Compounds on HCV1a (D168V) Activity**

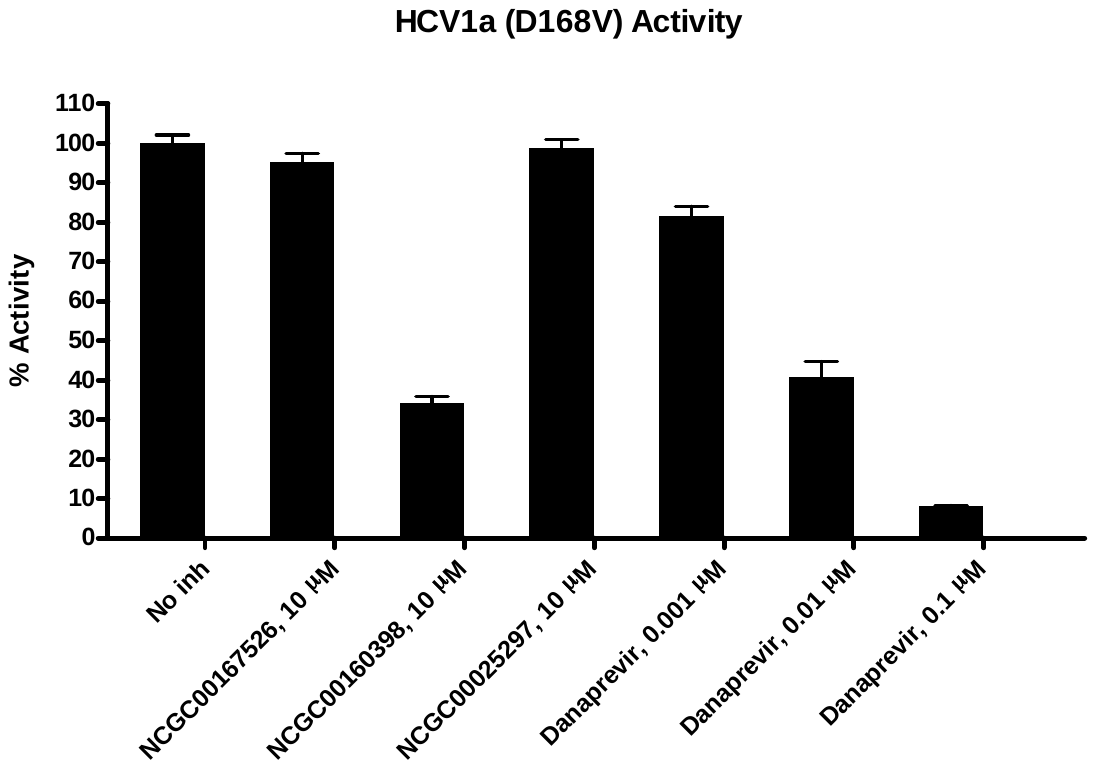

3.2.42. HCV1b

| Compounds | HCV1b Activity (Fluorescence count) | | Background  (Fluorescence count) | | % Activity | | % Inhibition |
| --- | --- | --- | --- | --- | --- | --- | --- |
|  | Repeat1 | Repeat2 | Repeat1 | Repeat2 | Repeat1 | Repeat2 |  |
| No Compound | 6588 | 6456 | 1245 | 1252 | 101 | 99 | 0 |
| NCGC00167526, 10 μM | 6301 | 5968 | 1432 | 1431 | 92 | 86 | 11 |
| NCGC00160398, 10 μM | 36598 | 37154 | 33942 | 34067 | 49 | 60 | 46 |
| NCGC00025297, 10 μM | 5076 | 5148 | 1353 | 1424 | 70 | 71 | 29 |
| Danoprevir, 0.0001 μM | 4972 | 5357 | 1254 | 1263 | 70 | 78 | 26 |
| Danoprevir, 0.001 μM | 3474 | 3493 | 1267 | 1274 | 42 | 42 | 58 |
| Danoprevir, 0.01 μM | 1681 | 1573 | 1306 | 1231 | 8 | 6 | 93 |
| Background | 1264 | 1252 | 1243 | 1245 |  |  |  |

**Table 3.2.42. Data for the Effect of the Compounds on HCV1b Activity**

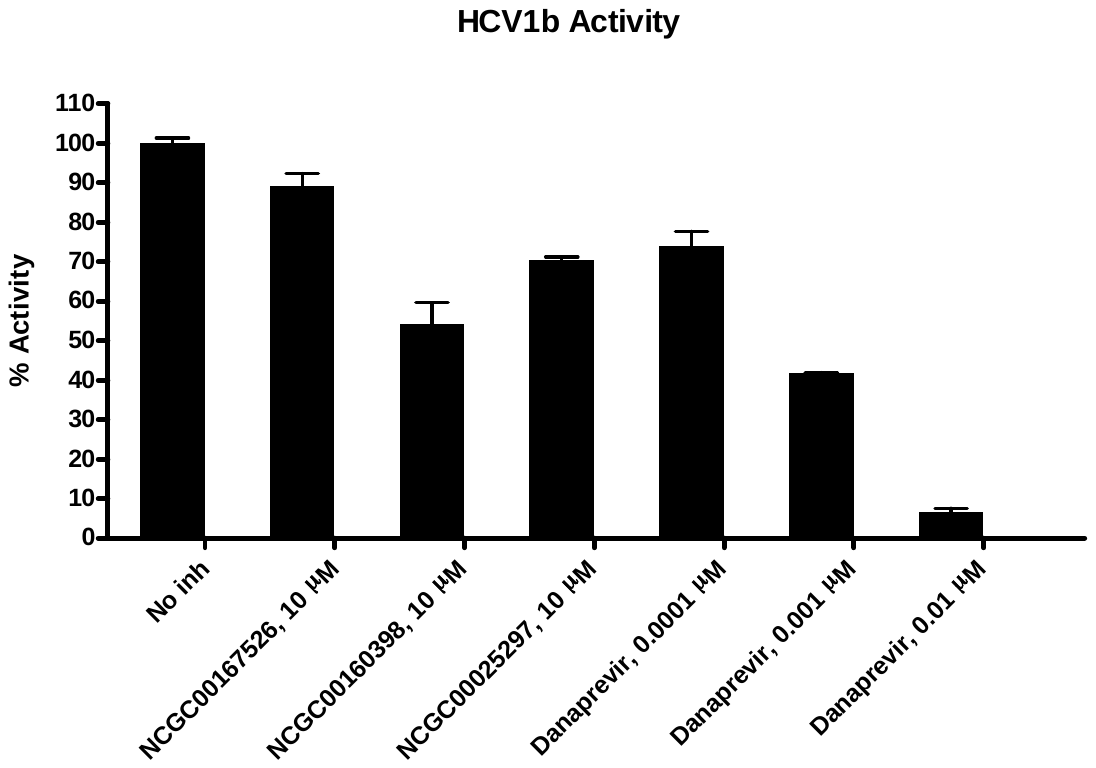

3.2.43. HCV1b (D168V)

| Compounds | HCV1b (D168V) Activity (Fluorescence count) | | Background  (Fluorescence count) | | % Activity | | % Inhibition |
| --- | --- | --- | --- | --- | --- | --- | --- |
|  | Repeat1 | Repeat2 | Repeat1 | Repeat2 | Repeat1 | Repeat2 |  |
| No Compound | 10955 | 10093 | 1245 | 1252 | 105 | 95 | 0 |
| NCGC00167526, 10 μM | 10444 | 10085 | 1432 | 1431 | 97 | 93 | 5 |
| NCGC00160398, 10 μM | 39196 | 39178 | 33942 | 34067 | 56 | 56 | 44 |
| NCGC00025297, 10 μM | 9567 | 9389 | 1353 | 1424 | 88 | 86 | 13 |
| Danoprevir, 0.001 μM | 7798 | 7735 | 1267 | 1274 | 70 | 70 | 30 |
| Danoprevir, 0.01 μM | 3387 | 3293 | 1306 | 1231 | 23 | 22 | 78 |
| Danoprevir, 0.1 μM | 1457 | 1467 | 1254 | 1275 | 2 | 2 | 98 |
| Background | 1257 | 1282 | 1243 | 1245 |  |  |  |

**Table 3.2.43. Data for the Effect of the Compounds on HCV1b (D168V) Activity**

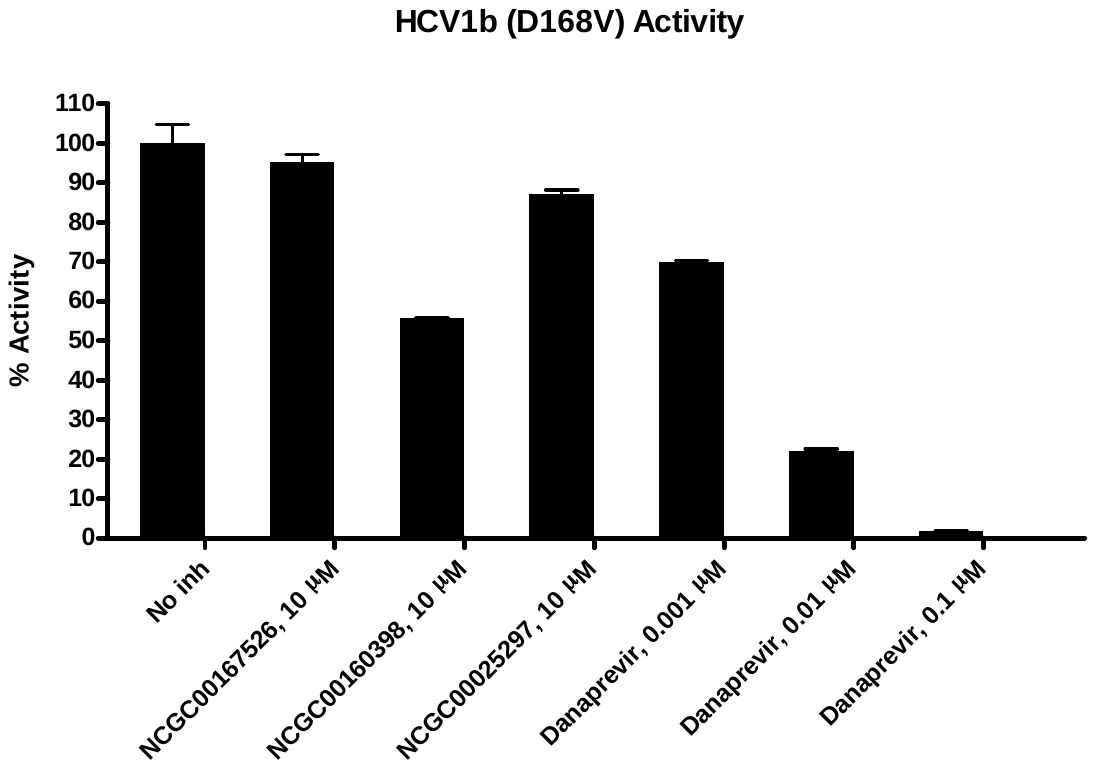

3.2.44. HCV1b (R155K)

| Compounds | HCV1b (R155K) Activity (Fluorescence count) | | Background  (Fluorescence count) | | % Activity | | % Inhibition |
| --- | --- | --- | --- | --- | --- | --- | --- |
|  | Repeat1 | Repeat2 | Repeat1 | Repeat2 | Repeat1 | Repeat2 |  |
| No Compound | 14521 | 14486 | 1245 | 1252 | 100 | 100 | 0 |
| NCGC00167526, 10 μM | 14539 | 14462 | 1432 | 1431 | 99 | 98 | 1 |
| NCGC00160398, 10 μM | 42200 | 41543 | 33942 | 34067 | 62 | 57 | 41 |
| NCGC00025297, 10 μM | 14743 | 15055 | 1353 | 1424 | 101 | 103 | 0 |
| Danoprevir, 0.01 μM | 12136 | 12244 | 1306 | 1231 | 82 | 83 | 18 |
| Danoprevir, 0.1 μM | 7275 | 6806 | 1254 | 1275 | 45 | 42 | 57 |
| Danoprevir, 1 μM | 2129 | 2062 | 1319 | 1376 | 6 | 5 | 94 |
| Background | 1248 | 1281 | 1243 | 1245 |  |  |  |

**Table 3.2.44. Data for the Effect of the Compounds on HCV1b (R155K) Activity**

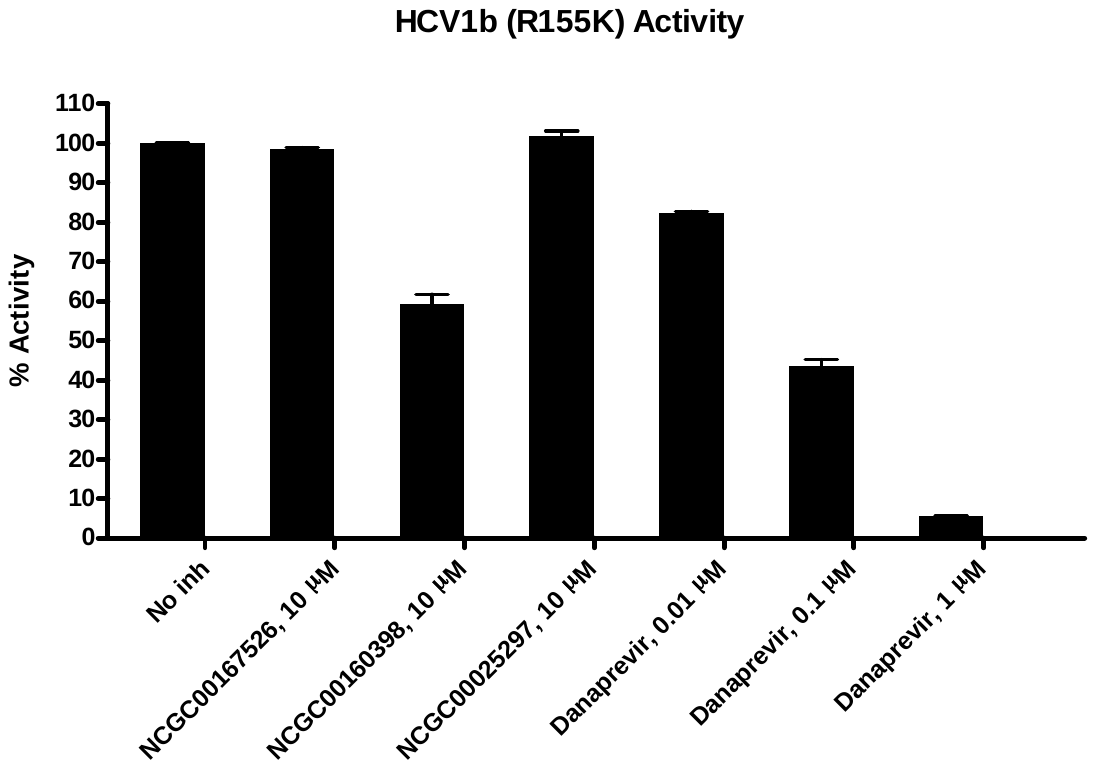

3.2.45. HCV1b (R155Q)

| Compounds | HCV1b (R155Q) Activity (Fluorescence count) | | Background  (Fluorescence count) | | % Activity | | % Inhibition |
| --- | --- | --- | --- | --- | --- | --- | --- |
|  | Repeat1 | Repeat2 | Repeat1 | Repeat2 | Repeat1 | Repeat2 |  |
| No Compound | 8468 | 8345 | 1245 | 1252 | 101 | 99 | 0 |
| NCGC00167526, 10 μM | 8433 | 8010 | 1432 | 1431 | 98 | 92 | 5 |
| NCGC00160398, 10 μM | 36677 | 37354 | 33942 | 34067 | 37 | 47 | 58 |
| NCGC00025297, 10 μM | 8003 | 7388 | 1353 | 1424 | 92 | 84 | 12 |
| Danoprevir, 0.01 μM | 6754 | 6904 | 1306 | 1231 | 77 | 79 | 22 |
| Danoprevir, 0.1 μM | 4774 | 4836 | 1254 | 1275 | 49 | 50 | 51 |
| Danoprevir, 1 μM | 1377 | 1452 | 1319 | 1376 | 0 | 1 | 99 |
| Background | 1264 | 1245 | 1243 | 1245 |  |  |  |

**Table 3.2.45. Data for the Effect of the Compounds on HCV1b (R155Q) Activity**

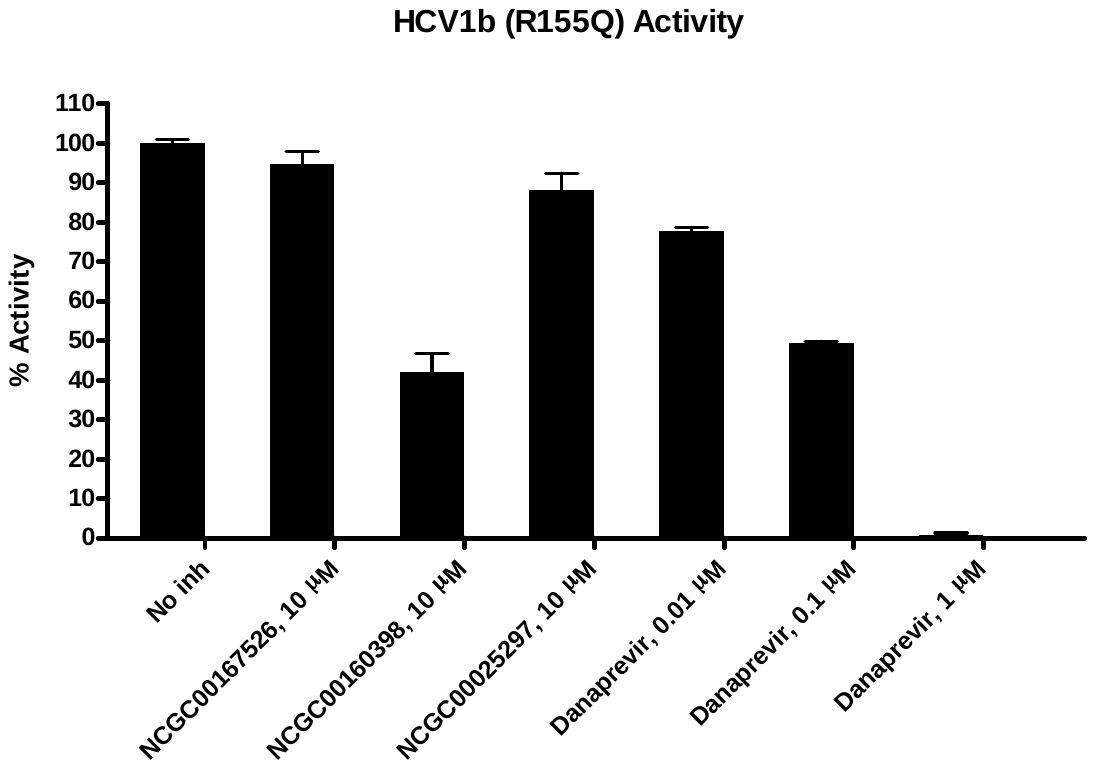

3.2.46. HCV2a

| Compounds | HCV2a Activity (Fluorescence count) | | Background  (Fluorescence count) | | % Activity | | % Inhibition |
| --- | --- | --- | --- | --- | --- | --- | --- |
|  | Repeat1 | Repeat2 | Repeat1 | Repeat2 | Repeat1 | Repeat2 |  |
| No Compound | 21181 | 20912 | 1245 | 1252 | 101 | 99 | 0 |
| NCGC00167526, 10 μM | 20492 | 21471 | 1432 | 1431 | 96 | 101 | 1 |
| NCGC00160398, 10 μM | 49497 | 50236 | 33942 | 34067 | 78 | 82 | 20 |
| NCGC00025297, 10 μM | 21575 | 20727 | 1353 | 1424 | 102 | 98 | 0 |
| Danoprevir, 0.001 μM | 19070 | 18665 | 1267 | 1274 | 90 | 88 | 11 |
| Danoprevir, 0.01 μM | 10037 | 10142 | 1306 | 1231 | 44 | 45 | 55 |
| Danoprevir, 0.1 μM | 1937 | 1773 | 1254 | 1275 | 3 | 2 | 97 |
| Background | 1259 | 1269 | 1243 | 1245 |  |  |  |

**Table 3.2.46. Data for the Effect of the Compounds on HCV2a Activity**

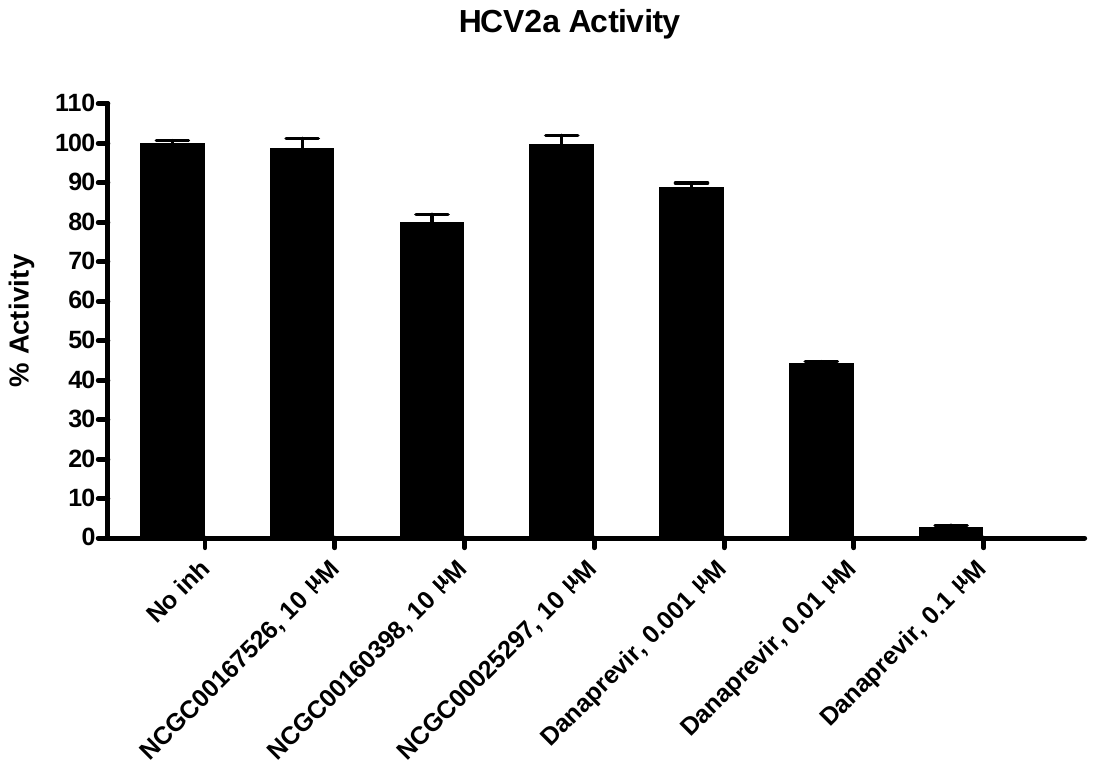

3.2.47. POP

| Compounds | POP Activity (Fluorescence count) | | Background  (Fluorescence count) | | % Activity | | % Inhibition |
| --- | --- | --- | --- | --- | --- | --- | --- |
|  | Repeat1 | Repeat2 | Repeat1 | Repeat2 | Repeat1 | Repeat2 |  |
| No Compound | 11881 | 12573 | 2048 | 2028 | 96 | 104 | 0 |
| NCGC00167526, 10 μM | 8714 | 8865 | 1697 | 1891 | 69 | 68 | 31 |
| NCGC00160398, 10 μM | 9486 | 9244 | 2251 | 2022 | 71 | 71 | 29 |
| NCGC00025297, 10 μM | 9558 | 9160 | 1939 | 1899 | 75 | 71 | 27 |
| Pro-Pro-CHO, 0.001 μM | 10899 | 11358 | 1660 | 1665 | 91 | 95 | 7 |
| Pro-Pro-CHO, 0.01 μM | 8688 | 9166 | 1622 | 1685 | 69 | 73 | 29 |
| Pro-Pro-CHO, 0.1 μM | 1156 | 1159 | 1163 | 1175 | 0 | 0 | 100 |
| Background | 1278 | 1233 | 1275 | 1190 |  |  |  |

**Table 3.2.47. Data for the Effect of the Compounds on POP Activity**

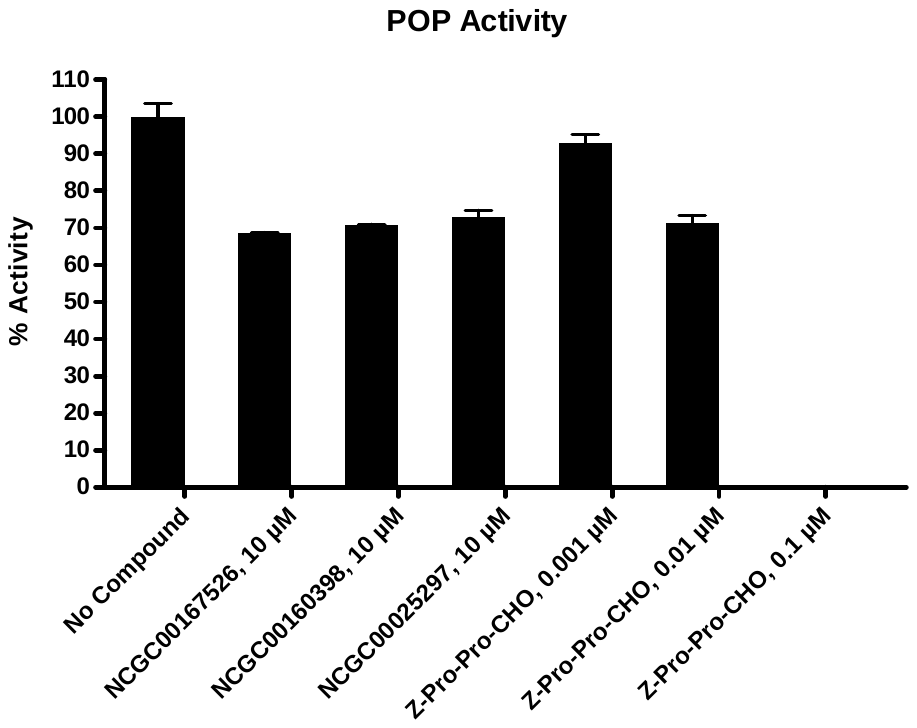

3.2.48. APC

| Compounds | APC Activity (Fluorescence count) | | Background  (Fluorescence count) | | % Activity | | % Inhibition |
| --- | --- | --- | --- | --- | --- | --- | --- |
|  | Repeat1 | Repeat2 | Repeat1 | Repeat2 | Repeat1 | Repeat2 |  |
| No Compound | 2659 | 2530 | 215 | 190 | 103 | 97 | 0 |
| NCGC00167526, 10 μM | 981 | 976 | 144 | 136 | 23 | 23 | 77 |
| NCGC00160398, 10 μM | 9268 | 9322 | 8853 | 8905 | 1 | 4 | 98 |
| NCGC00025297, 10 μM | 498 | 532 | 126 | 137 | 0 | 2 | 99 |
| Dabigatran, 0.2 μM | 2243 | 2216 | 136 | 175 | 85 | 84 | 16 |
| Dabigatran, 2 μM | 1676 | 1711 | 195 | 157 | 56 | 58 | 43 |
| Dabigatran, 20 μM | 615 | 598 | 133 | 131 | 6 | 5 | 95 |
| Background | 550 | 559 | 190 | 184 |  |  |  |

**Table 3.2.48. Data for the Effect of the Compounds on APC Activity**

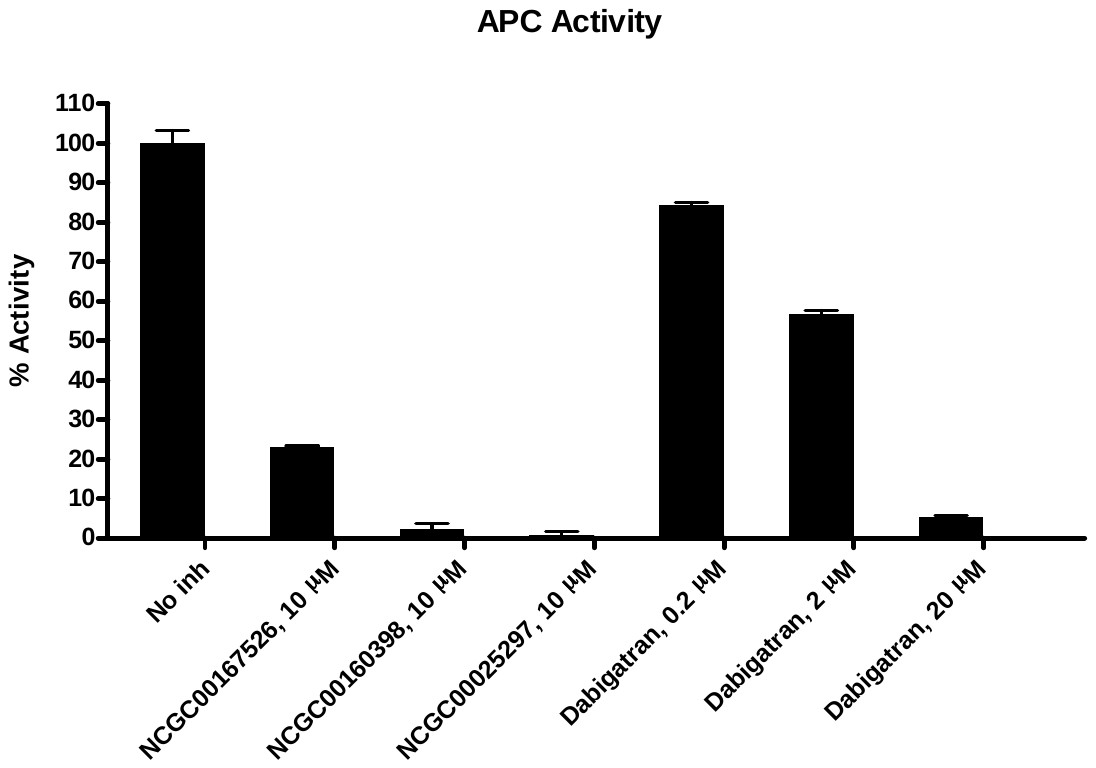

4. Quality Assurance Statement

I certify that the results presented in this report were generated using the materials and methods mentioned and that these results reflect the Raw Data.

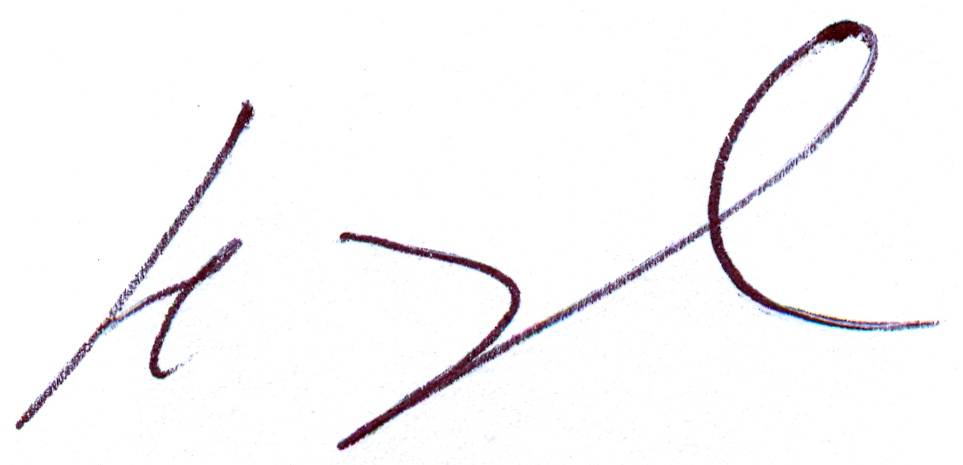

_____________________

Henry Zhu, Ph.D.

President

06-03-2020

______________________

Date
