## Supplementary Figures 1 - 3 for "An Enzymatic TMPRSS2 Assay for Assessment of Clinical Candidates and Discovery of Inhibitors as Potential Treatment of COVID-19"

Table of Contents:

Figure S1. Calculating $K_i^{app}$ for gabexate and FOY-251 using the Morrison Equation

Figure S2. Camostat, nafamostat and gabexate inhibition of recombinant human proteases from a BPS Bioscience protease panel

Figure S3. LC-MS chromatogram of bromhexine

**A**

$$v = v_0 \frac{K_i^{app}}{[I]_0 + K_i^{app}}$$

**B**

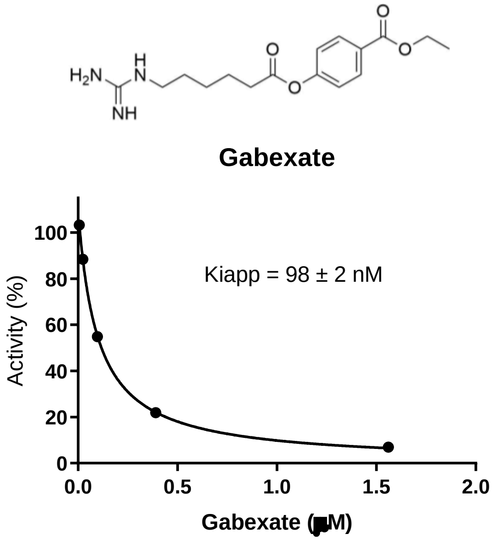


**C**

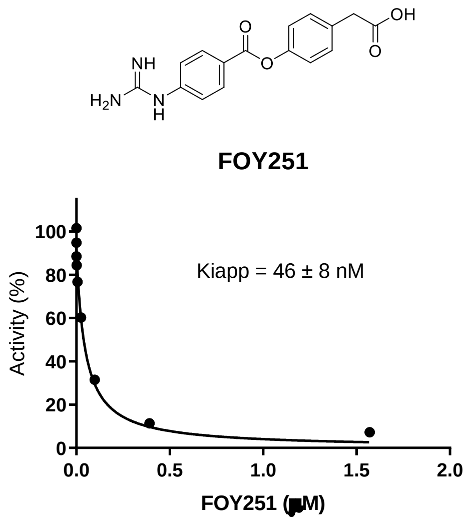


**Supplementary Figure 1**: **A)** Equation derived from the Morrison equation for when $K_i^{app}$ is relatively high compared to the active enzyme concentration. **B)** $K_i^{app}$ value for gabexate by fitting the dose-response data to the equation in A. **C)** $K_i^{app}$ value for FOY251 by fitting the dose-response data to the equation in A.

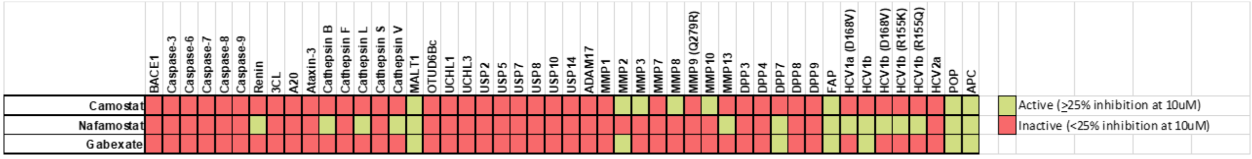


**Supplementary Figure 2**: Activity of camostat, nafamostat and gabexate against a panel of proteases. Compounds were tested at a single concentration (10μM) against all proteases, and activity data was conditionally formatted, where light green = inhibition (≥25% inhibition at 10μM) and red = inhibition (<25% inhibition at 10μM) (full report in Supplemental BPS Bioscience Report).

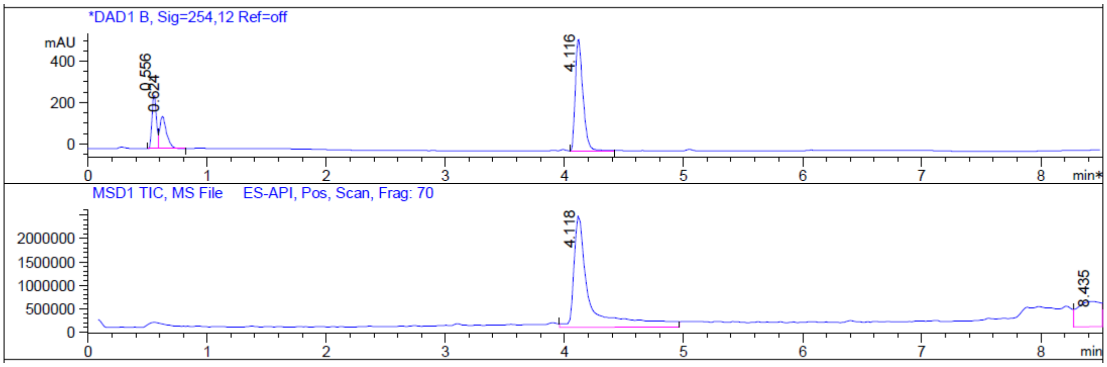


LC trace

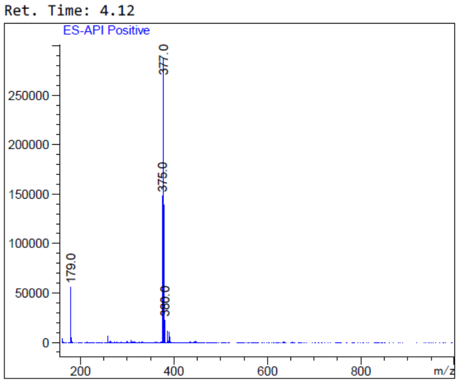


MS trace

**Supplementary Figure 3**: LC-MS chromatogram of bromhexine. Purity at 99.9%
